# Alignment-Integrated Reconstruction of Ancestral Sequences Improves Accuracy

**DOI:** 10.1101/2020.02.26.965186

**Authors:** Kelsey Aadland, Bryan Kolaczkowski

## Abstract

Ancestral sequence reconstruction (ASR) uses an alignment of extant protein sequences, a phylogeny describing the history of the protein family and a model of the molecular-evolutionary process to infer the sequences of ancient proteins, allowing researchers to directly investigate the impact of sequence evolution on protein structure and function. Like all statistical inferences, ASR can be sensitive to violations of its underlying assumptions. Previous studies have shown that, while phylogenetic uncertainty has only a very weak impact on ASR accuracy, uncertainty in the protein sequence alignment can more strongly affect inferred ancestral sequences. Here we show that errors in sequence alignment can produce errors in ASR across a range of realistic and simplified evolutionary scenarios. Importantly, sequence reconstruction errors can lead to errors in estimates of structural and functional properties of ancestral proteins, potentially undermining the reliability of analyses relying on ASR. We introduce an alignment-integrated ASR approach that combines information from many different sequence alignments. We show that integrating alignment uncertainty improves ASR accuracy and the accuracy of downstream structural and functional inferences, often performing as well as highly-accurate structure-guided alignment. Given the growing evidence that sequence alignment errors can impact the reliability of ASR studies, we recommend that future studies incorporate approaches to mitigate the impact of alignment uncertainty. Probabilistic modeling of insertion and deletion events has the potential to radically improve ASR accuracy when the model reflects the true underlying evolutionary history, but further studies are required to thoroughly evaluate the reliability of these approaches under realistic conditions.

## Introduction

Aside from happening upon a piece of preserved ancient DNA (Meyer et al. 2016) or reversing the arrow of time (Micadei et al. 2019), ancestral sequence reconstruction (ASR) is the only available technique for directly investigating the sequence, structure and function of ancient molecules. Because ASR studies rely on statistical inferences of ancestral sequences that cannot be validated directly, the accuracy with which ancestral protein sequences can be inferred has been a major concern of the ASR research community (Hall 2006; Randall et al. 2016; Eick et al. 2017). Previous studies have suggested that ASR is expected to be highly accurate in many cases (Randall et al. 2016; Vialle et al. 2018). Interestingly, studies have shown that the accuracy of the phylogenetic tree describing the evolutionary history of the protein family has only a very weak impact on ASR accuracy and generally only affects the most statistically-ambiguous sequence positions (Hanson-Smith et al. 2010). This largely counterintuitive result is due to the fact that the same evolutionary scenarios that make the phylogenetic tree uncertain also make ancestral sequences more similar across different phylogenies.

Some studies have suggested that there may be a trade-off between sequence reconstruction accuracy and the accuracy with which some structural and functional properties of the sequence can be inferred (Williams et al. 2006; Matsumoto et al. 2015; Arenas et al. 2017). Specifically, maximum-a-posteriori (MAP) ASR (also referred to as maximum-likelihood ASR), which reconstructs the most accurate protein sequences, can produce biased inferences of structural stability. This stability bias can be alleviated using a sampling approach that randomly generates ancestral sequences from the posterior probability distributions at each site. However, this sampling approach produces sequences that are less accurate than MAP reconstruction, which can impact inferences of other structural or functional properties (Eick et al. 2017).

One recent study found that the alignment of extant protein sequences forming the basis for phylogenetic inference and ASR can have a potentially strong affect on ancestral sequence reconstruction accuracy (Vialle et al. 2018). That ASR accuracy depends on alignment accuracy is concerning, as the ‘correct’ alignment of extant protein sequences can almost never be known with certainty, and there are few reliable methods for diagnosing alignment error or ambiguity (Dickson et al. 2010; Penn et al. 2010). It is currently unknown whether the same alignment errors that cause ancestral sequence reconstruction errors also impact the inferred structural or functional properties of reconstructed ancestral sequences, and no general methodologies exist to alleviate the impact of alignment error on ASR.

Here we develop and evaluate a novel ASR approach that combines information from many different sequence alignments to infer ‘alignment-integrated’ ancestral sequences. Although this approach does not completely eliminate the impact of alignment errors on ASR accuracy, we found that integrating sequence alignments reduces both ASR errors and errors in the structural and functional properties of inferred ancestral sequences, often performing as well as structure-guided sequence alignment. Our study suggests that, particularly for cases in which diverse structures of different protein family members are not available to guide the alignment process, integrating different alignments can be a reliable approach for mitigating the impact of alignment errors on ASR accuracy.

## Results and Discussion

### Alignment Errors Vary with Alignment Method and Protein Domain Family

To assess the impact of alignment errors on ancestral sequence reconstruction (ASR) accuracy, we used structural alignments of individual protein domains to simulate sequence data along empirical domain-family phylogenies (see Supplementary Table S1, Fig S1), with sequence composition and insertion-deletion (indel) patterns inferred from the structural alignment (see Methods). Simulated data were then aligned using a variety of sequence-based methods as well as a “structure-guided” approach that used the original structural alignment to ‘seed’ the alignment of additional sequences (see Methods). Comparing sequence-alignments and structure-guided alignments to the correct simulated alignment allowed us to evaluate the extent to which the simulation conditions generated alignment errors that could potentially impact ASR accuracy.

In general, both sequence-alignments and structure-guided alignments underestimated the correct alignment length by placing fewer gaps in the alignment, resulting in overestimation of the proportion of variable and parsimony-informative positions (Supplementary Table S2, Fig. S2). Across the 5 different protein-domain families used in this study, inferred alignments underestimated alignment length by 1.3-fold, on average (t-test *p*=6.41e^-4^), and the number of gaps by 1.2-fold (*p*=1.69e^-3^). The proportion of variable sites was overestimated by 1.8-fold (*p*=4.97e^-19^), and the proportion of parsimony-informative sites was overestimated by 1.9-fold (*p*=2.28e^-13^). Structure-guided alignments were no different from sequence-alignment methods in any of the calculated alignment attributes (t-test *p*>0.10), suggesting structure-guided and sequence-alignment methods tend to make similar errors in alignment length and the numbers of variable and parsimony-informative sites.

Although the general trend of alignment length underestimation is strongly supported by our data and is consistent with results from a previous study (Fletcher and Yang 2010), we observed significant variation in alignment errors, both across protein domain families and across alignment methods (Supplementary Table S2, Fig. S2). For example, clustalw tended to underestimate alignment length to a greater degree than other sequence-alignment methods (by 2.3-fold on average, vs 1.1-fold for other methods; t-test *p*<0.034). Across all alignment methods, the CARD protein domain family’s correct alignment length was underestimated to a greater degree (1.9-fold) than the other protein domain families (1.2-fold; t-test *p*<0.06). In contrast to this general trend of alignment length underestimation, mafft, probalign and tcoffee tended to overestimate the lengths of the correct DSRM1 and DSRM2 protein domain family alignments (by 1.3-fold; *p*<4.07e^-4^). These results suggest different alignment methods produce different types of alignment errors for different protein domain families. However, variation in alignment accuracy across replicate data sets of the same protein domain family was low, with the standard deviation never exceeding 11% of the inferred mean value of each alignment attribute (see Supplementary Table S2). This suggests that most of the variation in alignment accuracy is expected to be due to the particular interaction between a chosen alignment algorithm and the way a protein domain family has evolved, rather than stochastic variation in the simulated evolutionary process.

We quantified the distance of each inferred alignment from the correct simulated alignment using a position-wise distance metric, which estimates the probability that a randomly-selected residue from a randomly-selected sequence was aligned to an incorrect residue from another randomly-selected sequence (Blackburne and Whelan 2012). In general, the results of this distance-based alignment assessment (Supplementary Table S3, Fig. S3) were consistent with those of more traditional alignment metrics (Supplementary Table S2, Fig. S2). Across all protein domain families and alignment methods, the probability of randomly selecting an incorrectly-aligned residue was 0.31. However, there was strong variation in alignment distances across both domain families and alignment methods (Supplementary Table S3, Fig. S3). The CARD domain family produced larger average alignment distances than the other domain families (0.72 across alignment methods, vs 0.21 for the other domain families; t-test *p*<9.67e^-6^). Across protein domains, structure-guided alignments were >1.25-fold closer to the correct alignment than any of the sequence-alignment methods (t-test *p*<0.033). There were no detectable systematic differences in alignment distances among sequence-alignment methods, which produced average distances between 0.24 (msaprobs and probcons) and 0.43 (probalign; one-factor ANOVA *p*=0.92).

Overall, these results suggest the test cases used in this study cover a range of alignment difficulties that don’t strongly favor particular sequence-alignment algorithms over others and represent a reasonable test suite for assessing the impact of alignment errors on ancestral sequence reconstruction under realistic conditions. Structure-guided alignment methods have been shown to out-perform sequence-alignment in previous studies (Kim and Lee 2007), which has typically been attributed to the generally stronger conservation of protein structure vs sequence (Ingles-Prieto et al. 2013). Our alignment-distance results are consistent with these findings, but more traditional alignment metrics did not strongly differentiate structure-guided from sequence-alignment methods, suggesting the structure-guided approach might produce only marginally-better alignments under the challenging conditions used in this study.

### Alignment Errors Reduce Ancestral Sequence Reconstruction Accuracy

Each set of empirical simulation conditions (see Methods) was used to generate replicate correctly-aligned extant protein sequences at the tips of the phylogeny and correct ancestral sequences at every internal node. We assessed ancestral sequence reconstruction (ASR) error rates at each node on the phylogeny by comparing the ancestral sequence inferred using only an alignment of the extant protein sequences to the correct simulated ancestral sequence at that node. Three types of ASR errors were considered: 1) residue errors, in which both the inferred and correct ancestral sequences have an amino-acid residue at the given alignment position, but the residues differ; 2) insertion errors, in which the inferred ancestral sequence has a residue at the given alignment position, but the correct ancestral sequence has a gap character and 3) deletion errors, in which the inferred ancestral sequence has a gap character, but the correct ancestral sequence has an amino-acid residue. The ASR error rate for an inferred ancestral sequence was calculated as the number of ASR errors, divided by the length of the pairwise alignment of the correct to the inferred ancestral sequence (ie, errors/site). The expected ASR error rate for a given ancestral node was calculated as the mean error rate over replicates.

Given a sequence alignment, phylogenetic tree and statistical model of the molecular-evolutionary process, reconstruction of the most likely residue at each position in the alignment and node on the phylogeny has been well-described, as has the assessment of statistical confidence in the residue reconstruction (Yang et al. 1995; Koshi and Goldstein 1996). However, due to the way gap characters are encoded in most phylogenetic models, standard ASR does not reconstruct historical insertions or deletions (indels), resulting in ancestral sequences with no gap characters (Hall 2006). Unfortunately, the methodological details of ancestral indel reconstruction are poorly described in many published ASR studies (Chang et al. 2002; Gaucher et al. 2003; Bridgham et al. 2006; Voordeckers et al. 2012; Tan et al. 2016), making it difficult to assess how ancestral gaps were inferred. Although some methods have been developed that infer ancestral gaps as part of a more complex likelihood model (Redelings and Suchard 2005; Herman et al. 2014; Holmes 2017; Shim and Larget 2018), these approaches are largely untested and have not been adopted in many ASR studies. Many historical ASR studies probably used maximum-parsimony reconstruction of presence-absence ancestral states or a parsimony-like subjective criterion to place ancestral gap characters (Hall 2006; Hanson-Smith and Johnson 2016). Other studies have suggested using maximumlikelihood (ML, aka maximum a posteriori, MAP) reconstruction of presence-absence states to infer ancestral gaps (Ashkenazy et al. 2012), which is the approach we take here (see Methods).

Assuming the correct simulated alignment, we found that site-wise MAP reconstruction of ancestral gap states generated error rates comparable to residue reconstruction (Supplementary Tables S4-S8, Figs. S4-S7). Averaged across all protein domain families and ancestral nodes, ASR error rates were low when the correct alignment was known in advance (Supplementary Table S4), and the rate of residue-reconstruction errors (6.18e^-3^ errors/site) was slightly worse than the rate of erroneous insertions (8.49e^-4^ errors/site) or deletions (8.17e^-4^; t-test *p*<0.013). This pattern was generally observed across all the protein domain families (Supplementary Tables S4-S8, Figs. S4-S7). Only in the case of the CARD domain family was the residue reconstruction error rate (3.37e^-3^) slightly lower than the rate of erroneously-inferred insertions (3.55e^-3^) or deletions (3.44e^-3^), and these differences were not statistically significant (t-test *p*>0.31). For all other domain families, residue reconstruction error rates were slightly higher than indel reconstruction error rates (*p*<0.039). These results suggest site-wise MAP reconstruction of ancestral gap states—although failing to accurately model statistical dependencies among contiguous gaps (Ashkenazy et al. 2012)—provides a robust methodology for systematically inferring ancestral insertions and deletions under realistic conditions, provided the alignment is accurate.

When the alignment is not known in advance and must be inferred from the sequence data, we found errors in ancestral sequence reconstruction were higher overall and increased with increasing distance from the correct alignment (Fig. 1; Supplementary Tables S4-S8, Figs. S4-S7). Total ASR error rates were >4.8-fold higher when ancestral sequences were inferred using sequence-alignment methods (t-test *p*<5.78e^-3^). Even when structure-guided alignments were used to infer ancestral sequences, total ASR error rates were 4.4-fold higher than when the correct alignment was known in advance (t-test *p*<6.84e^-3^). These results were generally consistent across ASR error types and protein domain families (Supplementary Tables S4-S8, Figs. S4-S7). Residue errors increased by at least 2.6-fold when sequence-alignment was used, compared to the correct alignment (t-test *p*<0.013), and insertion-deletion errors increased by 12.3- and 12.7-fold, respectively (t-test *p*<4.37e^-3^). In all cases, structure-guided alignments produced ancestral sequences that had >1.5-fold fewer errors than sequence-alignment methods (t-test *p*<8.43e^-3^).

**Figure 1.**
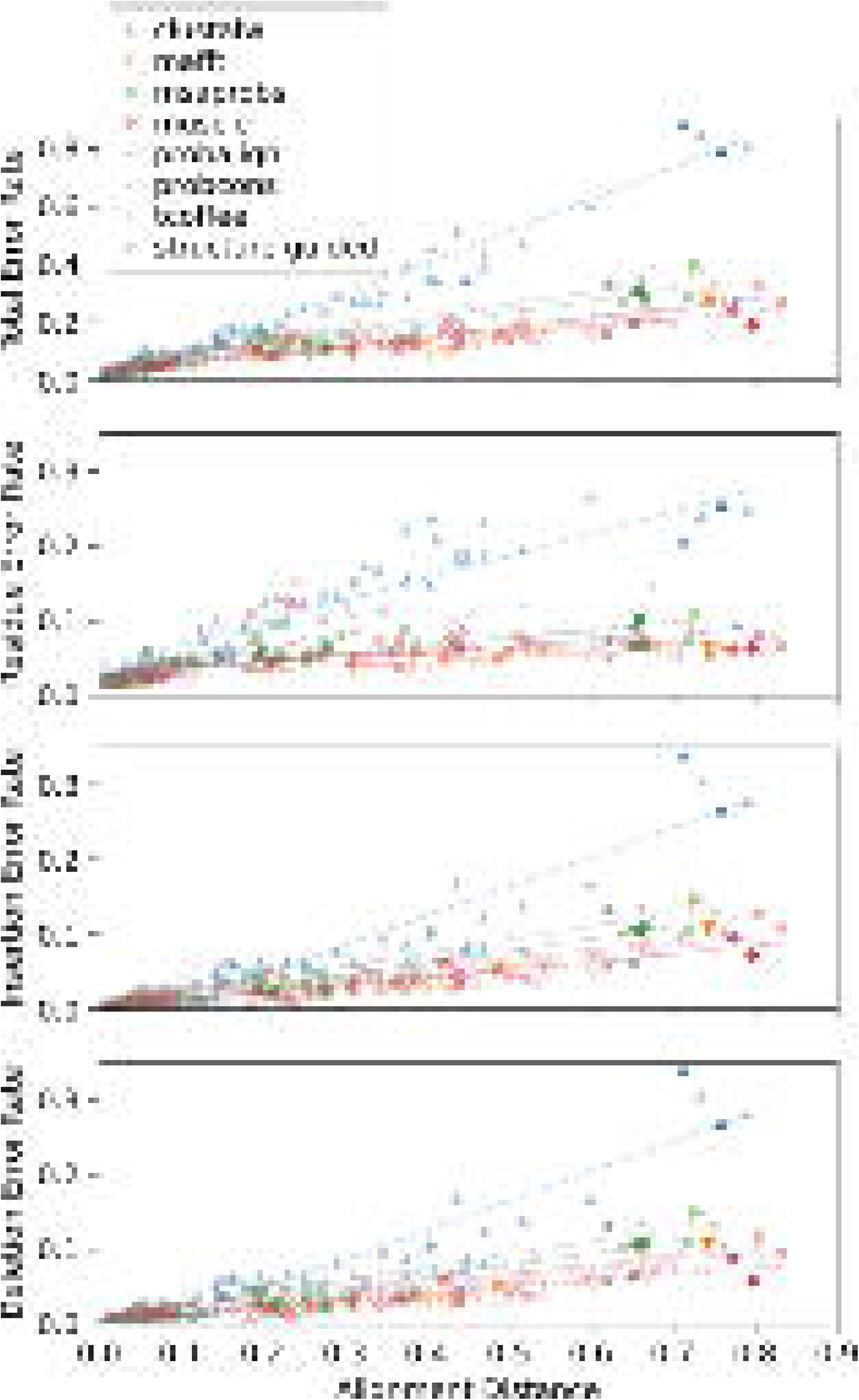
Errors in ancestral sequence reconstruction correlate with alignment errors. We simulated protein evolution using empirically-derived conditions from 5 protein domain families and aligned the resulting sequences using structure-guided and 7 different sequence-alignment methods. We measured the position-wise distance of each alignment from the correct simulated alignment (x-axis), which estimates the probability of selecting 2 incorrectly-aligned residues at random. We used each alignment to infer the most likely ancestral sequence at each node on the phylogeny and compared the inferred ancestral sequence to the correct simulated sequence to estimate ancestral sequence reconstruction (ASR) error rates. ASR errors were divided into 4 categories: 1) residue errors, in which both correct and inferred ancestral sequences have a residue at a given position in the alignment, but the residues differ; 2) insertion errors, in which the inferred sequence has a residue at a given alignment position, but the correct sequence has a gap; 3) deletion errors, in which the inferred sequence has a gap, but the correct sequence has a residue, and 4) total errors. Sequence-wide error rates (errors/site; y-axes) were computed by dividing the number of errors by the length of the pairwise alignment of the inferred and correct ancestral sequences. Dotted lines indicate least-squares linear regressions.

In general, there was a strong correlation between an inferred alignment’s distance from the correct alignment and total ASR error rate (Fig. 1; *r^2^*>0.78, mean *r^2^*=0.88). Similarly high correlations were observed for rates of insertion and deletion errors (*r^2^*>0.76, mean *r^2^*>0.85). However, residue errors were less strongly correlated with the distance from the correct alignment (mean *r^2^*=0.55), largely because muscle and probcons residue-errors were only very weakly correlated with alignment distance (*r^2^*<0.23). Interestingly, while error rate increased with alignment distance at roughly the same rate for alignment algorithms other than clustalw (slope of the best-fit regression line was 0.25-0.45 for total ASR error rate), clustalw’s total ASR error rate increased much more rapidly as the alignment diverged from the correct alignment (slope=1.08; ANCOVA *p*<1.63e^-3^). Clustalw’s ASR error rates were generally higher than the other alignment methods, even at comparable alignment distances (see Fig. 1), suggesting that, in addition to an alignment’s distance from the correct alignment, the specific types of alignment errors made can also affect how alignment errors impact ASR accuracy.

Our results confirm that ASR accuracy can be negatively impacted by alignment errors (Vialle et al. 2018) and suggest structure-guided alignment—although not a panacea—generally produces more accurate ancestral sequences than sequence-alignment methods.

### Alignment-Integrated Ancestral Sequence Reconstruction Improves Accuracy

Unlike the phylogeny (Hanson-Smith et al. 2010), our results and those of a previous study (Vialle et al. 2018) strongly suggest the sequence alignment—which can never be known with certainty in practice—can have a strong impact on ancestral sequence reconstruction. We hypothesized that integrating over alignment uncertainty could potentially alleviate this negative impact. To test this hypothesis, we developed a heuristic approach that reconstructs ancestral residues and gap states by integrating information from the 7 sequence-alignment methods examined in this study, placing equal prior weight on each alignment (see Methods).

We found that integrating information from many sequence-alignment algorithms reduced ASR error rates, compared to relying on any single sequence-alignment method (Fig. 2; Supplementary Tables S4-S8, Figs. S4-S7). On average, integrating over alignment uncertainty improved total ASR error rates by >1.3-fold, compared to choosing any single sequence-alignment method (t-test *p*<0.022). Although alignment-integrated ASR always generated fewer errors in residue and deletion reconstructions, improvements in these types of ASR errors were generally small and not always statistically significant. The fold-improvement in residue reconstruction error rates ranged from 1.1 (compared to tcoffee; t-test *p*=0.17) to 1.8 (clustalw; t-test *p*=0.012), while the improvement in deletion error rates ranged from 1.1-fold (probcons; *p*=0.18) to 2.1-fold (probalign; *p*=8.43e^-3^). The most dramatic reduction in ASR error rate was observed for insertion errors, for which alignment-integration improved error rates by >2.9-fold, compared to single sequence-alignment methods (t-test *p*<7.04e^-3^).

**Figure 2.**
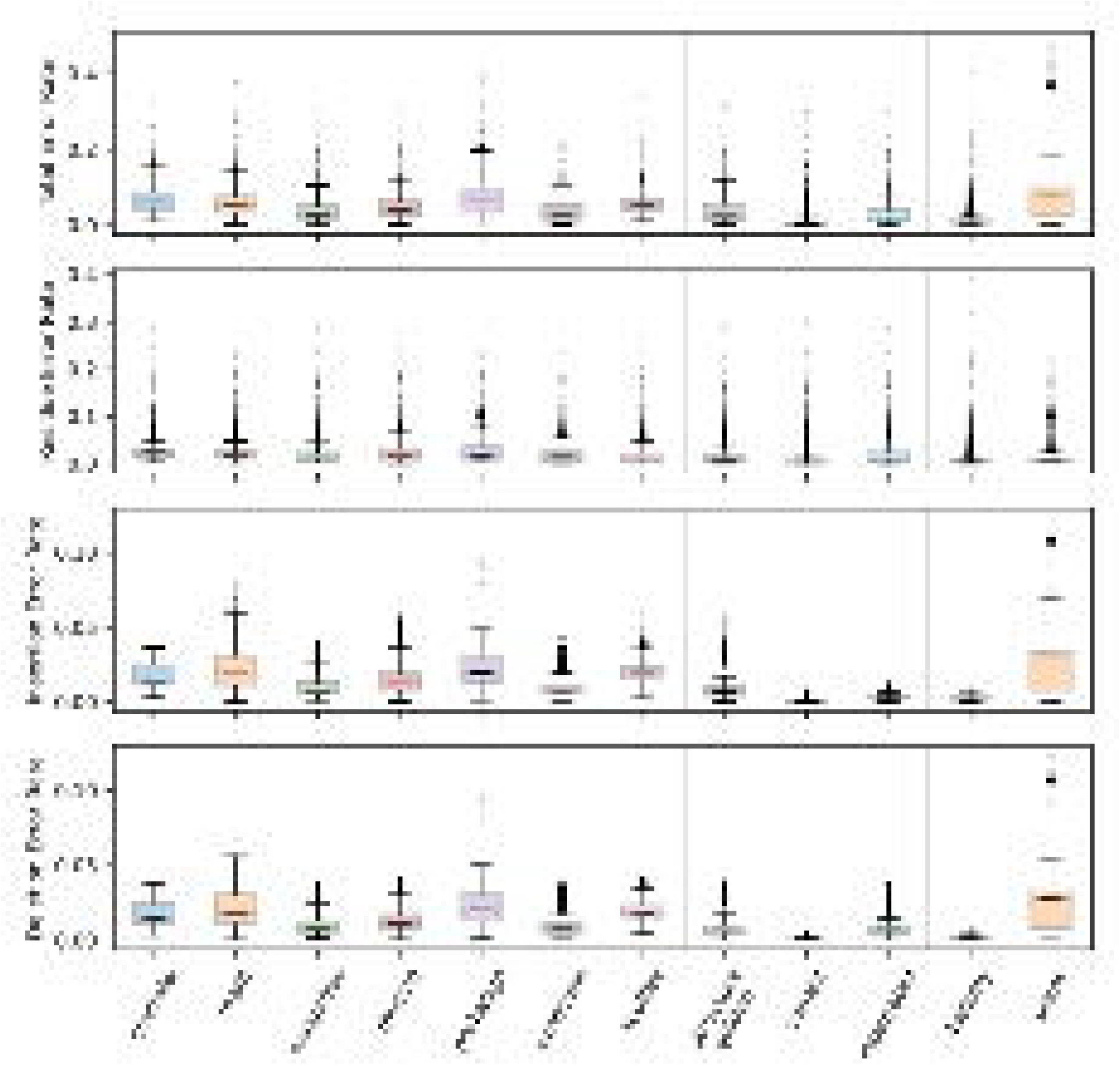
Alignment-integrated and structure-guided approaches produce fewer ancestral sequence reconstruction errors than single sequence-alignment methods. We simulated extant and ancestral sequences for 5 protein domain families, using empirically-derived conditions, and aligned the resulting extant sequence data using the correct simulated alignment, structure-guided and 7 different sequence-alignment methods. We used each alignment to infer the most likely ancestral sequence at each node on the phylogeny. Additionally, alignment-integrated ancestral sequences were generated by combining inferences from the 7 sequence-alignment methods. In each case, we compared the inferred ancestral sequence to the correct simulated ancestral sequence to estimate ancestral sequence reconstruction (ASR) error rates (expected errors/site). Error rate distributions were calculated over 10 replicate simulations and all nodes on each of the 5 protein domain family phylogenies. We plot the distributionsof total- (top), residue-, insertion- and deletion-errors (bottom) for each alignment method.

The same general pattern was consistently observed across all of the protein domain families examined in this study (Supplementary Tables S5-S8, Figs. S4-S7): compared to choosing a single sequence-alignment method, alignment-integration improved overall ASR error rates in all cases (by >1.1-fold), primarily by reducing the rate of insertion errors (by >1.8-fold; t-test *p*<0.022). In the case of the DSRM3 and RD domain families, the improvement in total ASR error rate was not always statistically significant, compared to some of the sequence-alignment methods. In both cases, mafft, msaprobs, muscle and probcons were statistically equivalent to alignment-integration (t-test *p*>0.057), and probalign was equivalent to alignment-integration for the RD domain family (t-test *p*=0.063).

These results suggest integrating over different sequence-alignment methods generally improves the accuracy of ancestral sequence reconstruction, compared to choosing a single sequence-alignment method, primarily by reducing the rate of erroneously-inferred insertions. The improvement in total ASR accuracy may be small for highly-conserved protein families or other scenarios in which sequence-based alignments are generally accurate.

Interestingly, integrating over many sequence-alignments slightly improved ASR error rates, even compared to the highly-accurate structure-guided alignment approach. Overall, total ASR error rates were reduced by 1.2-fold using alignment-integration, compared to structure-guided alignment (Fig. 2; t-test *p*=0.035), even though each of the individual sequence-alignments was farther away from the correct alignment than was the structure-guided alignment (see Supplementary Table S3, Fig. S3). Compared to structure-guided alignment, alignment-integration produced 1.2-fold more residue reconstruction errors (t-test *p*=0.059), the same number of deletion errors (t-test *p*=0.088) and 3.0-fold fewer insertion errors (t-test *p*=7.68e^-3^). This improvement in insertion error rate was observed for all domain families except the RD domain (*p*<0.033; see Supplementary Table S7, Fig. S6), and total ASR error rates were never significantly better using the structure-guided alignment (t-test *p*>0.11). These results suggest alignment-integration is a promising technique for reducing ASR error rates, even for protein families for which diverse structural data are not available to generate structure-guided alignments.

Protein sequence alignments are inferred using diverse methodologies, and new alignment methods are developed regularly. Most widely-used methods rely on heuristic strategies to place gap characters, with a rough ‘guide tree’ being used to order pairwise alignments (Chatzou et al. 2016). However, some methods have extended phylogenetic models to explicitly incorporate insertion-deletion (indel) events within a probabilistic framework (Redelings and Suchard 2005; Löytynoja 2014). We evaluated the accuracy of ancestral sequences reconstructed from two different ‘phylogenetically-aware’ probabilistic alignment methods: PRANK, which uses an indel model assuming a rough guide-tree approximation of the phylogeny (Löytynoja and Goldman 2008; Löytynoja 2014), and BAli-Phy, which uses Bayesian co-estimation of phylogeny and sequence alignment (Redelings and Suchard 2005). Because BAli-Phy is extremely computationally costly (Nute et al. 2019), analyses were conducted assuming the correct phylogenetic tree. To facilitate comparisons, PRANK and BAli-Phy were used only to generate sequence alignments, with ancestral sequences reconstructed using the same approach as for other alignment programs (see Methods).

Interestingly, we found that using the MAP alignment generated by BAli-Phy to reconstruct ancestral sequences was statistically indistinguishable from assuming the correct simulated alignment (see Fig. 2; t-test *p*=0.16), whereas PRANK alignments generated using a similar indel model (but without conditioning on the correct phylogenetic tree topology) resulted in among the least accurate ancestral sequences (Fig. 2). These results are consistent with those of a recent study examining protein sequence alignment accuracy, in which BAli-Phy generated highly-accurate sequence alignments from simulated data, whereas PRANK did not (Nute et al. 2019). while these results suggest probabilistic modeling of indel events could be a productive strategy for improving the accuracy of protein sequence alignment and ancestral reconstruction, future studies will be required to determine why similar approaches can produce very different results.

### Alignment-Integration Increases Ancestral Sequence Reconstruction Ambiguity

It is common practice in many ASR studies to reconstruct ‘plausible alternative’ states at positions with ambiguous reconstructions, in order to evaluate the impact of ASR uncertainty on downstream analyses (Eick et al. 2017). ASR errors that are only weakly supported are likely to be identified by this approach, whereas errors with very high posterior probability will likely be accepted as ‘correct,’ potentially undermining the validity of downstream structural or functional analyses.

We found that integrating many sequence-alignment methods, in addition to reducing ASR error rates, also reduced the posterior probabilities of erroneous ancestral states, when they were inferred (Fig. 3A; Supplementary Table S9). On average, the posterior probability of erroneously-inferred ancestral states was >0.9 when any single sequence-alignment or the structure-guided alignment was used to infer ancestral sequences. In contrast, the alignment-integrated approach had an average posterior probability of 0.67 for erroneous ancestral states, which was much more similar to that of the correct alignment (0.65; t-test *p*=0.066). Interestingly, this strong similarity between the alignment-integrated approach and the correct alignment was primarily confined to residue errors, for which alignment-integration produced an average posterior probability of 0.59, and the correct alignment’s mean posterior probability was 0.57 (*p*=0.093). In the case of insertion or deletion errors, the correct alignment produced low average posterior probabilities for erroneously-inferred ancestral states (<0.43), while alignment-integration’s posterior probabilities were much higher (>0.71, on average; t-test *p*<6.85e^-3^). Even in these cases, however, alignment-integration produced much more weakly-supported errors than any single sequence-alignment or structure-guided alignment, whose mean posterior probabilities for erroneously-inferred ancestral states were always >0.9 (t-test *p*<8.15e^-3^).

**Figure 3.**
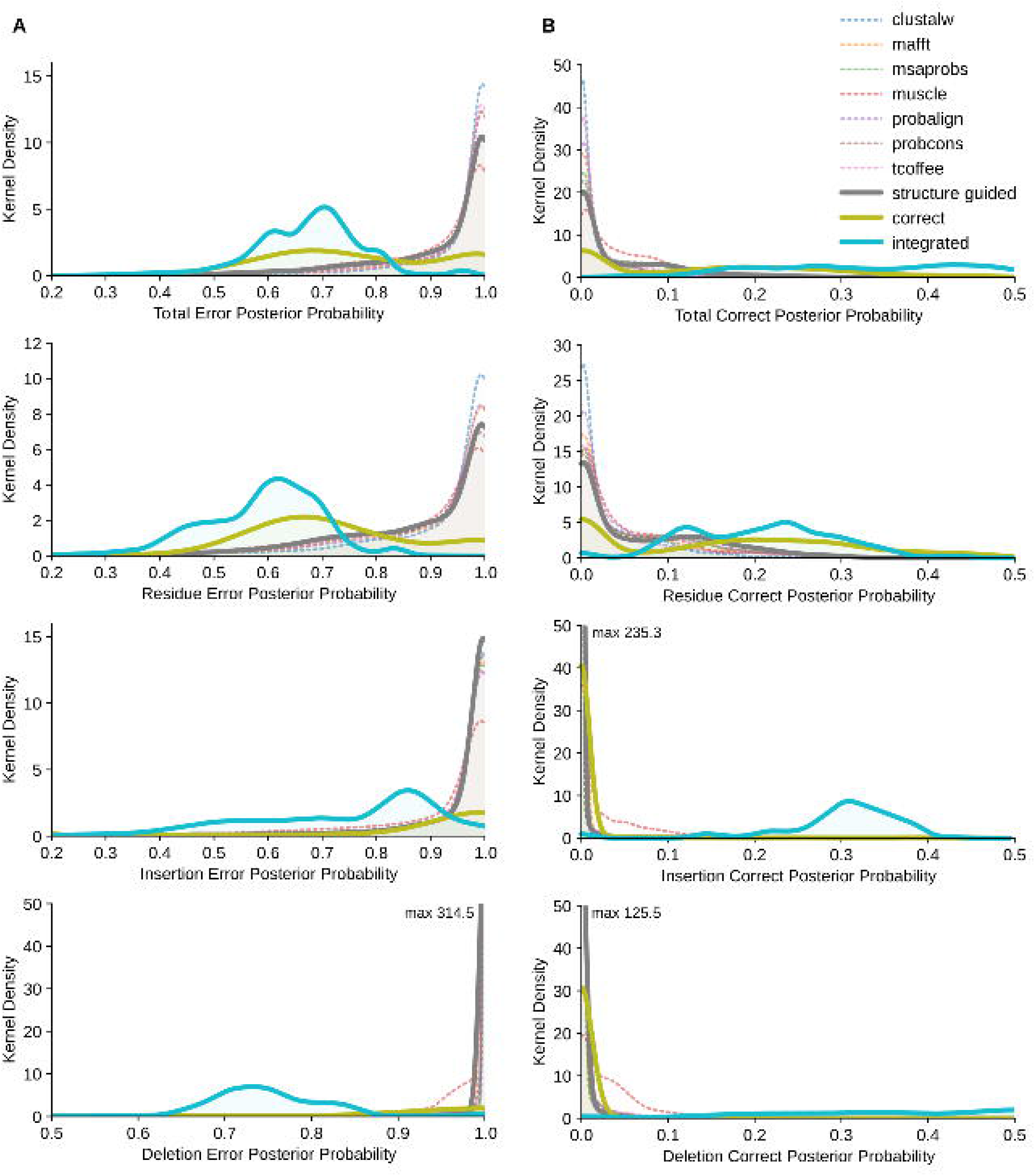
Alignment-integrated ancestral sequence reconstruction generates lower statistical confidence in erroneous ancestral states and stronger support for the correct state when errors are made. We used the correct simulated alignment, 7 different sequence-alignment methods, structure-guided alignment and alignment-integration to infer ancestral protein sequences from 5 empirically-derived simulation conditions. Total- (top), residue-, insertion- and deletion- (bottom) error rates (expected errors/site; x-axes) were calculated by comparing the inferred ancestral sequence to the correct simulated sequence at each node on the phylogeny. We used kernel density estimation to calculate the frequency distributions (y-axes) of posterior-probabilities for erroneous maximum a posteriori (MAP) ancestral states (left) and the correct ancestral state (right) when the correct state was not the inferred MAP state.

When ASR errors were made, alignment-integration also generated much stronger support for the correct ancestral state than any of the other alignment strategies, including the correct alignment (Fig. 3B; Supplementary Table S10). On average, the posterior probability of the correct ancestral state using sequence-alignment or structure-guided alignment was <0.046 when the correct state was not the MAP reconstruction. Assuming the correct alignment improved the mean posterior probability of the correct state by ~3-fold (to 0.14; t-test *p*<8.42e^-3^). However, alignment-integration further increased the mean posterior probability of the correct ancestral state by 2.4-fold, compared to the correct alignment (*p*=5.78e^-3^). Importantly, alignment-integration increased the posterior probability of the correct state to 0.33, on average, which is higher than the cutoff of 0.2-0.3 commonly used to identify plausible alternative ancestral reconstructions (Eick et al. 2017). This large increase in statistical support for the correct ancestral state when errors were made by alignment-integration was most pronounced for deletion errors (mean posterior probability 0.46) and less pronounced for insertion (mean PP=0.30) or residue errors (mean PP=0.21). For all types of errors, however, alignment integration produced >1.3-fold higher posterior probabilities for the correct ancestral state, compared to the correct alignment (t-test *p*<0.021) and >3.1-fold higher support than any other alignment method (*p*<4.78e^-3^).

While alignment-integration is obviously not a panacea, these results suggest, in addition to improving ASR accuracy, alignment-integration might be an important approach for ‘exposing’ some potential errors to downstream robustness analysis by reducing the statistical support for erroneously-inferred ancestral states and increasing the posterior probability of the correct ancestral state when it is not inferred as the MAP state.

The generally favorable increase in statistical ambiguity when ASR errors are made by alignment-integration does come at the cost of increased ambiguity for correctly-inferred ancestral states (Supplementary Table S11, Fig. S8). On average, correct ancestral state inferences were made with high statistical confidence using any of the methods examined in this study (>0.97 mean posterior probability). However, alignment-integration generated lower statistical confidence in correct ancestral state inferences than any of the other methods, all of which had >0.99 mean posterior probability for correctly-inferred states (t-test *p*<4.07e^-3^). All of the ASR methods exhibited stronger statistical support for correctly-inferred ancestral gap states vs correctly-inferred amino-acid residues (t-test *p*<0.011). However, this difference was more pronounced for alignment-integration, compared to the other ASR methods. When using alignment-integration, the mean posterior probability of correctly-inferred residues was 0.84 (>0.94 for the other methods), whereas the mean posterior probability was 0.98 for gap states correctly inferred by alignment-integrated ASR. These results suggest the reduced susceptibility to ASR error enjoyed by alignment-integration is also associated with increased ambiguity in reconstructed amino-acid residues, even when they are correctly inferred. This increased ASR ambiguity could potentially increase the operational costs of evaluating robustness to uncertainty as part of a typical ASR study.

### Alignment-Integration Improves Structural and Functional Inferences

In many ASR studies, the actual ancestral sequences are only of secondary interest, being commonly used to better understand how the protein’s structural and functional properties evolved (Eick et al. 2017). In some cases, researchers may decide to tolerate additional errors in sequence reconstruction, provided they result in more accurate inferences of specific structural or functional properties (Williams et al. 2006; Matsumoto et al. 2015; Arenas et al. 2017). To investigate the potential impact of alignment errors on the accuracy of downstream structural and functional investigations, we generated structural models of ancestral DSRM1 protein sequences inferred using each alignment method and estimated each protein’s structural stability and dsRNA-binding affinity using computational approaches (see Methods).

We found alignment-integrated ASR generally improved computational estimates of structural stability and RNA-binding affinity, compared to relying on a single sequence-alignment method (Fig. 4, Supplementary Table S12, Fig. S9). Aside from probcons, alignment-integration had significantly smaller errors in inferred structural stability than sequence-alignment methods (>1.14-fold; t-test *p*<0.048), performing similarly to structure-guided alignment (Fig. 4; t-test *p*=0.19). In this case, alignment-integration and probcons produced equivalent errors in structural stability estimates (t-test *p*=0.12). We observed similar results for estimated dsRNA-binding affinity (Supplementary Fig. S9). Alignment-integration produced >1.14-fold smaller errors in affinity estimates, compared to all sequence-alignment methods other than msaprobs (t-test *p*<0.041). Binding affinity errors were equivalent, on average, among alignment-integrated, structure-guided and msaprobs alignments (t-test *p*>0.055).

**Figure 4.**
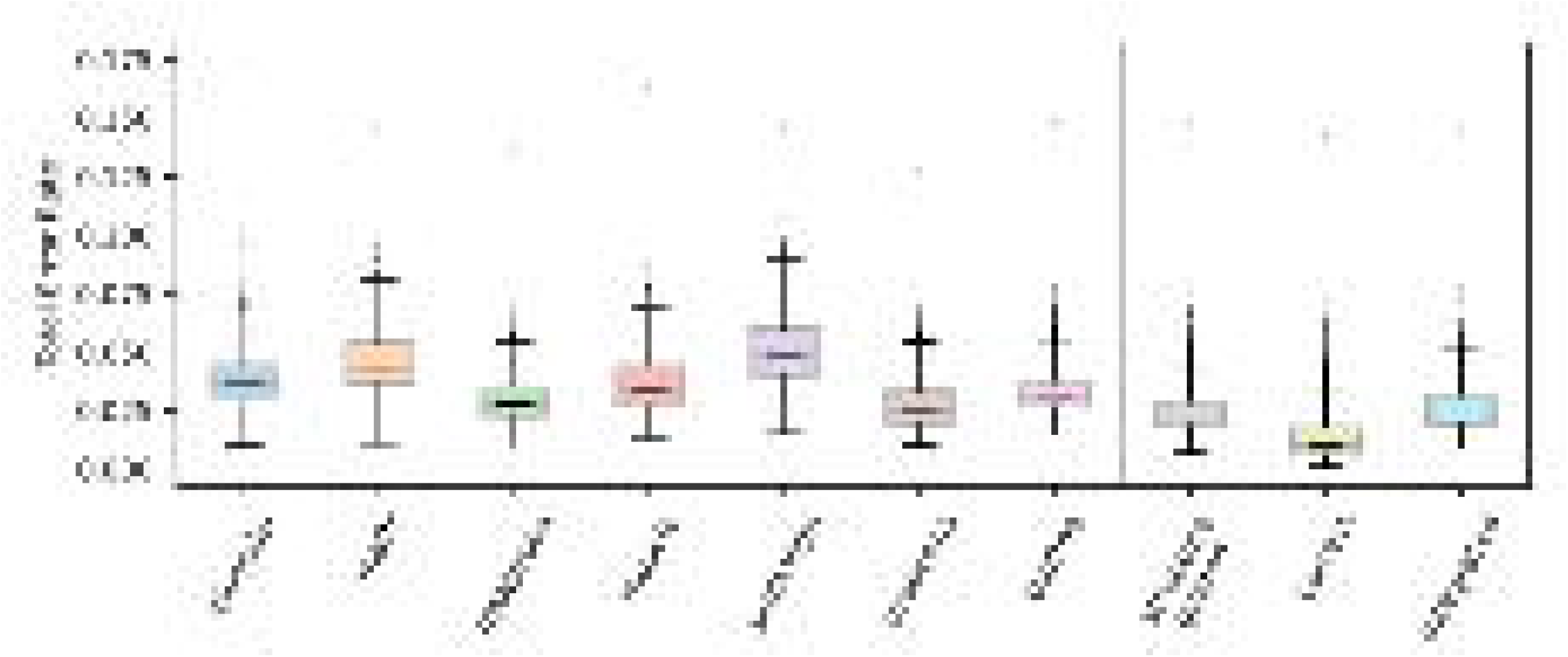
Alignment-integrated and structure-guided approaches produce less error in inferred structural properties of ancestral proteins than single sequence-alignment methods. We simulated replicate extant and ancestral sequences by evolving an RNA-binding protein domain along its empirically-determined phylogeny, using a structure-guided alignment to determine the amino-acid composition and pattern of insertions/deletions. Ancestral sequences were inferred using the correct simulated alignment, structure-guided alignment, 7 different sequence-alignment methods and alignment-integration. We modeled the structure of each ancestral sequence and estimated its structural stability (ΔG) using a computational approach. Errors in structural stability were calculated by comparing values estimated from the correct ancestral sequences to those estimated using each alignment method. Error rate distributions were calculated over 10 replicate simulations and all nodes on the phylogeny.

As expected, having the correct sequence alignment improved inferences of ancestral structural and functional properties in all cases (t-test *p*<0.017) but did not completely alleviate errors in structural stability or binding affinity estimates. On average, stability and affinity estimates deviated by >25% from the values inferred using the correct ancestral sequences (Supplementary Table S12, Fig. S10). The mean structural stability (ΔG) of correct ancestral DSRM1 domains was 0.087 cal/(mol x K) per residue (ie, the change in per-residue free energy of the native state, compared to misfolded or unfolded states, calculated using a contact-based energy model; see Methods), and structural stability estimates were typically 29.7% away from the correct values. Similarly, dsRNA-affinity estimates were, on average, 26.7% away from the values inferred using the correct ancestral sequences.

Together, these results suggest ambiguity or bias in the ancestral sequence reconstruction process can itself contribute to errors in downstream structural and functional inferences under challenging conditions (Williams et al. 2006; Arenas et al. 2017). Alignment errors appear to exacerbate errors in estimated structural stability and binding affinity of ancestral proteins, but structure-guided alignment or alignment-integration significantly reduced these errors.

There was a weak but significant positive correlation between ancestral sequence reconstruction errors and errors in structural and functional estimates for all alignment methods (Supplementary Table S13, Fig. S11). Errors in the inference of the ancestral sequence explained 40% of the variation in structural stability error (*r^2^*<0.78) and 29% of the variation in binding affinity error (*r^2^*<0.64). The mean slope of the best-fit regression line across all alignment methods was 0.44 for structural stability and 4.25 for binding affinity, and all slopes were significantly greater than zero (t-test *p*<2.50e^-3^). There were some differences in both correlation and slope across alignment methods. For example, clustalw, mafft, probalign and tcoffee showed weaker correlations between ASR error rates and structural stability errors (*r^2^*<0.29), while the remaining alignments had generally higher correlations (*r^2^*>0.41). Similarly, the slope of the best-fit regression line varied from a minimum of 0.17 (probalign) to a maximum of 0.74 (for the correct alignment). Similar results were observed for the correlation between ASR error rate and binding affinity errors: correlation varied between *r^2^*=0.021 (clustalw) and *r^2^*=0.64 (correct alignment), and slope varied from 1.07 (clustalw) to 7.00 (probalign; Supplementary Table S12). Qualitatively similar results were observed when considering different types of ASR sequence errors (see Supplementary Table S13, Fig. S11).

These results suggest the overall rate of sequence reconstruction errors is positively correlated with errors in estimates of structural and functional properties of ancestral sequences. The generally lower magnitudes of structural and functional errors observed for the structure-guided and alignment-integrated methods can be at least partially explained by their generally lower sequence-error rates. However, precisely how sequence-reconstruction errors translate into errors in structural or functional estimates is expected to be complex in realistic cases, and the specific types of sequence-reconstruction errors made by different alignment algorithms likely also plays a role in determining errors in structural or functional estimates.

### Integrating Posterior Probability Distributions Improves ASR Accuracy

To begin systematically investigating the factors impacting ASR accuracy, we varied the branch lengths and insertion-deletion (indel) rates along a minimal 3-taxon phylogeny with equal branch lengths and the same indel rate on all branches (Supplementary Fig. S12). Sequences were simulated using the JTT+G evolutionary model, and indels were placed randomly along the sequence. The ancestral sequence at the only node on the phylogeny was reconstructed using the correct simulated alignment, 7 sequence-alignment methods and the alignment-integrated approach (see Methods).

Results from this 3-taxon simulation were largely consistent with those obtained using larger, more realistic phylogenies (Fig. 5; Supplementary Figs. S13-S16). Across all simulation conditions, alignment-integration slightly improved ASR accuracy by 1.06-fold, compared to choosing a single sequence-alignment strategy at random (t-test *p*=4.01e^-9^). Even when the optimal sequence-alignment strategy was chosen for each set of simulation conditions, alignment-integration improved ASR accuracy by 1.03-fold (t-test *p*=3.33e^-4^). Although the improvement in alignment-integrated ASR accuracy was typically small in this case (~1%, on average), it was consistent across the vast majority of simulation conditions. Only under 5/64 conditions was the best sequence-alignment method as accurate or more accurate than alignment-integration, and most of these conditions had short branch lengths and low indel rates, leading to very low ASR errors across all methods.

**Figure 5.**
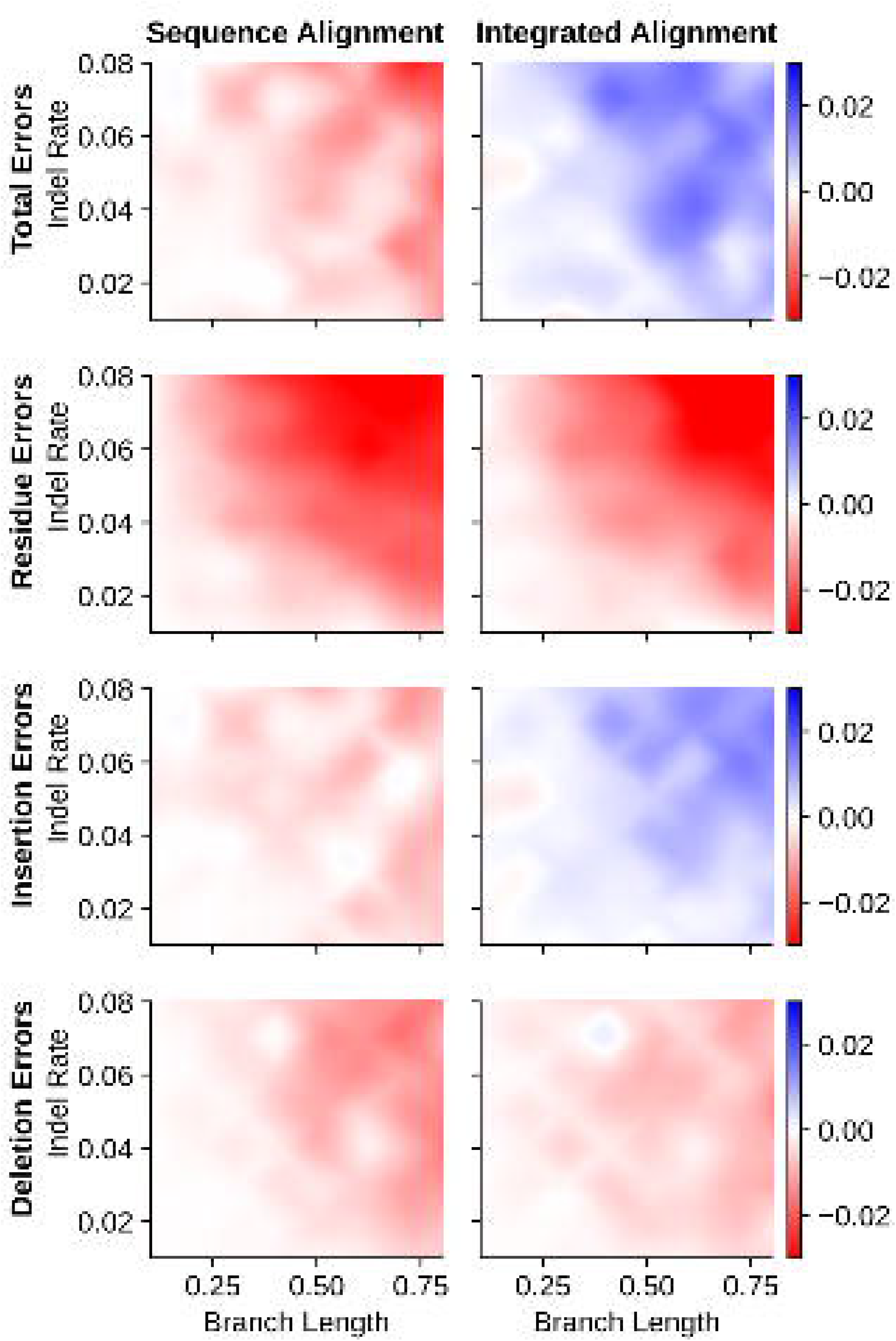
Alignment-integrated ancestral sequence reconstruction produces low rates of reconstruction errors in simplified 3-taxon simulations. We simulated protein sequences along a 3-taxon phylogeny with equal branch lengths (x-axes) and the same insertion-deletion rate (y-axes) across the phylogeny (see Supplementary Fig. S12). The most likely ancestral sequence at the single node on the phylogeny was inferred using the correct simulated alignment, 7 different sequence-alignment methods and alignment-integration. Total- (top), residue-, insertion- and deletion- (bottom) error rates were calculated by comparing inferred ancestral sequences to the correct simulated ancestral sequence. For each set of simulation conditions, we calculated the difference in error rates between the correct alignment vs the least-erroneous sequence-alignment method (left column) or vs alignment-integration (right column). Positive values (blue) indicate that the correct alignment produced more errors than the given inference method, and negative values (red) indicate that the correct alignment produced fewer errors.

As expected, ASR error rates increased with increasing branch lengths for all methods (linear regression slope >0.38, *r^2^*>0.85, t-test *p*<1.09e^-3^; Supplementary Table S14, Figs. S13-S16). For short branch lengths, ASR error rates were weakly correlated with increasing indel rates: when branches were <0.6 substitutions/site, linear regression slope was >0.19 (*r^2^*>0.68, t-test *p*<0.012; Supplementary Table S14, Figs. S13-S16). However, the correlation between ASR error and indel rates was not observed for branch lengths >0.6 (t-test *p*>0.044).

Interestingly, alignment-integration appeared slightly more accurate than using the correct sequence alignment under some conditions, improving ASR error rates by ~0.6%, on average, compared to the correct alignment (Fig. 5); however, this difference was not statistically significant (t-test *p*=0.42). We did observe slightly lower insertion error rates using alignment-integration, compared to the correct alignment (t-test *p*=9.56e^-3^). Residue errors were more frequent for alignment-integration (t-test *p*=4.85e^-8^), and there was no overall difference in deletion error rates between the two methods (t-test *p*=0.25).

These results suggest alignment-integration can consistently improve ASR error rates, compared to single sequence-alignment methods, even under an extremely simplified 3-taxon model system with random indels. Under some of these simplified conditions, alignment-integration can produce error rates comparable to those obtained when the correct alignment is known in advance.

Results from 3-taxon simulations suggest that simple ‘majority-rule’ is sufficient to explain most of the cases in which alignment-integration improves ASR accuracy (Fig. 6). On average, when one of the alignment methods produced an error that was not present in the alignment-integrated ancestral sequence, 66.3% of the other sequence-alignment methods reconstructed the correct ancestral state. This result was consistent across all simulation conditions (standard error 0.002; see Fig. 6). Interestingly, residue-reconstruction errors tended to have a weaker majority in favor of the correct ancestral residue; on average only 60.0% of other alignments recovered the correct ancestral residue when one alignment made a residue-reconstruction error (z-test *p*=9.17e^-32^). Insertion errors had the strongest majority in favor of the correct ancestral state, with, 82.5% of alternative alignments recovering the correct gap state when one alignment erroneously inferred an insertion at that position (z-test *p*<1.75e^-48^).

**Figure 6.**
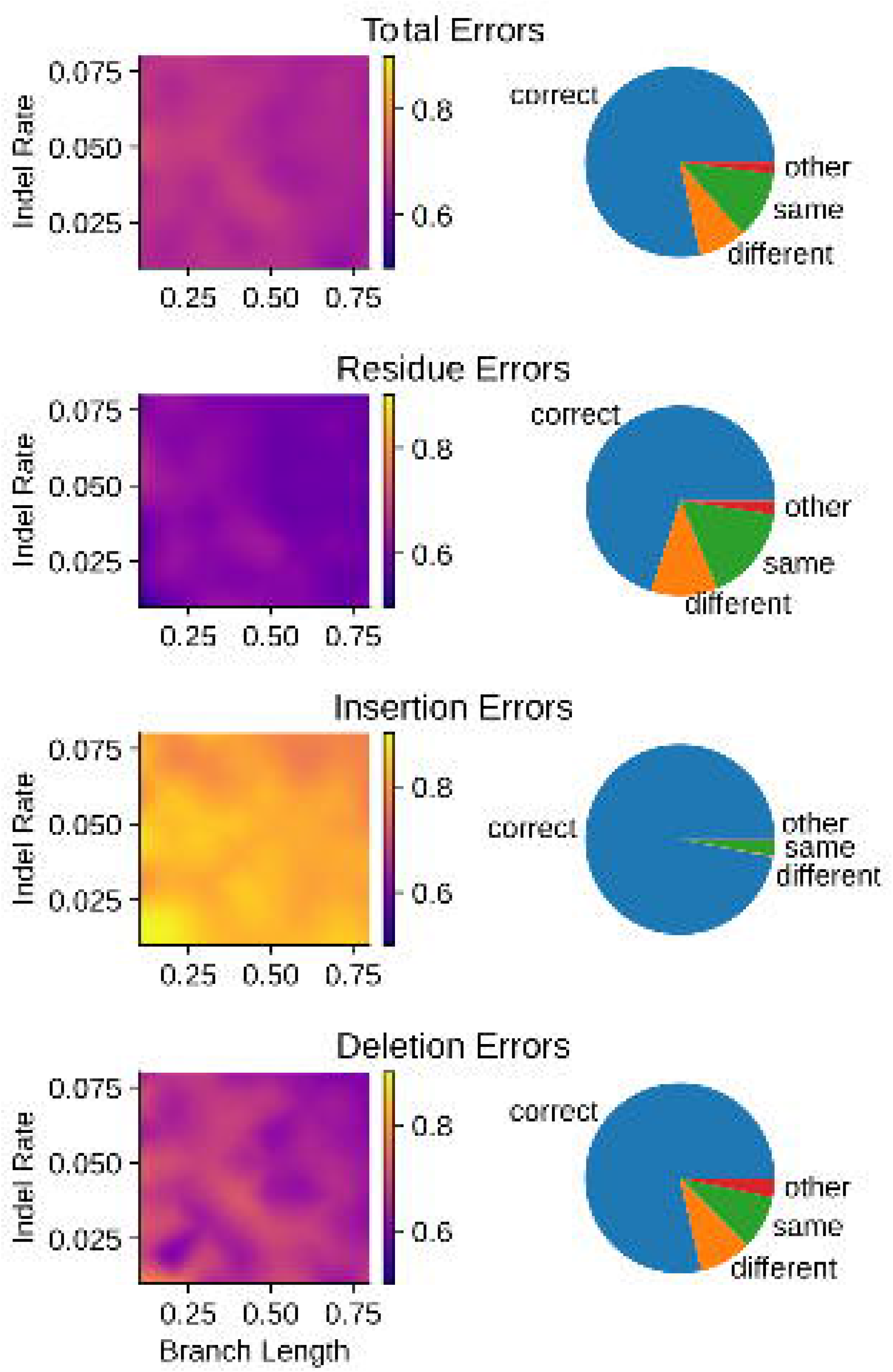
‘Majority-rule’ explains the majority of cases in which alignment-integration improves ancestral sequence reconstruction error rates. We simulated protein sequences along a simplified 3-taxon phylogeny, varying the branch length and insertion-deletion (indel) rate (see Supplementary Fig. S12). Ancestral sequences were inferred using 7 different sequence-alignment methods and alignment-integration. Total- (top), residue-, insertion- and deletion- (bottom) errors were calculated by comparing inferred ancestral sequences to the correct simulated ancestral sequence. Here, we consider only those cases in which a single sequence-alignment method produces an ASR error that is ‘repaired’ by alignment-integration. **Left panel:** For each branch length (x-axis) and indel rate (y-axis), we plot the proportion of alternative sequence-alignment methods that inferred the correct ancestral state when one sequence-alignment method generated an ASR error that was not found in the alignment-integrated ancestral sequence. **Right panel:** Across all simulation conditions, we consider all cases in which a sequence-alignment method makes an error, and that error is repaired by alignment-integration. We report the proportion of such cases in which 1) the majority of alternative sequence-alignment methods infer the correct ancestral state (“correct”; blue), 2) the majority of alternative sequence-alignment methods infer an incorrect ancestral state, but that state is different from the original error (“different”; orange), 3) the majority of alternative sequence-alignment methods infer the same incorrect ancestral state (“same”; green), and 4) other scenarios (“other”; red).

When alignment-integration was able to ‘repair’ an ASR error made by a single sequence-alignment method, 78.4% of these repairs were explainable by majority-rule (Fig. 6). However, in 8.1% of cases, the majority of alternative alignments also produced ASR errors, but the errors differed across alignments. Interestingly, in 11.5% of cases, the majority of sequence-alignments actually produced the same ASR error, even though alignment-integration reconstructed the correct ancestral state. Although the specific proportions differed somewhat across different types of ASR errors (see Fig. 6), the pattern of ASR error repairs due to alignment-integration was consistent: most repairable errors (70.0-97.0%, depending on error type) could be attributed to majority-rule, with smaller proportions of errors being repaired by alignment-integration when most sequence-alignments generate different ASR errors (0.2-11.4%) or when most alignments make the same ASR error (2.6-16.3%).

For cases in which majority-rule could not explain alignment-integration repair of the ASR error, we observed an upward shift in the posterior probability of the correct ancestral state, compared to similar scenarios that were not repaired by alignment-integration (Fig. 7). The proportion of cases in which the correct ancestral state had posterior probability <0.1 fell from 0.65 when alignment-integration did not repair the ASR error to 0.51 when alignment-integration repaired the ASR error, even though the majority of sequence-alignments produced an erroneous ancestral state (z-test *p*<1.0e^-20^). Similarly, the proportion of correct ancestral states with posterior probability <0.05 fell from 0.48 when alignment-integration did not repair the error to 0.21 when it did (z-test *p*<1.0e^-20^).

**Figure 7.**
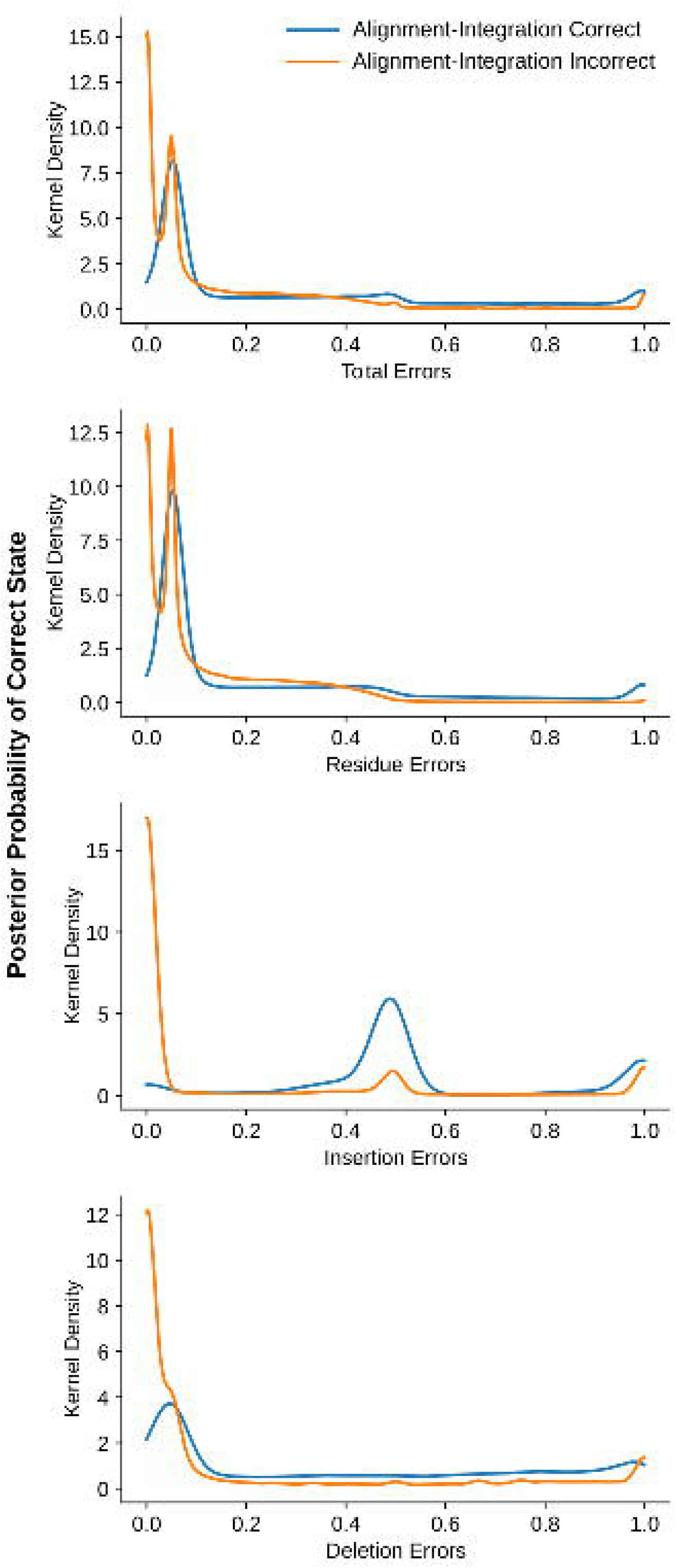
Alignment-integration can ‘repair’ ancestral sequence reconstruction errors when the correct ancestral state is not strongly-disfavored across sequence-alignment methods. We simulated protein sequences along a 3-taxon phylogeny, varying the branch length and insertion-deletion (indel) rate (see Supplementary Fig. S12). Ancestral sequences were inferred using 7 sequence-alignment methods and alignment-integration. Total- (top), residue-, insertion- and deletion- (bottom) errors were calculated by comparing inferred ancestral sequences to the correct ancestral sequence. Here, we consider only those cases in which a single sequence-alignment method produces an ASR error, and the majority of alternative sequence-alignment methods do not infer the correct ancestral state. We further divide such cases into 1) errors that are ‘repaired’ by alignment-integration (blue) and 2) errors that are not repaired by alignment-integration (orange). Under each scenario, we estimate the frequency distribution of the posterior probability of the correct ancestral state by kernel density estimation.

When alignment-integration was able to repair an ASR error by mechanisms other than majority-rule, we observed a large peak in frequency at the posterior probability making the ancestral reconstruction maximally-ambiguous (Fig. 7), which occurs at 0.05 for residue- and deletion-errors (for which the correct ancestral state is one of the 20 amino-acid residues) and at 0.5 for insertion-errors (for which the correct ancestral state is the gap state). This suggests that a relatively large proportion of ASR errors which majority-rule fails to repair—but which alignment-integration still repairs by other mechanisms—occur at highly-ambiguous positions with relatively flat posterior-probability distributions across many possible ancestral states. Interestingly, the peak at 0.5 posterior probability for insertion-errors was particularly pronounced when alignment-integration repaired the ASR error (Fig. 7), suggesting ‘balance-of-probability’ repairs may be particularly efficacious in cases of insertion errors, which may contribute to alignment-integration’s very low insertion error rates (see Fig. 2).

Overall, these results suggest majority-rule accounts for ~80% of cases in which alignment-integration is able to ‘repair’ an ancestral reconstruction error generated by a single sequence-alignment method, but more subtle effects of integrating posterior probability distributions also contribute to improved ASR accuracy by alignment-integration. When the correct ancestral state does not have very low posterior probability across all sequence alignments, alignment-integration can sometimes repair ancestral sequence reconstruction errors, even when the majority of sequence alignments reconstruct the wrong ancestral state.

## Conclusions

For future ASR studies, our results add to the emerging evidence that alignment errors cannot always be ignored when evaluating the accuracy of ancestral sequence reconstruction (Vialle et al. 2018), and in practice, sequence-alignment methods cannot always be relied upon to generate alignments accurate enough to ensure reliable ASR. When multiple structures from across the protein family are available, our results suggest that structure-guided alignment is an efficient approach for improving ASR accuracy, but many protein families lack the rich empirical structural data necessary for structure-guided alignment. In these cases, we recommend that future studies make some effort to evaluate the impact of alignment ambiguity on ASR results. The alignment-integration approach we present here is one mechanism for incorporating alignment ambiguity into ASR studies, which so-far appears to perform well across a variety of realistic and simplified model problems.

The empirically-derived simulation conditions used in this study represent realistic but highly-challenging ASR problems, with generally long branches and large phylogenies (see Supplementary Fig. S1), which are expected to contribute to elevated alignment and ASR error rates (Vialle et al. 2018). Under less challenging conditions, standard ASR methodology has been found to be highly reliable (Hanson-Smith et al. 2010; Randall et al. 2016; Vialle et al. 2018), and alignment-integration is unlikely to provide any benefits under conditions in which most sequence-alignment methods are extremely accurate.

Alignment-integration is computationally costly, as many different alignments need to be inferred, and phylogenetic model parameters and ancestral sequences need to be computed using each alignment before being combined. Alignments that are very similar to one another may be at best redundant and at worst could bias the ancestral sequence reconstruction toward a ‘false consensus.’ Similarly, wildly-inaccurate alignments could introduce statistical noise or generate biased results when included in the integration process. The identification of a set of alignment algorithms that tend to produce highly accurate but different alignments is expected to be important for reducing the computational demands of alignment-integrated ASR while maintaining its useful statistical properties and low error rates.

In many of our analyses, specific sequence-alignment algorithms are able to generate ancestral sequences that are nearly as accurate as those generated by alignment integration or structure-guided alignment, suggesting that using a single sequence-alignment method may be adequate, even in some challenging cases. However, the specific alignment algorithm that will perform well for a specific ASR study may be difficult to determine in practice and would likely require comparisons with other alignment algorithms, which would be nearly as computationally costly as alignment-integrated ASR. Because there is no known alignment method that performs optimally in all cases, we recommend that, at minimum, future ASR studies that rely on a single sequence alignment strongly justify the specific approach chosen as the most appropriate for the study.

Our cursory analysis of probabilistic alignment methods suggests Bayesian co-estimation of the alignment and phylogenetic tree has the potential to provide exceptionally accurate ancestral sequence reconstructions, at least under some conditions. However, we remain cautious about recommending this approach for a number of reasons. First, our analyses were conditioned on the correct phylogenetic tree, which is almost never known with certainty, and the accuracy of ASR under the more realistic case of joint alignment-phylogeny co-estimation has not been investigated. Second, the existing implementation of this approach is extraordinarily computationally intensive, which might necessitate trade-offs in practice that could partially undermine the method’s high accuracy. Finally, a recent study of alignment accuracy found that BAli-Phy’s co-estimation approach was accurate only when sequence data was simulated and not when biological benchmark data sets were used to evaluate alignment accuracy (Nute et al. 2019), suggesting the exceptional accuracy of this approach could be partially explained by strong similarity between the simulation model and that used to analyze the data, which might not translate to high accuracy on biological data. By using Bayesian Markov chain Monte Carlo to sample alignments, phylogenies and ancestral sequences from the posterior probability distribution, BAli-Phy implements an elegant approach at alignment-integrated ASR with a stronger formal justification than the heuristic method we present here. However, it is unknown whether integrating over the uncertainty associated with a single alignment model will accrue the same benefits as integrating many different alignment algorithms. Future studies will be needed to address these questions before the BAli-Phy approach or similar methods can be recommended in general for ancestral sequence reconstruction.

## Methods

### Software and Data Availability

All analyses presented in this study were performed using objective, transparent, reproducible algorithms documented in readable source code. All input data and analysis/visualization scripts are freely available under the General Public License (GPL) as open-access documentation associated with this publication at: https://github.com/bryankolaczkowski/airas

### Empirical Sequence Simulations

Empirical structures of diverse CARD, DSRM and RD domains were obtained from the protein data bank (Berman et al. 2000) and edited to remove any ligands or structural data from outside the annotated domain of interest. Structures from each domain family were aligned using the iterative_structural_align function in MODELLER v9.19 (Sali and Blundell 1993; Madhusudhan et al. 2009) to generate a multiple sequence alignment based primarily on structural superposition. This alignment was further edited manually to ensure that all aligned residues overlapped in the aligned structures.

Sequence data sets and consensus phylogenies for each domain family were curated from previous studies of DSRM (Dias et al. 2017), RIG-like receptor (Mukherjee et al. 2014; Pugh et al. 2016) and CARD-domain (Korithoski et al. 2015) families. Sequences were aligned to the structure-based alignment using the --seed option in mafft ginsi v7.402 (Katoh et al. 2002), and sequence regions not globally alignable to the structure-based alignment were trimmed. In order to simulate sequences with more realistic distributions of insertions and deletions (indels) across the sequence, we used the distribution of indels in the structure-guided alignment to determine the placement of indels in simulated sequences. Positions in the structure-guided alignment having at least 3 contiguous non-gap residues were considered impermissible to indels for the purposes of sequence simulation, whereas indels were allowed at all other positions in the alignment.

Simulation of 10 replicate data sets for each protein domain family—including correct ancestral sequences at each node on the phylogeny—was performed using indel-seq-gen v2.1.03 (Strope et al. 2007), assuming the consensus phylogeny, the JTT evolutionary model (Jones et al. 1992) and a 4-category discrete gamma model of among-site rate variation with shape parameter α=1.75 (Yang 1994). For each replicate, the root sequence was generated randomly from the structure-guided multiple sequence alignment by sampling amino-acid residues at each position based on the frequency of the amino-acid at that position. Columns in the multiple sequence alignment with >50% gap characters were not sampled when generating root sequences. Insertions and deletions were generated at permissible positions using the distributions from (Chang and Benner 2004), with a maximum indel size of 2 for CARD and DSRM3 domains, 4 for DSRM1 and DSRM2 domains, and 5 for the RD domain.

### Sequence Alignment

Simulated sequences were aligned using clustalw v2.1 (Sievers et al. 2011), mafft ginsi v7.402 (Katoh et al. 2002), msaprobs v0.9.7 (Liu et al. 2010), muscle v3.8.31 (Edgar 2004), probalign v1.4 (Roshan and Livesay 2006), probcons v1.12 (Do et al. 2005) and tcoffee v10.00.r1613 (Notredame et al. 2000), all with default parameters. In addition to sequence-based alignments, structure-guided alignments were generated by aligning each set of simulated sequences to the structure-based alignment (see above) using the --seed option in mafft ginsi (Katoh et al. 2002). Alignment errors were quantified by measuring the distance of each sequence alignment from the correct simulated alignment, using the d_pos option in MetAl v1.1, which estimates the probability that a randomly-selected residue aligns to an incorrect position in a randomly-selected sequence (Blackburne and Whelan 2012).

### Ancestral Sequence Reconstruction

Ancestral sequences were reconstructed from each alignment using marginal reconstruction (Yang et al. 1995) implemented in RAxML v8.2.10 (Stamatakis 2014), assuming the correct phylogeny and evolutionary model but estimating branch lengths and model parameters from each input data set. Each sequence alignment was converted to a binary presence-absence alignment, and ancestral gap states were inferred by marginal reconstruction using the BINCAT model in RAxML (Lewis 2001; Stamatakis 2006), assuming the correct tree topology with branch lengths and model parameters estimated from each data set. If the posterior probability of the gap state was >0.5 in the presence-absence reconstruction, that position was reconstructed as a gap character; otherwise, the position was reconstructed as whichever amino-acid residue had the largest posterior probability in the sequence reconstruction.

Alignment-integrated ancestral sequence reconstructions were produced by respectively combining sequence-reconstruction posterior probabilities and presence-absence posterior probabilities across all sequence-alignment methods (excluding data from structure-guided and correct alignments), assuming equal prior weights over sequence alignments. Let *P_i_* be the prior weight of alignment method *i*, and *P*(*j,k,m* | *i*) be the probability of ancestral state *j* at sequence position *k* and node *m* on the phylogeny, assuming alignment method *i*. Then, the heuristic ‘alignment-integrated posterior probability’ of ancestral state *j* at position *k* and node *m* is given by:

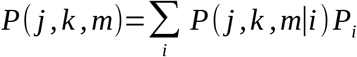

Here, we set the prior weight of each alignment method *P_i_* = 1/*n*, where *n* is the number of alignment methods.

Alignment-integration requires mapping all sequence alignments to one another, so that homologous columns from different alignments can be integrated. This was done using the – merge option in mafft ginsi.

After respective integration of sequence- and presence-absence reconstructions, the maximum a posteriori (MAP) ancestral sequence was generated as described above for single sequence-alignments.

Ancestral sequence reconstruction errors were calculated by comparing the MAP reconstructed ancestral state to the correct simulated ancestral state. For each inferred ancestral sequence, we calculated the number of errors divided by the length of the alignment generated by mapping the inferred ancestral sequence to the correct ancestral sequence. In addition to total ASR error rates, we also separately calculated the 3 possible types of ASR errors: 1) residue errors, in which both correct and inferred sequences have amino-acid residues at the same alignment position, but the inferred residue is different from the correct residue; 2) insertion errors, in which the correct ancestral sequence has a gap character, but the inferred sequence has a residue at that position, and 3) deletion errors, in which the correct sequence has a residue, but the inferred sequence has a gap. For each ancestral node on the phylogeny, we calculated the expected ASR error rate as the mean over 10 replicate data sets. Differences in error rates among methods across all replicates and nodes on the phylogeny were assessed using the 2-tailed independent 2-sample t-test, assuming unequal variances. Gaussian kernel density estimates were generated using least squares cross validation to estimate the smoothing parameter (Rudemo 1982).

### Probabilistic Sequence Alignment

Probabilistic sequence alignments were inferred using PRANK v170427 (Löytynoja 2014), with default parameters, and BAli-Phy v3.5 (Redelings and Suchard 2005). BAli-Phy analyses were conducted assuming the correct phylogeny, the JTT+G evolutionary model and the rs07 indel model (Redelings and Suchard 2007). Following the approach described in (Nute et al. 2019), we concatenated the Markov chain Monte Carlo samples from 32 independent BAli-Phy runs, each executed for a minimum of 1000 generations, after discarding the first 25% of samples from each run. The maximum a posteriori (MAP) alignment calculated over all BAli-Phy runs was used to reconstruct ancestral sequences, using the approach outlined above.

### Structural Modeling and RNA-Affinity Estimation

Structural homology models of DSRM1 domains were generated using MODELLER v9.19 (Sali and Blundell 1993). We used multi-template modeling (Larsson et al. 2008), assuming the structures and structure-based alignment generated for the DSRM1 domain simulations (see above). For each ancestral sequence, 50 models were generated and ranked using the MODELLER objective function, DOPE and DOPEHR assessment scores (Shen and Sali 2006). Each score was normalized by dividing it by its standard deviation across the 50 models, and we chose the best structural model as that with the optimal mean of normalized scores.

The structural stability of each protein structural model was inferred using DeltaGREM 2009, which estimates the change in free-energy/sequence-length of a given protein structure, compared to a statistical model of misfolded or unfolded protein ensembles, using a contact-based energy function (Minning et al. 2013; Bastolla 2014). We calculated structural stability errors as the absolute value of the difference in estimated stabilities between the correct ancestral sequence’s structural model and that of the inferred ancestral sequence. The expected stability error for each node on the phylogeny was calculated as the mean over 10 replicates.

DSRM1-dsRNA binding affinities were inferred using a previously-developed statistical machine learning approach (Dias and Kolazckowski 2015). For each ancestral sequence, a structural homology model was generated as described above, but including the dsRNA ligand from PDBID 5N8L. The pKd = -log10(Kd) was estimated using a support-vector machine trained on a large ensemble of protein-RNA and protein-DNA complexes with associated empirically-determined binding affinities. Errors in affinity predictions were calculated as the absolute value of the difference in estimated affinities between the correct ancestral sequence’s protein-RNA structural model and that of the inferred ancestral sequence, with expected errors calculated as the mean over 10 replicates.

### 3-Taxon Simulations

The JTT+G model (4-catetgory discrete gamma approximation with shape parameter α=1.75) was used to simulate 100 replicate data sets along 3-taxon phylogenies with branch lengths ranging from 0.1 to 0.8 substitutions/site (see Supplementary Information, Fig. S9). Starting sequences of 200 residues were generated at random from the JTT amino-acid frequency distribution and ‘evolved’ along the phylogeny using indel-seq-gen v2.1.03 (Strope et al. 2007). Insertions and deletions were generated at random, with the indel rate varying from 0.001 to 0.05 times the branch length (Pervez et al. 2014). Indel length was capped at 20 residues, with the length distribution of insertions and deletions taken from (Chang and Benner 2004).

## Abbreviations

ASR: ancestral sequence reconstruction
ML: maximum likelihood
MAP: maximum a posteriori
CARD: caspase activation and recruitment domain
DSRM: doublestranded RNA-binding motif
RD: RNA recognition domain
indel: insertion-deletion
PP: posterior probability
dsRNA: double-stranded RNA

