## SupplementaryMaterial for "Alignment-Integrated Reconstruction of Ancestral Sequences Improves Accuracy"

### Supplementary Information

#### Supplementary Tables

| domain | tree depth | max distance to root | branch lengths | | |
|---|---|---|---|---|---|
| | | | median | mean (stdev) | maximum |
| CARD | 39 | 5.11 | 0.0141 | 0.067 (0.179) | 3.19 |
| DSRM1 | 43 | 4.41 | 0.0173 | 0.099 (0.227) | 2.63 |
| DSRM2 | 44 | 4.41 | 0.0156 | 0.092 (0.212) | 2.37 |
| DSRM3 | 43 | 3.50 | 0.0029 | 0.042 (0.152) | 2.34 |
| RD | 35 | 3.33 | 0.0055 | 0.038 (0.101) | 1.90 |

**Table S1. Empirical protein domain family phylogenies used to simulate sequence data.** We simulated protein sequence data along empirical phylogenies with the indicated properties. Tree depth indicates the number of nodes on the longest path from the root to any leaf node on the phylogeny. Max distance to root indicates the largest cumulative number of expected substitutions/site along any path from the root to any leaf node. Branch length summary statistics are also indicated.

| domain | alignment | alignment_len | P(gap) | P(variable) | P(parsimony) |
|---|---|---|---|---|---|
| CARD | correct | 530.5 (5.4513) | 0.84 (0.0016) | 0.32 (0.0031) | 0.25 (0.0024) |
| 1273 sequences | structaligned | 283.8 (5.3329) | 0.71 (0.0044) | 0.88 (0.0075) | 0.68 (0.0051) |
| | aligned_clustalw | 123.6 (1.7333) | 0.33 (0.0060) | 0.98 (0.0014) | 0.97 (0.0014) |
| | aligned_mafft | 365.9 (10.0316) | 0.77 (0.0076) | 0.85 (0.0051) | 0.67 (0.0107) |
| | aligned_msaprobs | 270.5 (15.3096) | 0.69 (0.0160) | 0.92 (0.0041) | 0.82 (0.0008) |
| | aligned_muscle | 428.1 (14.1378) | 0.81 (0.0079) | 0.91 (0.0085) | 0.77 (0.0143) |
| | aligned_probalign | 353.6 (18.4272) | 0.76 (0.0107) | 0.93 (0.0069) | 0.85 (0.0150) |
| | aligned_probcons | 422.0 (14.5304) | 0.80 (0.0063) | 0.87 (0.0078) | 0.64 (0.0231) |
| | aligned_tcoffee | 394.9 (8.0380) | 0.79 (0.0056) | 0.80 (0.0155) | 0.61 (0.0064) |
| DSRM1 | correct | 180.4 (4.8378) | 0.62 (0.0089) | 0.50 (0.0097) | 0.45 (0.0104) |
| 1376 sequences | structaligned | 124.1 (2.7787) | 0.45 (0.0117) | 0.89 (0.0113) | 0.77 (0.0166) |
| | aligned_clustalw | 90.0 (1.3250) | 0.25 (0.0105) | 0.98 (0.0045) | 0.95 (0.0082) |
| | aligned_mafft | 270.3 (10.9990) | 0.75 (0.0111) | 0.89 (0.0150) | 0.72 (0.0219) |
| | aligned_msaprobs | 159.7 (6.4894) | 0.57 (0.0163) | 0.85 (0.0186) | 0.70 (0.0189) |
| | aligned_muscle | 204.5 (11.3463) | 0.66 (0.0173) | 0.92 (0.0138) | 0.78 (0.0160) |
| | aligned_probalign | 232.2 (13.5809) | 0.70 (0.0183) | 0.93 (0.0104) | 0.85 (0.0218) |
| | aligned_probcons | 160.5 (4.5662) | 0.58 (0.0113) | 0.79 (0.0200) | 0.64 (0.0186) |
| | aligned_tcoffee | 197.3 (7.9233) | 0.65 (0.0127) | 0.79 (0.0241) | 0.64 (0.0215) |
| DSRM2 | correct | 263.1 (8.0518) | 0.74 (0.0070) | 0.42 (0.0061) | 0.34 (0.0066) |
| 1497 sequences | structaligned | 156.2 (5.0789) | 0.56 (0.0118) | 0.89 (0.0114) | 0.75 (0.0116) |
| | aligned_clustalw | 97.8 (1.7876) | 0.30 (0.0104) | 0.99 (0.0048) | 0.95 (0.0077) |
| | aligned_mafft | 302.2 (6.4770) | 0.77 (0.0043) | 0.88 (0.0074) | 0.72 (0.0150) |
| | aligned_msaprobs | 187.0 (3.9861) | 0.63 (0.0068) | 0.85 (0.0169) | 0.68 (0.0181) |
| | aligned_muscle | 247.5 (13.4075) | 0.72 (0.0157) | 0.92 (0.0137) | 0.78 (0.0155) |
| | aligned_probalign | 265.5 (16.1983) | 0.73 (0.0218) | 0.93 (0.0137) | 0.83 (0.0256) |
| | aligned_probcons | 197.1 (7.8407) | 0.65 (0.0125) | 0.82 (0.0114) | 0.64 (0.0219) |
| | aligned_tcoffee | 253.8 (14.2773) | 0.72 (0.0156) | 0.79 (0.0178) | 0.65 (0.0147) |
| DSRM3 | correct | 86.2 (1.8123) | 0.19 (0.0172) | 0.84 (0.0143) | 0.82 (0.0170) |
| 718 sequences | structaligned | 84.9 (1.7729) | 0.18 (0.0169) | 0.91 (0.0120) | 0.86 (0.0124) |
| | aligned_clustalw | 77.3 (0.9667) | 0.10 (0.0099) | 0.98 (0.0050) | 0.95 (0.0076) |
| | aligned_mafft | 109.5 (6.8219) | 0.35 (0.0408) | 0.86 (0.0215) | 0.72 (0.0352) |
| | aligned_msaprobs | 86.2 (2.2988) | 0.19 (0.0212) | 0.89 (0.0184) | 0.84 (0.0177) |
| | aligned_muscle | 85.9 (1.5878) | 0.19 (0.0158) | 0.93 (0.0114) | 0.86 (0.0137) |

| | | | | | |
|---|---|---|---|---|---|
| | aligned_probalign | 125.2 (5.9643) | 0.44 (0.0279) | 0.87 (0.0145) | 0.69 (0.0265) |
| | aligned_probcons | 86.0 (2.0494) | 0.19 (0.0187) | 0.91 (0.0126) | 0.85 (0.0156) |
| | aligned_tcoffee | 107.9 (4.6078) | 0.35 (0.0271) | 0.80 (0.0189) | 0.71 (0.0230) |
| RD | correct | 256.1 (7.2823) | 0.54 (0.0115) | 0.54 (0.0115) | 0.52 (0.0109) |
| 935 sequences | structaligned | 195.3 (3.9131) | 0.39 (0.0095) | 0.81 (0.0084) | 0.71 (0.0080) |
| | aligned_clustalw | 162.9 (2.2083) | 0.27 (0.0085) | 0.96 (0.0058) | 0.93 (0.0051) |
| | aligned_mafft | 197.5 (4.9424) | 0.40 (0.0119) | 0.87 (0.0080) | 0.76 (0.0129) |
| | aligned_msaprobs | 208.8 (4.9549) | 0.43 (0.0116) | 0.75 (0.0126) | 0.65 (0.0119) |
| | aligned_muscle | 197.3 (3.9834) | 0.40 (0.0121) | 0.89 (0.0066) | 0.77 (0.0118) |
| | aligned_probalign | 209.0 (4.1339) | 0.43 (0.0101) | 0.74 (0.0080) | 0.66 (0.0099) |
| | aligned_probcons | 208.1 (4.4857) | 0.43 (0.0111) | 0.76 (0.0113) | 0.67 (0.0131) |
| | aligned_tcoffee | 212.4 (5.6592) | 0.44 (0.0130) | 0.78 (0.0115) | 0.71 (0.0143) |

**Table S2. Structure-guided and sequence-alignment methods typically underestimate correct alignment lengths and gap proportions while overestimating the proportions of variable and parsimony-informative sites.** We simulated alignments of 5 protein domain families, using empirically-derived simulation conditions, and aligned the resulting replicate sequence data using structure-guided and 7 different sequence-alignment methods (see Methods). For each alignment method and protein domain, we report the mean and standard error (in parentheses, over 10 replicates) in total alignment length (alignment_len), the proportions of gap characters (P(gap)), variable sites (P(variable)) and parsimony-informative sites (P(parsimony)). The number of sequences in each protein domain tree is also reported.

| domain | alignment | meanDistance | medianDistance | highestMode | SEmean |
|---|---|---|---|---|---|
| CARD | structaligned | 6.44E-01 | 6.55E-01 | 6.55E-01 | 5.15E-03 |
| | aligned_clustalw | 7.46E-01 | 7.59E-01 | 7.58E-01 | 8.43E-03 |
| | aligned_mafft | 7.42E-01 | 7.42E-01 | 7.44E-01 | 2.42E-03 |
| | aligned_msaprobs | 6.74E-01 | 6.63E-01 | 6.62E-01 | 8.05E-03 |
| | aligned_muscle | 7.94E-01 | 7.97E-01 | 7.97E-01 | 5.73E-03 |
| | aligned_probalign | 7.79E-01 | 7.78E-01 | 7.74E-01 | 5.25E-03 |
| | aligned_probcons | 6.47E-01 | 6.52E-01 | 6.54E-01 | 6.44E-03 |
| | aligned_tcoffee | 7.02E-01 | 7.06E-01 | 7.07E-01 | 4.16E-03 |
| DSRM1 | structaligned | 1.03E-01 | 9.29E-02 | 7.76E-02 | 1.42E-02 |
| | aligned_clustalw | 3.36E-01 | 3.33E-01 | 3.22E-01 | 3.75E-02 |
| | aligned_mafft | 3.83E-01 | 3.92E-01 | 3.78E-01 | 3.23E-02 |
| | aligned_msaprobs | 1.93E-01 | 1.95E-01 | 2.05E-01 | 2.36E-02 |
| | aligned_muscle | 3.18E-01 | 3.18E-01 | 2.40E-01 | 3.19E-02 |
| | aligned_probalign | 5.38E-01 | 5.06E-01 | 4.85E-01 | 3.20E-02 |
| | aligned_probcons | 1.65E-01 | 1.57E-01 | 1.28E-01 | 2.59E-02 |
| | aligned_tcoffee | 2.65E-01 | 2.87E-01 | 3.22E-01 | 2.23E-02 |
| DSRM2 | structaligned | 1.73E-01 | 1.57E-01 | 1.35E-01 | 2.66E-02 |
| | aligned_clustalw | 4.20E-01 | 4.42E-01 | 4.46E-01 | 2.24E-02 |
| | aligned_mafft | 4.97E-01 | 4.85E-01 | 4.82E-01 | 2.64E-02 |
| | aligned_msaprobs | 2.73E-01 | 2.55E-01 | 2.47E-01 | 1.93E-02 |
| | aligned_muscle | 4.03E-01 | 4.03E-01 | 3.91E-01 | 2.32E-02 |
| | aligned_probalign | 5.44E-01 | 5.34E-01 | 5.53E-01 | 2.69E-02 |
| | aligned_probcons | 2.43E-01 | 2.47E-01 | 2.44E-01 | 2.19E-02 |
| | aligned_tcoffee | 4.27E-01 | 4.18E-01 | 4.11E-01 | 1.62E-02 |
| DSRM3 | structaligned | 2.08E-02 | 1.50E-02 | 1.27E-02 | 6.14E-03 |
| | aligned_clustalw | 1.10E-01 | 9.95E-02 | 6.88E-02 | 2.02E-02 |
| | aligned_mafft | 8.08E-02 | 3.16E-02 | 2.62E-02 | 2.83E-02 |
| | aligned_msaprobs | 5.54E-02 | 2.70E-02 | 2.83E-02 | 2.01E-02 |
| | aligned_muscle | 9.38E-02 | 5.12E-02 | 3.87E-02 | 2.82E-02 |
| | aligned_probalign | 2.50E-01 | 2.07E-01 | 1.57E-01 | 4.98E-02 |
| | aligned_probcons | 9.15E-02 | 3.84E-02 | 3.66E-02 | 3.37E-02 |
| | aligned_tcoffee | 1.00E-01 | 5.49E-02 | 5.22E-02 | 3.23E-02 |

| | | | | | |
|---|---|---|---|---|---|
| RD | structaligned | 1.93E-02 | 1.79E-02 | 1.35E-02 | 3.79E-03 |
| | aligned_clustalw | 1.68E-01 | 1.64E-01 | 1.58E-01 | 1.06E-02 |
| | aligned_mafft | 5.17E-02 | 5.39E-02 | 5.53E-02 | 4.18E-03 |
| | aligned_msaprobs | 4.37E-02 | 4.80E-02 | 5.35E-02 | 5.06E-03 |
| | aligned_muscle | 5.32E-02 | 5.79E-02 | 6.90E-02 | 7.82E-03 |
| | aligned_probalign | 4.96E-02 | 5.60E-02 | 5.79E-02 | 5.79E-03 |
| | aligned_probcons | 5.15E-02 | 5.67E-02 | 6.06E-02 | 7.01E-03 |
| | aligned_tcoffee | 9.12E-02 | 9.00E-02 | 8.99E-02 | 6.43E-03 |

**Table S3. Alignment distances differ across alignment methods and protein domain families.** We simulated alignments of 5 protein domain families, using empirically-derived simulation conditions, and aligned the resulting replicate sequence data using structure-guided and 7 different sequence-alignment methods (see Methods). For each sequence alignment, we measured the position-wise distance of that alignment to the correct simulated alignment, which estimates the probability of randomly selecting an incorrectly-aligned residue from the inferred alignment (see Methods). We estimated the distribution of alignment distances across replicate simulations using kernel density estimation. We report the mean (and standard error), median and mode of each alignment-distance distribution.

| errorType | alignment | meanErrorRate | medianErrorRate | highestMode | SEmean |
|---|---|---|---|---|---|
| Total | aligned_clustalw | 6.31E-02 | 6.12E-02 | 4.20E-02 | 3.41E-04 |
| | aligned_mafft | 5.89E-02 | 5.63E-02 | 5.68E-02 | 4.43E-04 |
| | aligned_msaprobs | 3.85E-02 | 2.99E-02 | 1.65E-02 | 3.61E-04 |
| | aligned_muscle | 4.97E-02 | 4.24E-02 | 3.21E-02 | 4.08E-04 |
| | aligned_probalign | 6.76E-02 | 6.09E-02 | 3.70E-02 | 6.06E-04 |
| | aligned_probcons | 3.84E-02 | 3.02E-02 | 1.73E-02 | 3.51E-04 |
| | aligned_tcoffee | 5.63E-02 | 5.16E-02 | 3.79E-02 | 3.17E-04 |
| | structaligned | 3.49E-02 | 2.45E-02 | 1.48E-02 | 3.86E-04 |
| | correctaligned | 7.85E-03 | 2.50E-03 | 8.23E-04 | 1.96E-04 |
| | integratedaln | 2.88E-02 | 2.07E-02 | 1.40E-02 | 2.81E-04 |
| Residues | aligned_clustalw | 2.66E-02 | 2.46E-02 | 2.53E-02 | 2.17E-04 |
| | aligned_mafft | 1.91E-02 | 1.81E-02 | 1.88E-02 | 1.99E-04 |
| | aligned_msaprobs | 1.61E-02 | 1.18E-02 | 5.30E-03 | 2.02E-04 |
| | aligned_muscle | 2.09E-02 | 1.42E-02 | 9.42E-03 | 2.32E-04 |
| | aligned_probalign | 2.57E-02 | 1.77E-02 | 9.42E-03 | 3.13E-04 |
| | aligned_probcons | 1.75E-02 | 1.36E-02 | 5.30E-03 | 2.17E-04 |
| | aligned_tcoffee | 1.59E-02 | 1.16E-02 | 5.89E-03 | 2.06E-04 |
| | structaligned | 1.31E-02 | 7.70E-03 | 4.12E-03 | 1.95E-04 |
| | correctaligned | 6.18E-03 | 8.00E-04 | 5.89E-04 | 1.95E-04 |
| | integratedaln | 1.52E-02 | 1.07E-02 | 5.30E-03 | 2.03E-04 |
| Insertions | aligned_clustalw | 1.82E-02 | 1.41E-02 | 1.25E-02 | 1.19E-04 |
| | aligned_mafft | 2.02E-02 | 1.87E-02 | 1.87E-02 | 1.65E-04 |
| | aligned_msaprobs | 1.12E-02 | 7.40E-03 | 6.18E-03 | 1.22E-04 |
| | aligned_muscle | 1.52E-02 | 1.20E-02 | 1.08E-02 | 1.54E-04 |
| | aligned_probalign | 2.10E-02 | 1.99E-02 | 1.45E-02 | 1.73E-04 |
| | aligned_probcons | 1.05E-02 | 7.50E-03 | 6.56E-03 | 1.13E-04 |
| | aligned_tcoffee | 2.03E-02 | 1.88E-02 | 1.62E-02 | 1.03E-04 |
| | structaligned | 1.10E-02 | 6.80E-03 | 6.37E-03 | 1.38E-04 |
| | correctaligned | 8.49E-04 | 0.00E+00 | 0.00E+00 | 1.95E-05 |
| | integratedaln | 3.65E-03 | 2.60E-03 | 2.32E-03 | 3.82E-05 |
| Deletions | aligned_clustalw | 1.83E-02 | 1.42E-02 | 1.27E-02 | 1.19E-04 |
| | aligned_mafft | 1.97E-02 | 1.86E-02 | 1.85E-02 | 1.58E-04 |
| | aligned_msaprobs | 1.12E-02 | 7.40E-03 | 6.24E-03 | 1.22E-04 |
| | aligned_muscle | 1.37E-02 | 1.17E-02 | 1.13E-02 | 1.21E-04 |
| | aligned_probalign | 2.09E-02 | 1.99E-02 | 1.44E-02 | 1.72E-04 |
| | aligned_probcons | 1.04E-02 | 7.40E-03 | 6.63E-03 | 1.10E-04 |
| | aligned_tcoffee | 2.01E-02 | 1.85E-02 | 1.62E-02 | 1.01E-04 |
| | structaligned | 1.09E-02 | 6.80E-03 | 6.44E-03 | 1.36E-04 |
| | correctaligned | 8.17E-04 | 0.00E+00 | 0.00E+00 | 1.87E-05 |
| | integratedaln | 9.99E-03 | 7.00E-03 | 6.44E-03 | 1.17E-04 |

**Table S4. Ancestral sequence reconstruction (ASR) error rates are low when the correct alignment is known in advance, higher when the alignment is inferred using sequence-alignment methods, and intermediate when structure-guided alignments or alignment-integrated ASR is used.** We simulated extant and ancestral sequences for 5 protein domain families, using empirically-derived conditions, and aligned the resulting extant sequence data using structure-guided and 7 different sequence-alignment methods (see Methods). We used each alignment to infer the most likely ancestral sequence at each node on the phylogeny. Additionally, alignment-integrated ancestral sequences were generated by combining inferences from the 7 sequence-alignment methods (see Methods). In each case, we compared the inferred ancestral sequences to the correct simulated sequence to estimate ancestral sequence reconstruction (ASR) error rates (see Methods). ASR errors were divided into 4 “errorType” categories: 1) residue errors, in which both correct and inferred ancestral sequences have a residue at a given position in the alignment, but the residues differ; 2) insertion errors, in which the inferred sequence has a residue at a given alignment position, but the correct sequence has a gap state; 3) deletion errors, in which the inferred sequence has a gap, but the correct sequence has a residue, and 4) total errors, which combine residue- insertion- and deletion-errors. Sequence-wide error rates (errors/site) were computed by dividing the number of errors by the length of the pairwise alignment of the inferred and correct ancestral sequences. Error rate distributions were inferred by kernel density estimation. We report the mean (and standard error), median and mode of each ASR-error distribution.

| domain | alignment | meanErrorRate | medianErrorRate | highestMode | SEmean |
|---|---|---|---|---|---|
| CARD | aligned_clustalw | 9.21E-02 | 9.15E-02 | 8.95E-02 | 9.60E-05 |
| | aligned_mafft | 9.53E-02 | 1.03E-01 | 1.06E-01 | 4.68E-04 |
| | aligned_msaprobs | 7.50E-02 | 8.03E-02 | 5.17E-02 | 5.26E-04 |
| | aligned_muscle | 8.72E-02 | 9.83E-02 | 1.05E-01 | 6.51E-04 |
| | aligned_probalign | 8.57E-02 | 8.77E-02 | 6.22E-02 | 5.71E-04 |
| | aligned_probcons | 7.26E-02 | 7.52E-02 | 5.29E-02 | 4.63E-04 |
| | aligned_tcoffee | 8.34E-02 | 8.56E-02 | 6.62E-02 | 3.91E-04 |
| | structaligned | 8.04E-02 | 8.28E-02 | 6.45E-02 | 3.72E-04 |
| | correctaligned | 1.04E-02 | 8.40E-03 | 7.47E-03 | 1.86E-04 |
| | integratedaln | 5.56E-02 | 5.82E-02 | 4.27E-02 | 2.96E-04 |
| DSRM1 | aligned_clustalw | 4.90E-02 | 4.07E-02 | 3.78E-02 | 4.34E-04 |
| | aligned_mafft | 5.52E-02 | 5.40E-02 | 3.49E-02 | 6.34E-04 |
| | aligned_msaprobs | 3.15E-02 | 2.61E-02 | 1.74E-02 | 4.33E-04 |
| | aligned_muscle | 4.47E-02 | 3.69E-02 | 2.81E-02 | 4.78E-04 |
| | aligned_probalign | 7.56E-02 | 6.61E-02 | 3.49E-02 | 1.00E-03 |
| | aligned_probcons | 2.76E-02 | 2.13E-02 | 1.71E-02 | 3.94E-04 |
| | aligned_tcoffee | 4.53E-02 | 4.08E-02 | 3.97E-02 | 3.97E-04 |
| | structaligned | 2.21E-02 | 1.63E-02 | 1.45E-02 | 3.70E-04 |
| | correctaligned | 7.49E-03 | 1.00E-03 | 9.68E-04 | 3.72E-04 |
| | integratedaln | 2.18E-02 | 1.78E-02 | 1.65E-02 | 3.53E-04 |
| DSRM2 | aligned_clustalw | 5.44E-02 | 5.18E-02 | 4.33E-02 | 3.24E-04 |
| | aligned_mafft | 6.74E-02 | 6.41E-02 | 6.34E-02 | 4.69E-04 |
| | aligned_msaprobs | 3.26E-02 | 2.64E-02 | 2.52E-02 | 4.14E-04 |
| | aligned_muscle | 4.77E-02 | 3.71E-02 | 3.38E-02 | 4.48E-04 |
| | aligned_probalign | 6.58E-02 | 4.63E-02 | 4.22E-02 | 7.38E-04 |
| | aligned_probcons | 3.30E-02 | 3.03E-02 | 3.01E-02 | 3.50E-04 |
| | aligned_tcoffee | 5.61E-02 | 5.16E-02 | 5.05E-02 | 4.35E-04 |
| | structaligned | 3.01E-02 | 2.57E-02 | 2.50E-02 | 2.64E-04 |
| | correctaligned | 5.95E-03 | 8.00E-04 | 8.60E-04 | 2.87E-04 |
| | integratedaln | 2.36E-02 | 1.86E-02 | 1.72E-02 | 3.06E-04 |
| DSRM3 | aligned_clustalw | 4.43E-02 | 2.35E-02 | 1.98E-02 | 1.40E-03 |
| | aligned_mafft | 2.91E-02 | 9.20E-03 | 6.59E-03 | 1.41E-03 |
| | aligned_msaprobs | 2.26E-02 | 9.20E-03 | 6.59E-03 | 1.09E-03 |
| | aligned_muscle | 3.11E-02 | 8.50E-03 | 6.59E-03 | 1.36E-03 |
| | aligned_probalign | 8.17E-02 | 1.94E-02 | 1.65E-02 | 3.39E-03 |
| | aligned_probcons | 3.03E-02 | 7.90E-03 | 5.76E-03 | 1.35E-03 |
| | aligned_tcoffee | 4.01E-02 | 2.28E-02 | 1.89E-02 | 1.25E-03 |
| | structaligned | 1.47E-02 | 5.90E-03 | 6.59E-03 | 1.02E-03 |
| | correctaligned | 8.09E-03 | 6.00E-04 | 8.23E-04 | 9.72E-04 |
| | integratedaln | 2.07E-02 | 5.10E-03 | 3.29E-03 | 1.10E-03 |
| RD | aligned_clustalw | 7.25E-02 | 6.84E-02 | 7.09E-02 | 7.57E-04 |
| | aligned_mafft | 2.42E-02 | 1.94E-02 | 1.64E-02 | 6.65E-04 |
| | aligned_msaprobs | 2.06E-02 | 1.59E-02 | 1.49E-02 | 5.60E-04 |
| | aligned_muscle | 2.33E-02 | 1.86E-02 | 1.74E-02 | 6.14E-04 |
| | aligned_probalign | 2.30E-02 | 1.97E-02 | 1.93E-02 | 5.54E-04 |
| | aligned_probcons | 2.22E-02 | 1.86E-02 | 1.69E-02 | 5.86E-04 |
| | aligned_tcoffee | 4.83E-02 | 4.44E-02 | 3.95E-02 | 5.45E-04 |
| | structaligned | 1.51E-02 | 9.70E-03 | 9.16E-03 | 5.51E-04 |
| | correctaligned | 7.83E-03 | 1.40E-03 | 1.45E-03 | 5.80E-04 |
| | integratedaln | 1.74E-02 | 1.32E-02 | 1.21E-02 | 5.53E-04 |

**Table S5. Total ancestral sequence reconstruction (ASR) error rates are low when the correct alignment is known in advance, higher when the alignment is inferred using sequence-alignment methods, and intermediate when structure-guided alignments or alignment-integrated ASR is used.** We simulated extant and ancestral sequences for 5 protein domain families, using empirically-derived conditions, and aligned the resulting extant sequence data using structure-guided and 7 different sequence-alignment methods (see

Methods). We used each alignment to infer the most likely ancestral sequence at each node on the phylogeny. Additionally, alignment-integrated ancestral sequences were generated by combining inferences from the 7 sequence-alignment methods (see Methods). In each case, we compared the inferred ancestral sequences to the correct simulated sequence to estimate ancestral sequence reconstruction (ASR) error rates (see Methods). ASR errors were divided into 4 “errorType” categories: 1) residue errors, in which both correct and inferred ancestral sequences have a residue at a given position in the alignment, but the residues differ; 2) insertion errors, in which the inferred sequence has a residue at a given alignment position, but the correct sequence has a gap state; 3) deletion errors, in which the inferred sequence has a gap, but the correct sequence has a residue, and 4) total errors, which combine residue- insertion- and deletion-errors. Sequence-wide error rates (errors/site) were computed by dividing the number of errors by the length of the pairwise alignment of the inferred and correct ancestral sequences. Error rate distributions were inferred by kernel density estimation. For each protein domain family, we report the mean (and standard error), median and mode of the total ASR-error distribution.

| domain | alignment | meanErrorRate | medianErrorRate | highestMode | SEmean |
|---|---|---|---|---|---|
| CARD | aligned_clustalw | 2.68E-02 | 2.62E-02 | 2.58E-02 | 7.65E-05 |
| | aligned_mafft | 2.07E-02 | 2.44E-02 | 2.67E-02 | 2.05E-04 |
| | aligned_msaprobs | 2.10E-02 | 2.38E-02 | 2.50E-02 | 2.25E-04 |
| | aligned_muscle | 2.50E-02 | 2.96E-02 | 3.18E-02 | 2.89E-04 |
| | aligned_probalign | 2.44E-02 | 2.48E-02 | 1.64E-02 | 2.03E-04 |
| | aligned_probcons | 2.29E-02 | 2.53E-02 | 2.64E-02 | 1.77E-04 |
| | aligned_tcoffee | 1.93E-02 | 2.03E-02 | 1.20E-02 | 1.70E-04 |
| | structaligned | 2.10E-02 | 2.45E-02 | 2.54E-02 | 1.82E-04 |
| | correctaligned | 3.37E-03 | 1.10E-03 | 8.70E-04 | 1.75E-04 |
| | integratedaln | 2.13E-02 | 2.48E-02 | 2.61E-02 | 2.05E-04 |
| DSRM1 | aligned_clustalw | 2.61E-02 | 1.79E-02 | 1.66E-02 | 3.83E-04 |
| | aligned_mafft | 2.17E-02 | 1.83E-02 | 1.28E-02 | 3.60E-04 |
| | aligned_msaprobs | 1.61E-02 | 1.25E-02 | 1.13E-02 | 3.48E-04 |
| | aligned_muscle | 2.19E-02 | 1.59E-02 | 1.06E-02 | 3.75E-04 |
| | aligned_probalign | 2.83E-02 | 2.53E-02 | 7.45E-03 | 4.95E-04 |
| | aligned_probcons | 1.47E-02 | 9.70E-03 | 7.45E-03 | 3.47E-04 |
| | aligned_tcoffee | 1.39E-02 | 9.10E-03 | 7.45E-03 | 3.49E-04 |
| | structaligned | 1.08E-02 | 4.90E-03 | 4.26E-03 | 3.51E-04 |
| | correctaligned | 7.40E-03 | 9.00E-04 | 8.51E-04 | 3.71E-04 |
| | integratedaln | 1.31E-02 | 9.40E-03 | 6.81E-03 | 3.39E-04 |
| DSRM2 | aligned_clustalw | 2.58E-02 | 2.37E-02 | 1.62E-02 | 2.68E-04 |
| | aligned_mafft | 2.25E-02 | 1.98E-02 | 1.85E-02 | 2.40E-04 |
| | aligned_msaprobs | 1.56E-02 | 1.06E-02 | 9.80E-03 | 3.08E-04 |
| | aligned_muscle | 2.03E-02 | 1.13E-02 | 9.44E-03 | 3.49E-04 |
| | aligned_probalign | 2.43E-02 | 1.44E-02 | 1.19E-02 | 3.79E-04 |
| | aligned_probcons | 1.62E-02 | 1.34E-02 | 1.32E-02 | 2.83E-04 |
| | aligned_tcoffee | 1.66E-02 | 1.14E-02 | 8.55E-03 | 3.46E-04 |
| | structaligned | 1.22E-02 | 7.80E-03 | 7.13E-03 | 2.48E-04 |
| | correctaligned | 5.78E-03 | 6.00E-04 | 5.35E-04 | 2.85E-04 |
| | integratedaln | 1.32E-02 | 8.60E-03 | 6.59E-03 | 2.86E-04 |
| DSRM3 | aligned_clustalw | 2.86E-02 | 8.70E-03 | 7.07E-03 | 1.25E-03 |
| | aligned_mafft | 1.49E-02 | 5.10E-03 | 4.12E-03 | 9.87E-04 |
| | aligned_msaprobs | 1.52E-02 | 4.50E-03 | 2.95E-03 | 9.97E-04 |
| | aligned_muscle | 2.39E-02 | 5.30E-03 | 2.95E-03 | 1.21E-03 |
| | aligned_probalign | 4.30E-02 | 1.09E-02 | 5.89E-03 | 1.86E-03 |
| | aligned_probcons | 2.31E-02 | 4.15E-03 | 2.36E-03 | 1.20E-03 |
| | aligned_tcoffee | 1.91E-02 | 5.00E-03 | 2.36E-03 | 1.05E-03 |
| | structaligned | 9.25E-03 | 2.20E-03 | 2.36E-03 | 9.57E-04 |
| | correctaligned | 7.98E-03 | 5.00E-04 | 1.18E-03 | 9.66E-04 |
| | integratedaln | 1.79E-02 | 3.75E-03 | 1.77E-03 | 1.05E-03 |
| RD | aligned_clustalw | 2.68E-02 | 2.26E-02 | 2.28E-02 | 6.09E-04 |
| | aligned_mafft | 1.05E-02 | 5.40E-03 | 4.19E-03 | 5.67E-04 |
| | aligned_msaprobs | 1.10E-02 | 4.90E-03 | 4.95E-03 | 5.63E-04 |
| | aligned_muscle | 1.24E-02 | 6.40E-03 | 6.09E-03 | 5.78E-04 |
| | aligned_probalign | 1.25E-02 | 7.30E-03 | 6.47E-03 | 5.55E-04 |
| | aligned_probcons | 1.21E-02 | 6.60E-03 | 5.71E-03 | 5.69E-04 |
| | aligned_tcoffee | 1.09E-02 | 5.70E-03 | 4.95E-03 | 5.49E-04 |
| | structaligned | 9.76E-03 | 3.60E-03 | 3.81E-03 | 5.63E-04 |
| | correctaligned | 7.49E-03 | 1.00E-03 | 1.14E-03 | 5.75E-04 |
| | integratedaln | 1.11E-02 | 5.20E-03 | 4.95E-03 | 5.55E-04 |

**Table S6. Ancestral sequence reconstruction (ASR) residue-reconstruction error rates are low when the correct alignment is known in advance, higher when the alignment is inferred using sequence-alignment methods, and intermediate when structure-guided alignments or alignment-integrated ASR is used.** We simulated extant and ancestral sequences for 5 protein domain families, using empirically-derived conditions, and aligned the resulting extant sequence data using structure-guided and 7 different sequence-alignment methods (see Methods). We used each alignment to infer the most likely ancestral sequence at each node on the phylogeny. Additionally, alignment-integrated ancestral sequences were generated by combining inferences from the 7 sequence-alignment methods (see Methods). In each case, we compared the inferred ancestral sequences to the correct simulated sequence to estimate ancestral sequence reconstruction (ASR) error rates (see Methods). ASR errors were divided into 4 “errorType” categories: 1) residue errors, in which both correct and inferred ancestral sequences have a residue at a given position in the alignment, but the residues differ; 2) insertion errors, in which the inferred sequence has a residue at a given alignment position, but the correct sequence has a gap state; 3) deletion errors, in which the inferred sequence has a gap, but the correct sequence has a residue, and 4) total errors, which combine residue- insertion- and deletion-errors. Sequence-wide error rates (errors/site) were computed by dividing the number of errors by the length of the pairwise alignment of the inferred and correct ancestral sequences. Error rate distributions were inferred by kernel density estimation. For each protein domain family, we report the mean (and standard error), median and mode of the residue ASR-error distribution.

| domain | alignment | meanErrorRate | medianErrorRate | highestMode | SEmean |
|---|---|---|---|---|---|
| CARD | aligned_clustalw | 3.26E-02 | 3.26E-02 | 3.22E-02 | 3.52E-05 |
| | aligned_mafft | 3.77E-02 | 3.81E-02 | 3.14E-02 | 1.67E-04 |
| | aligned_msaprobs | 2.70E-02 | 2.65E-02 | 2.05E-02 | 1.74E-04 |
| | aligned_muscle | 3.37E-02 | 3.56E-02 | 2.28E-02 | 2.60E-04 |
| | aligned_probalign | 3.08E-02 | 2.96E-02 | 2.30E-02 | 2.28E-04 |
| | aligned_probcons | 2.50E-02 | 2.42E-02 | 1.87E-02 | 1.70E-04 |
| | aligned_tcoffee | 3.23E-02 | 3.24E-02 | 2.71E-02 | 1.33E-04 |
| | structaligned | 2.98E-02 | 2.89E-02 | 2.59E-02 | 1.15E-04 |
| | correctaligned | 3.55E-03 | 3.30E-03 | 3.07E-03 | 1.99E-05 |
| | integratedaln | 8.74E-03 | 9.20E-03 | 9.81E-03 | 4.13E-05 |
| DSRM1 | aligned_clustalw | 1.14E-02 | 1.13E-02 | 1.11E-02 | 3.90E-05 |
| | aligned_mafft | 1.70E-02 | 1.56E-02 | 1.13E-02 | 1.77E-04 |
| | aligned_msaprobs | 7.71E-03 | 6.50E-03 | 5.78E-03 | 7.39E-05 |
| | aligned_muscle | 1.17E-02 | 1.04E-02 | 9.45E-03 | 9.22E-05 |
| | aligned_probalign | 2.37E-02 | 2.03E-02 | 1.36E-02 | 2.75E-04 |
| | aligned_probcons | 6.48E-03 | 5.70E-03 | 5.23E-03 | 4.94E-05 |
| | aligned_tcoffee | 1.58E-02 | 1.55E-02 | 1.55E-02 | 7.17E-05 |
| | structaligned | 5.65E-03 | 5.60E-03 | 4.67E-03 | 2.61E-05 |
| | correctaligned | 5.14E-05 | 0.00E+00 | 0.00E+00 | 3.38E-06 |
| | integratedaln | 2.07E-03 | 1.80E-03 | 1.56E-03 | 2.06E-05 |
| DSRM2 | aligned_clustalw | 1.43E-02 | 1.39E-02 | 1.30E-02 | 4.24E-05 |
| | aligned_mafft | 2.28E-02 | 2.21E-02 | 1.89E-02 | 1.47E-04 |
| | aligned_msaprobs | 8.56E-03 | 7.70E-03 | 7.62E-03 | 7.22E-05 |
| | aligned_muscle | 1.42E-02 | 1.28E-02 | 1.21E-02 | 8.42E-05 |
| | aligned_probalign | 2.08E-02 | 1.61E-02 | 1.51E-02 | 1.99E-04 |
| | aligned_probcons | 8.42E-03 | 8.20E-03 | 8.18E-03 | 5.31E-05 |
| | aligned_tcoffee | 2.00E-02 | 2.00E-02 | 2.03E-02 | 7.62E-05 |
| | structaligned | 8.93E-03 | 8.90E-03 | 8.96E-03 | 3.19E-05 |
| | correctaligned | 9.36E-05 | 0.00E+00 | 0.00E+00 | 5.13E-06 |
| | integratedaln | 2.50E-03 | 2.40E-03 | 2.35E-03 | 1.09E-05 |
| DSRM3 | aligned_clustalw | 7.83E-03 | 7.80E-03 | 1.14E-02 | 1.08E-04 |
| | aligned_mafft | 7.10E-03 | 2.80E-03 | 1.74E-03 | 3.07E-04 |
| | aligned_msaprobs | 3.69E-03 | 2.20E-03 | 1.93E-03 | 9.47E-05 |
| | aligned_muscle | 3.55E-03 | 2.10E-03 | 1.54E-03 | 1.11E-04 |
| | aligned_probalign | 1.94E-02 | 5.60E-03 | 4.83E-03 | 7.89E-04 |
| | aligned_probcons | 3.55E-03 | 1.90E-03 | 1.74E-03 | 1.03E-04 |
| | aligned_tcoffee | 1.05E-02 | 8.50E-03 | 8.49E-03 | 1.62E-04 |
| | structaligned | 2.73E-03 | 1.40E-03 | 7.72E-04 | 1.05E-04 |
| | correctaligned | 4.55E-05 | 0.00E+00 | 0.00E+00 | 6.41E-06 |
| | integratedaln | 1.29E-03 | 1.00E-03 | 1.93E-04 | 4.31E-05 |
| RD | aligned_clustalw | 2.28E-02 | 2.26E-02 | 2.00E-02 | 1.02E-04 |
| | aligned_mafft | 6.83E-03 | 6.40E-03 | 3.84E-03 | 9.85E-05 |
| | aligned_msaprobs | 4.75E-03 | 4.70E-03 | 4.87E-03 | 4.96E-05 |
| | aligned_muscle | 5.36E-03 | 5.10E-03 | 5.83E-03 | 6.50E-05 |
| | aligned_probalign | 5.26E-03 | 5.10E-03 | 3.64E-03 | 5.73E-05 |
| | aligned_probcons | 5.04E-03 | 5.00E-03 | 3.22E-03 | 5.82E-05 |
| | aligned_tcoffee | 1.87E-02 | 1.81E-02 | 1.67E-02 | 8.59E-05 |
| | structaligned | 2.64E-03 | 2.70E-03 | 1.99E-03 | 2.90E-05 |
| | correctaligned | 1.72E-04 | 0.00E+00 | 0.00E+00 | 1.03E-05 |
| | integratedaln | 2.68E-03 | 2.60E-03 | 2.61E-03 | 3.26E-05 |

**Table S7. Ancestral sequence reconstruction (ASR) insertion error rates are low when the correct alignment is known in advance, higher when the alignment is inferred using sequence-alignment methods, and intermediate when structure-guided alignments or alignment-integrated ASR is used.** We simulated extant and ancestral sequences for 5 protein domain families, using empirically-derived conditions, and aligned the resulting extant sequence data using structure-guided and 7 different sequence-alignment methods (see

Methods). We used each alignment to infer the most likely ancestral sequence at each node on the phylogeny. Additionally, alignment-integrated ancestral sequences were generated by combining inferences from the 7 sequence-alignment methods (see Methods). In each case, we compared the inferred ancestral sequences to the correct simulated sequence to estimate ancestral sequence reconstruction (ASR) error rates (see Methods). ASR errors were divided into 4 “errorType” categories: 1) residue errors, in which both correct and inferred ancestral sequences have a residue at a given position in the alignment, but the residues differ; 2) insertion errors, in which the inferred sequence has a residue at a given alignment position, but the correct sequence has a gap state; 3) deletion errors, in which the inferred sequence has a gap, but the correct sequence has a residue, and 4) total errors, which combine residue- insertion- and deletion-errors. Sequence-wide error rates (errors/site) were computed by dividing the number of errors by the length of the pairwise alignment of the inferred and correct ancestral sequences. Error rate distributions were inferred by kernel density estimation. For each protein domain family, we report the mean (and standard error), median and mode of the insertion ASR-error distribution.

| domain | alignment | meanErrorRate | medianErrorRate | highestMode | SEmean |
|---|---|---|---|---|---|
| CARD | aligned_clustalw | 3.27E-02 | 3.27E-02 | 3.21E-02 | 3.25E-05 |
| | aligned_mafft | 3.70E-02 | 3.78E-02 | 3.13E-02 | 1.30E-04 |
| | aligned_msaprobs | 2.69E-02 | 2.62E-02 | 2.05E-02 | 1.73E-04 |
| | aligned_muscle | 2.85E-02 | 2.89E-02 | 2.25E-02 | 1.53E-04 |
| | aligned_probalign | 3.06E-02 | 2.95E-02 | 2.29E-02 | 2.25E-04 |
| | aligned_probcons | 2.46E-02 | 2.39E-02 | 1.87E-02 | 1.61E-04 |
| | aligned_tcoffee | 3.18E-02 | 3.19E-02 | 2.70E-02 | 1.27E-04 |
| | structaligned | 2.95E-02 | 2.86E-02 | 2.59E-02 | 1.05E-04 |
| | correctaligned | 3.44E-03 | 3.30E-03 | 3.17E-03 | 1.69E-05 |
| | integratedaln | 2.55E-02 | 2.47E-02 | 2.08E-02 | 1.42E-04 |
| DSRM1 | aligned_clustalw | 1.15E-02 | 1.13E-02 | 1.12E-02 | 3.81E-05 |
| | aligned_mafft | 1.65E-02 | 1.52E-02 | 1.13E-02 | 1.57E-04 |
| | aligned_msaprobs | 7.70E-03 | 6.50E-03 | 5.77E-03 | 7.37E-05 |
| | aligned_muscle | 1.12E-02 | 1.02E-02 | 9.71E-03 | 7.36E-05 |
| | aligned_probalign | 2.36E-02 | 2.03E-02 | 1.36E-02 | 2.73E-04 |
| | aligned_probcons | 6.48E-03 | 5.70E-03 | 5.22E-03 | 4.91E-05 |
| | aligned_tcoffee | 1.56E-02 | 1.54E-02 | 1.55E-02 | 7.40E-05 |
| | structaligned | 5.68E-03 | 5.70E-03 | 4.67E-03 | 2.63E-05 |
| | correctaligned | 4.75E-05 | 0.00E+00 | 0.00E+00 | 3.71E-06 |
| | integratedaln | 6.61E-03 | 6.50E-03 | 4.85E-03 | 4.18E-05 |
| DSRM2 | aligned_clustalw | 1.43E-02 | 1.39E-02 | 1.31E-02 | 4.18E-05 |
| | aligned_mafft | 2.21E-02 | 2.19E-02 | 1.88E-02 | 1.23E-04 |
| | aligned_msaprobs | 8.51E-03 | 7.70E-03 | 7.62E-03 | 7.12E-05 |
| | aligned_muscle | 1.32E-02 | 1.24E-02 | 1.20E-02 | 5.81E-05 |
| | aligned_probalign | 2.07E-02 | 1.60E-02 | 1.51E-02 | 1.98E-04 |
| | aligned_probcons | 8.39E-03 | 8.20E-03 | 8.23E-03 | 5.25E-05 |
| | aligned_tcoffee | 1.96E-02 | 1.99E-02 | 2.02E-02 | 7.65E-05 |
| | structaligned | 8.91E-03 | 8.90E-03 | 8.94E-03 | 3.09E-05 |
| | correctaligned | 6.94E-05 | 0.00E+00 | 0.00E+00 | 3.45E-06 |
| | integratedaln | 7.87E-03 | 7.60E-03 | 7.53E-03 | 4.42E-05 |
| DSRM3 | aligned_clustalw | 7.89E-03 | 7.80E-03 | 1.15E-02 | 1.09E-04 |
| | aligned_mafft | 7.12E-03 | 2.80E-03 | 1.76E-03 | 3.09E-04 |
| | aligned_msaprobs | 3.70E-03 | 2.20E-03 | 1.95E-03 | 9.53E-05 |
| | aligned_muscle | 3.63E-03 | 2.10E-03 | 1.56E-03 | 1.14E-04 |
| | aligned_probalign | 1.94E-02 | 5.60E-03 | 4.88E-03 | 7.89E-04 |
| | aligned_probcons | 3.57E-03 | 1.90E-03 | 1.76E-03 | 1.05E-04 |
| | aligned_tcoffee | 1.05E-02 | 8.50E-03 | 8.58E-03 | 1.64E-04 |
| | structaligned | 2.75E-03 | 1.40E-03 | 7.80E-04 | 1.07E-04 |
| | correctaligned | 5.89E-05 | 0.00E+00 | 0.00E+00 | 8.09E-06 |
| | integratedaln | 1.57E-03 | 1.00E-03 | 9.75E-04 | 4.80E-05 |
| RD | aligned_clustalw | 2.29E-02 | 2.26E-02 | 2.01E-02 | 1.04E-04 |
| | aligned_mafft | 6.89E-03 | 6.40E-03 | 3.84E-03 | 1.01E-04 |
| | aligned_msaprobs | 4.78E-03 | 4.70E-03 | 4.80E-03 | 4.91E-05 |
| | aligned_muscle | 5.55E-03 | 5.40E-03 | 5.69E-03 | 6.96E-05 |
| | aligned_probalign | 5.29E-03 | 5.10E-03 | 3.64E-03 | 5.69E-05 |
| | aligned_probcons | 5.08E-03 | 5.00E-03 | 3.22E-03 | 5.82E-05 |
| | aligned_tcoffee | 1.87E-02 | 1.81E-02 | 1.66E-02 | 8.59E-05 |
| | structaligned | 2.67E-03 | 2.70E-03 | 1.99E-03 | 2.81E-05 |
| | correctaligned | 1.62E-04 | 0.00E+00 | 0.00E+00 | 7.31E-06 |
| | integratedaln | 3.68E-03 | 3.50E-03 | 3.02E-03 | 5.24E-05 |

**Table S8. Ancestral sequence reconstruction (ASR) deletion error rates are low when the correct alignment is known in advance, higher when the alignment is inferred using sequence-alignment methods, and intermediate when structure-guided alignments or alignment-integrated ASR is used.** We simulated extant and ancestral sequences for 5 protein domain families, using empirically-derived conditions, and aligned the resulting extant sequence data using structure-guided and 7 different sequence-alignment methods (see

Methods). We used each alignment to infer the most likely ancestral sequence at each node on the phylogeny. Additionally, alignment-integrated ancestral sequences were generated by combining inferences from the 7 sequence-alignment methods (see Methods). In each case, we compared the inferred ancestral sequences to the correct simulated sequence to estimate ancestral sequence reconstruction (ASR) error rates (see Methods). ASR errors were divided into 4 "errorType" categories: 1) residue errors, in which both correct and inferred ancestral sequences have a residue at a given position in the alignment, but the residues differ; 2) insertion errors, in which the inferred sequence has a residue at a given alignment position, but the correct sequence has a gap state; 3) deletion errors, in which the inferred sequence has a gap, but the correct sequence has a residue, and 4) total errors, which combine residue- insertion- and deletion-errors. Sequence-wide error rates (errors/site) were computed by dividing the number of errors by the length of the pairwise alignment of the inferred and correct ancestral sequences. Error rate distributions were inferred by kernel density estimation. For each protein domain family, we report the mean (and standard error), median and mode of the deletion ASR-error distribution.

| errorType | alignment | meanErrorPP | medianErrorPP | highestMode | SEmean |
|---|---|---|---|---|---|
| Total | aligned_clustalw | 9.48E-01 | 9.94E-01 | 9.96E-01 | 1.36E-03 |
| | aligned_mafft | 9.37E-01 | 9.92E-01 | 9.94E-01 | 1.50E-03 |
| | aligned_msaprobs | 9.24E-01 | 9.83E-01 | 9.94E-01 | 1.56E-03 |
| | aligned_muscle | 9.02E-01 | 9.64E-01 | 9.90E-01 | 1.72E-03 |
| | aligned_probalign | 9.35E-01 | 9.89E-01 | 9.94E-01 | 1.43E-03 |
| | aligned_probcons | 9.22E-01 | 9.83E-01 | 9.94E-01 | 1.58E-03 |
| | aligned_tcoffee | 9.41E-01 | 9.92E-01 | 9.94E-01 | 1.47E-03 |
| | structaligned | 9.21E-01 | 9.84E-01 | 9.94E-01 | 1.55E-03 |
| | correctaligned | 6.46E-01 | 7.05E-01 | 6.82E-01 | 4.28E-03 |
| | integratedaln | 6.74E-01 | 6.86E-01 | 7.04E-01 | 1.35E-03 |
| Residues | aligned_clustalw | 9.23E-01 | 9.86E-01 | 9.94E-01 | 1.72E-03 |
| | aligned_mafft | 9.03E-01 | 9.78E-01 | 9.94E-01 | 1.88E-03 |
| | aligned_msaprobs | 8.91E-01 | 9.59E-01 | 9.94E-01 | 1.90E-03 |
| | aligned_muscle | 8.71E-01 | 9.40E-01 | 9.88E-01 | 2.11E-03 |
| | aligned_probalign | 9.05E-01 | 9.75E-01 | 9.92E-01 | 1.84E-03 |
| | aligned_probcons | 8.86E-01 | 9.58E-01 | 9.92E-01 | 2.07E-03 |
| | aligned_tcoffee | 8.93E-01 | 9.62E-01 | 9.92E-01 | 1.94E-03 |
| | structaligned | 8.89E-01 | 9.53E-01 | 9.94E-01 | 1.85E-03 |
| | correctaligned | 5.66E-01 | 6.56E-01 | 6.66E-01 | 4.33E-03 |
| | integratedaln | 5.86E-01 | 6.05E-01 | 6.18E-01 | 1.54E-03 |
| Insertions | aligned_clustalw | 9.50E-01 | 9.97E-01 | 9.96E-01 | 1.55E-03 |
| | aligned_mafft | 9.48E-01 | 9.98E-01 | 9.96E-01 | 1.65E-03 |
| | aligned_msaprobs | 9.43E-01 | 9.99E-01 | 9.96E-01 | 1.63E-03 |
| | aligned_muscle | 9.07E-01 | 9.85E-01 | 9.92E-01 | 2.04E-03 |
| | aligned_probalign | 9.40E-01 | 9.97E-01 | 9.96E-01 | 1.69E-03 |
| | aligned_probcons | 9.47E-01 | 9.99E-01 | 9.96E-01 | 1.55E-03 |
| | aligned_tcoffee | 9.42E-01 | 9.96E-01 | 9.96E-01 | 1.68E-03 |
| | structaligned | 9.54E-01 | 1.00E+00 | 9.98E-01 | 1.45E-03 |
| | correctaligned | 4.21E-01 | 0.00E+00 | 0.00E+00 | 6.30E-03 |
| | integratedaln | 7.17E-01 | 8.00E-01 | 8.58E-01 | 2.84E-03 |
| Deletions | aligned_clustalw | 9.98E-01 | 1.00E+00 | 1.00E+00 | 1.03E-04 |
| | aligned_mafft | 9.96E-01 | 1.00E+00 | 1.00E+00 | 2.68E-04 |
| | aligned_msaprobs | 9.99E-01 | 1.00E+00 | 1.00E+00 | 8.91E-05 |
| | aligned_muscle | 9.86E-01 | 1.00E+00 | 1.00E+00 | 3.10E-04 |
| | aligned_probalign | 9.98E-01 | 1.00E+00 | 1.00E+00 | 1.30E-04 |
| | aligned_probcons | 9.98E-01 | 1.00E+00 | 1.00E+00 | 1.12E-04 |
| | aligned_tcoffee | 9.97E-01 | 1.00E+00 | 1.00E+00 | 1.90E-04 |
| | structaligned | 9.97E-01 | 1.00E+00 | 1.00E+00 | 4.35E-04 |
| | correctaligned | 4.29E-01 | 0.00E+00 | 0.00E+00 | 6.47E-03 |
| | integratedaln | 7.46E-01 | 7.42E-01 | 7.32E-01 | 1.36E-03 |

**Table S9. Support for erroneously-inferred ancestral states is reduced by alignment-integrated ancestral sequence reconstruction.** We simulated extant and ancestral sequences for 5 protein domain families, using empirically-derived conditions, and aligned the resulting extant sequence data using structure-guided and 7 different sequence-alignment methods (see Methods). We used each alignment to infer the most likely ancestral sequence at each node on the phylogeny. Additionally, alignment-integrated ancestral sequences were generated by combining inferences from the 7 sequence-alignment methods (see Methods). In each case, we compared the inferred ancestral sequences to the correct simulated sequence to estimate ancestral sequence reconstruction (ASR) error rates (see Methods). ASR errors were divided into 4 “errorType” categories: 1) residue errors, in which both correct and inferred ancestral sequences have a residue at a given position in the alignment, but the residues differ; 2) insertion errors, in which the inferred sequence has a residue at a given alignment position, but the correct sequence has a gap state; 3) deletion errors, in which the inferred sequence has a gap, but the correct sequence has a residue, and 4) total errors, which combine residue-insertion- and deletion-errors. The posterior probability (PP) of each erroneously-inferred ancestral state was determined using an empirical-Bayesian approach (see Methods). The distribution of posterior probabilities for erroneously-inferred ancestral states was calculated by kernel density estimation. We report the mean (and standard error), median and mode of the posterior-probability distribution for each type of ASR error.

| errorType | alignment | meanCorrectPP | medianCorrectPP | highestMode | SEmean |
|---|---|---|---|---|---|
| Total | aligned_clustalw | 1.64E-02 | 2.60E-03 | 2.00E-03 | 3.94E-04 |
| | aligned_mafft | 2.66E-02 | 2.80E-03 | 2.00E-03 | 6.36E-04 |
| | aligned_msaprobs | 3.33E-02 | 7.20E-03 | 2.00E-03 | 6.48E-04 |
| | aligned_muscle | 4.53E-02 | 2.81E-02 | 4.00E-03 | 6.80E-04 |
| | aligned_probalign | 2.35E-02 | 3.90E-03 | 2.00E-03 | 5.70E-04 |
| | aligned_probcons | 3.68E-02 | 8.80E-03 | 2.00E-03 | 6.87E-04 |
| | aligned_tcoffee | 2.09E-02 | 4.00E-03 | 2.00E-03 | 4.61E-04 |
| | structaligned | 4.23E-02 | 8.10E-03 | 2.00E-03 | 8.03E-04 |
| | correctaligned | 1.39E-01 | 1.34E-01 | 2.00E-03 | 1.75E-03 |
| | integratedaln | 3.29E-01 | 3.29E-01 | 4.30E-01 | 1.69E-03 |
| Residues | aligned_clustalw | 2.98E-02 | 6.70E-03 | 1.99E-03 | 5.92E-04 |
| | aligned_mafft | 4.80E-02 | 9.90E-03 | 1.99E-03 | 9.13E-04 |
| | aligned_msaprobs | 5.49E-02 | 2.21E-02 | 2.99E-03 | 9.06E-04 |
| | aligned_muscle | 5.01E-02 | 2.21E-02 | 3.98E-03 | 8.62E-04 |
| | aligned_probalign | 3.81E-02 | 8.60E-03 | 1.99E-03 | 7.78E-04 |
| | aligned_probcons | 5.72E-02 | 2.19E-02 | 2.99E-03 | 9.64E-04 |
| | aligned_tcoffee | 5.29E-02 | 1.97E-02 | 1.99E-03 | 8.88E-04 |
| | structaligned | 6.52E-02 | 2.77E-02 | 1.99E-03 | 1.01E-03 |
| | correctaligned | 1.61E-01 | 1.69E-01 | 9.95E-04 | 1.83E-03 |
| | integratedaln | 2.06E-01 | 2.12E-01 | 2.35E-01 | 1.11E-03 |
| Insertions | aligned_clustalw | 2.09E-03 | 0.00E+00 | 0.00E+00 | 1.39E-04 |
| | aligned_mafft | 5.10E-03 | 0.00E+00 | 0.00E+00 | 3.40E-04 |
| | aligned_msaprobs | 1.25E-03 | 0.00E+00 | 0.00E+00 | 9.21E-05 |
| | aligned_muscle | 2.28E-02 | 2.00E-04 | 9.88E-04 | 4.98E-04 |
| | aligned_probalign | 1.71E-03 | 0.00E+00 | 0.00E+00 | 1.15E-04 |
| | aligned_probcons | 2.02E-03 | 0.00E+00 | 0.00E+00 | 1.30E-04 |
| | aligned_tcoffee | 4.04E-03 | 0.00E+00 | 0.00E+00 | 2.23E-04 |
| | structaligned | 2.47E-03 | 0.00E+00 | 0.00E+00 | 2.09E-04 |
| | correctaligned | 9.90E-03 | 0.00E+00 | 0.00E+00 | 6.90E-04 |
| | integratedaln | 2.97E-01 | 3.10E-01 | 3.09E-01 | 1.02E-03 |
| Deletions | aligned_clustalw | 4.59E-03 | 0.00E+00 | 0.00E+00 | 2.59E-04 |
| | aligned_mafft | 4.39E-03 | 0.00E+00 | 0.00E+00 | 2.62E-04 |
| | aligned_msaprobs | 3.06E-03 | 0.00E+00 | 0.00E+00 | 2.10E-04 |
| | aligned_muscle | 3.22E-02 | 1.87E-02 | 2.00E-03 | 6.86E-04 |
| | aligned_probalign | 6.98E-03 | 0.00E+00 | 0.00E+00 | 3.23E-04 |
| | aligned_probcons | 5.38E-03 | 0.00E+00 | 0.00E+00 | 2.63E-04 |
| | aligned_tcoffee | 8.14E-03 | 0.00E+00 | 0.00E+00 | 3.48E-04 |
| | structaligned | 4.39E-03 | 0.00E+00 | 0.00E+00 | 4.07E-04 |
| | correctaligned | 8.41E-03 | 0.00E+00 | 0.00E+00 | 9.29E-04 |
| | integratedaln | 4.63E-01 | 4.93E-01 | 5.14E-01 | 2.89E-03 |

**Table S10. Support for the correct ancestral state is increased by alignment-integrated ancestral sequence reconstruction when an incorrect ancestral state is inferred.** We simulated extant and ancestral sequences for 5 protein domain families, using empirically-derived conditions, and aligned the resulting extant sequence data using structure-guided and 7 different sequence-alignment methods (see Methods). We used each alignment to infer the most likely ancestral sequence at each node on the phylogeny. Additionally, alignment-integrated ancestral sequences were generated by combining inferences from the 7 sequence-alignment methods (see Methods). In each case, we compared the inferred ancestral sequences to the correct simulated sequence to estimate ancestral sequence reconstruction (ASR) error rates (see Methods). ASR errors were divided into 4 “errorType” categories: 1) residue errors, in which both correct and inferred ancestral sequences have a residue at a given position in the alignment, but the residues differ; 2) insertion errors, in which the inferred sequence has a residue at a given alignment position, but the correct sequence has a gap state; 3) deletion errors, in which the inferred sequence has a gap, but the correct sequence has a residue, and 4) total errors, which combine residue- insertion- and deletion-errors. For each error type, when an erroneous ancestral state of that error type was inferred as the maximum-likelihood state, we calculated the posterior probability (PP) of the correct state using an empirical-Bayesian approach (see Methods). The distribution of posterior probabilities for the correct ancestral state, when ASR errors of each type were made, was inferred by kernel density estimation. We report the mean (and standard error), median and mode of the posterior-probability distribution of correct ancestral states for each type of ASR error.

| ancState Type | alignment | meanCorrectPP | medianCorrectPP | highestMode | SEmean |
|---|---|---|---|---|---|
| Total | aligned_clustalw | 9.98E-01 | 1.00E+00 | 1.00E+00 | 8.60E-05 |
| | aligned_mafft | 9.97E-01 | 1.00E+00 | 1.00E+00 | 1.05E-04 |
| | aligned_msaprobs | 9.97E-01 | 1.00E+00 | 1.00E+00 | 9.17E-05 |
| | aligned_muscle | 9.95E-01 | 9.99E-01 | 9.99E-01 | 1.05E-04 |
| | aligned_probalign | 9.98E-01 | 1.00E+00 | 1.00E+00 | 8.34E-05 |
| | aligned_probcons | 9.97E-01 | 1.00E+00 | 1.00E+00 | 9.21E-05 |
| | aligned_tcoffee | 9.97E-01 | 1.00E+00 | 1.00E+00 | 9.85E-05 |
| | structaligned | 9.97E-01 | 1.00E+00 | 1.00E+00 | 9.31E-05 |
| | correctaligned | 9.97E-01 | 1.00E+00 | 1.00E+00 | 9.56E-05 |
| | integratedaln | 9.74E-01 | 9.76E-01 | 9.80E-01 | 1.75E-04 |
| Residues | aligned_clustalw | 9.65E-01 | 9.98E-01 | 9.97E-01 | 1.19E-03 |
| | aligned_mafft | 9.63E-01 | 9.98E-01 | 9.97E-01 | 1.25E-03 |
| | aligned_msaprobs | 9.64E-01 | 9.98E-01 | 9.97E-01 | 1.11E-03 |
| | aligned_muscle | 9.48E-01 | 9.94E-01 | 9.95E-01 | 1.31E-03 |
| | aligned_probalign | 9.56E-01 | 9.98E-01 | 9.97E-01 | 1.35E-03 |
| | aligned_probcons | 9.65E-01 | 9.98E-01 | 9.97E-01 | 1.10E-03 |
| | aligned_tcoffee | 9.60E-01 | 9.97E-01 | 9.97E-01 | 1.24E-03 |
| | structaligned | 9.69E-01 | 9.98E-01 | 9.97E-01 | 9.71E-04 |
| | correctaligned | 9.69E-01 | 9.98E-01 | 9.97E-01 | 9.69E-04 |
| | integratedaln | 8.39E-01 | 8.93E-01 | 9.07E-01 | 1.87E-03 |
| Gaps | aligned_clustalw | 1.00E+00 | 1.00E+00 | 1.00E+00 | 2.76E-06 |
| | aligned_mafft | 9.99E-01 | 1.00E+00 | 1.00E+00 | 4.02E-05 |
| | aligned_msaprobs | 1.00E+00 | 1.00E+00 | 1.00E+00 | 3.27E-06 |
| | aligned_muscle | 9.99E-01 | 1.00E+00 | 1.00E+00 | 4.18E-05 |
| | aligned_probalign | 1.00E+00 | 1.00E+00 | 1.00E+00 | 1.31E-05 |
| | aligned_probcons | 1.00E+00 | 1.00E+00 | 1.00E+00 | 9.23E-06 |
| | aligned_tcoffee | 1.00E+00 | 1.00E+00 | 1.00E+00 | 1.60E-05 |
| | structaligned | 1.00E+00 | 1.00E+00 | 1.00E+00 | 8.91E-06 |
| | correctaligned | 1.00E+00 | 1.00E+00 | 1.00E+00 | 6.77E-07 |
| | integratedaln | 9.83E-01 | 9.86E-01 | 9.87E-01 | 1.05E-04 |

**Table S11. Support for the correct ancestral state is reduced by alignment-integrated ancestral sequence reconstruction when the correct state is inferred by maximum-likelihood.** We simulated extant and ancestral sequences for 5 protein domain families, using empirically-derived conditions, and aligned the resulting extant sequence data using structure-guided and 7 different sequence-alignment methods (see Methods). We used each alignment to infer the most likely ancestral sequence at each node on the phylogeny. Additionally, alignment-integrated ancestral sequences were generated by combining inferences from the 7 sequence-alignment methods (see Methods). In each case, we compared the inferred ancestral sequences to the correct simulated sequence to identify ancestral sequence reconstruction (ASR) errors (see Methods). When the correct ancestral state was inferred as the maximum-likelihood state, we calculated the posterior probability (PP) of the correct state using an empirical-Bayesian approach (see Methods). The distribution of posterior probabilities for the correct ancestral state, when no ASR error was made, was inferred by kernel density estimation. We report the mean (and standard error), median and mode of the posterior-probability distribution of correct ancestral states for each type of ancestral state (residues, gaps).

| Property | alignment | meanError | medianError | highestMode | SEmean |
|---|---|---|---|---|---|
| Structural Stability | aligned_clustalw | 3.81E-02 | 3.74E-02 | 3.95E-02 | 3.39E-04 |
| | aligned_mafft | 4.58E-02 | 4.38E-02 | 4.22E-02 | 3.78E-04 |
| | aligned_msaprobs | 2.95E-02 | 2.79E-02 | 2.63E-02 | 2.80E-04 |
| | aligned_muscle | 3.65E-02 | 3.32E-02 | 2.77E-02 | 3.53E-04 |
| | aligned_probalign | 5.12E-02 | 4.92E-02 | 4.33E-02 | 4.01E-04 |
| | aligned_probcons | 2.75E-02 | 2.43E-02 | 2.05E-02 | 3.02E-04 |
| | aligned_tcoffee | 3.34E-02 | 3.18E-02 | 3.12E-02 | 2.70E-04 |
| | structaligned | 2.51E-02 | 2.35E-02 | 2.32E-02 | 2.72E-04 |
| | correctaligned | 1.40E-02 | 1.01E-02 | 7.32E-03 | 3.11E-04 |
| | integratedaln | 2.60E-02 | 2.35E-02 | 2.01E-02 | 2.87E-04 |
| Binding Affinity | aligned_clustalw | 4.58E-01 | 4.58E-01 | 4.73E-01 | 3.22E-03 |
| | aligned_mafft | 4.84E-01 | 4.27E-01 | 4.03E-01 | 5.43E-03 |
| | aligned_msaprobs | 3.26E-01 | 3.05E-01 | 2.73E-01 | 3.41E-03 |
| | aligned_muscle | 4.10E-01 | 3.93E-01 | 3.67E-01 | 2.93E-03 |
| | aligned_probalign | 8.25E-01 | 7.37E-01 | 7.38E-01 | 1.22E-02 |
| | aligned_probcons | 3.55E-01 | 3.48E-01 | 3.50E-01 | 2.69E-03 |
| | aligned_tcoffee | 3.70E-01 | 3.30E-01 | 2.79E-01 | 3.99E-03 |
| | structaligned | 3.46E-01 | 3.30E-01 | 3.03E-01 | 3.03E-03 |
| | correctaligned | 1.70E-01 | 1.42E-01 | 1.08E-01 | 2.62E-03 |
| | integratedaln | 3.13E-01 | 2.99E-01 | 2.61E-01 | 2.70E-03 |

**Table S12. Structure-guided and alignment-integrated methods reduce errors in estimated structural and functional properties of ancestral sequences, compared to sequence-alignment methods.** We simulated replicate extant and ancestral sequence data sets by evolving the DSRM1 protein domain family along its empirically-determined phylogeny, using a structure-guided alignment of empirical DSRM1 sequences to determine the amino-acid composition and pattern of insertions/deletions (see Methods). Structural and functional properties were unconstrained during the simulations. We modeled the structure of each simulated ancestral sequence and estimated its structural stability (ΔG) and dsRNA-binding affinity (pKd) using computational approaches (see Methods). For each alignment method, we inferred the maximum-likelihood ancestral sequence at each node on the phylogeny, modeled the structure of the inferred ancestral sequence at each node, and estimated structural stability and binding affinity from the modeled structure. Structural stability and binding affinity errors were calculated as the absolute value of the difference between the value calculated from the correct simulated ancestral sequence and that calculated from the inferred ancestral sequence. The distributions of structural-stability and binding-affinity errors were calculated by kernel density estimation. We report the mean (and standard error), median and mode of each error distribution.

| Structural Stability | | linear regression | | | | Binding Affinity | | linear regression | | | |
|---|---|---|---|---|---|---|---|---|---|---|---|
| errorType | alignment | slope | intercept | rsquared | SEslope | errorType | alignment | slope | intercept | rsquared | SEslope |
| Total | aligned_clustalw | 0.4149 | 0.0177 | 0.2828 | 5.64E-03 | Total | aligned_clustalw | 1.0682 | 0.406 | 0.0208 | 6.26E-02 |
| | aligned_mafft | 0.2011 | 0.0347 | 0.1135 | 4.80E-03 | | aligned_mafft | 4.599 | 0.2302 | 0.2886 | 6.16E-02 |
| | aligned_msaprobs | 0.4331 | 0.0158 | 0.4482 | 4.10E-03 | | aligned_msaprobs | 5.1293 | 0.1642 | 0.4238 | 5.10E-02 |
| | aligned_muscle | 0.4757 | 0.0152 | 0.4142 | 4.83E-03 | | aligned_muscle | 2.2903 | 0.3071 | 0.1394 | 4.86E-02 |
| | aligned_probalign | 0.1684 | 0.0385 | 0.1771 | 3.10E-03 | | aligned_probalign | 7.0038 | 0.2962 | 0.3337 | 8.44E-02 |
| | aligned_probcons | 0.5531 | 0.0122 | 0.5207 | 4.53E-03 | | aligned_probcons | 3.4615 | 0.2597 | 0.2562 | 5.03E-02 |
| | aligned_tcoffee | 0.3193 | 0.019 | 0.2203 | 5.13E-03 | | aligned_tcoffee | 5.1166 | 0.1383 | 0.2581 | 7.40E-02 |
| | structaligned | 0.5173 | 0.0136 | 0.4956 | 4.45E-03 | | structaligned | 4.0694 | 0.2558 | 0.2471 | 6.06E-02 |
| | correctaligned | 0.7388 | 0.0085 | 0.7793 | 3.35E-03 | | correctaligned | 5.6261 | 0.1274 | 0.6371 | 3.62E-02 |
| | integratedaln | 0.6029 | 0.0128 | 0.5495 | 4.66E-03 | | integratedaln | 4.1519 | 0.2221 | 0.2935 | 5.50E-02 |
| Residues | aligned_clustalw | 0.4498 | 0.0263 | 0.2596 | 6.48E-03 | Residues | aligned_clustalw | 1.3831 | 0.4223 | 0.0272 | 7.06E-02 |
| | aligned_mafft | 0.4367 | 0.0363 | 0.1723 | 8.17E-03 | | aligned_mafft | 7.2244 | 0.3271 | 0.2293 | 1.13E-01 |
| | aligned_msaprobs | 0.5147 | 0.0212 | 0.4089 | 5.28E-03 | | aligned_msaprobs | 5.3511 | 0.2396 | 0.2979 | 7.01E-02 |
| | aligned_muscle | 0.5907 | 0.0236 | 0.3928 | 6.27E-03 | | aligned_muscle | 2.528 | 0.3542 | 0.1045 | 6.32E-02 |
| | aligned_probalign | 0.32 | 0.0421 | 0.1559 | 6.35E-03 | | aligned_probalign | 13.6434 | 0.4396 | 0.3085 | 1.74E-01 |
| | aligned_probcons | 0.5895 | 0.0189 | 0.4579 | 5.47E-03 | | aligned_probcons | 3.6324 | 0.302 | 0.2184 | 5.86E-02 |
| | aligned_tcoffee | 0.4168 | 0.0276 | 0.2906 | 5.56E-03 | | aligned_tcoffee | 4.0606 | 0.3138 | 0.1259 | 9.13E-02 |
| | structaligned | 0.5626 | 0.019 | 0.526 | 4.56E-03 | | structaligned | 3.9771 | 0.3029 | 0.2117 | 6.55E-02 |
| | correctaligned | 0.74 | 0.0085 | 0.7779 | 3.37E-03 | | correctaligned | 5.6492 | 0.1278 | 0.6391 | 3.62E-02 |
| | integratedaln | 0.6054 | 0.018 | 0.5111 | 5.05E-03 | | integratedaln | 3.8968 | 0.2615 | 0.2386 | 5.94E-02 |
| Insertions | aligned_clustalw | 3.5195 | -0.002 | 0.1648 | 6.76E-02 | Insertions | aligned_clustalw | -3.4471 | 0.4977 | 0.0018 | 7.02E-01 |
| | aligned_mafft | 0.5275 | 0.0368 | 0.0612 | 1.76E-02 | | aligned_mafft | 15.9376 | 0.2128 | 0.2716 | 2.23E-01 |
| | aligned_msaprobs | 1.7365 | 0.0161 | 0.2098 | 2.88E-02 | | aligned_msaprobs | 28.7101 | 0.1048 | 0.3867 | 3.09E-01 |
| | aligned_muscle | 1.6341 | 0.0174 | 0.1822 | 2.95E-02 | | aligned_muscle | 11.2059 | 0.2785 | 0.1244 | 2.54E-01 |
| | aligned_probalign | 0.6086 | 0.0368 | 0.174 | 1.13E-02 | | aligned_probalign | 24.651 | 0.2413 | 0.3109 | 3.13E-01 |
| | aligned_probcons | 3.0809 | 0.0075 | 0.2543 | 4.50E-02 | | aligned_probcons | 20.7379 | 0.2208 | 0.1447 | 4.30E-01 |
| | aligned_tcoffee | 0.1163 | 0.0316 | 0.001 | 3.21E-02 | | aligned_tcoffee | 31.1141 | -0.1216 | 0.3119 | 3.94E-01 |
| | structaligned | 0.8825 | 0.0201 | 0.0072 | 8.87E-02 | | structaligned | 48.5075 | 0.0716 | 0.174 | 9.02E-01 |
| | correctaligned | 15.692 | 0.0132 | 0.029 | 7.75E-01 | | correctaligned | 55.4388 | 0.1667 | 0.0051 | 6.61E+00 |
| | integratedaln | 3.5127 | 0.0187 | 0.0639 | 1.15E-01 | | integratedaln | 33.1239 | 0.244 | 0.064 | 1.08E+00 |
| Deletions | aligned_clustalw | 4.5652 | -0.0144 | 0.2634 | 6.51E-02 | Deletions | aligned_clustalw | 2.5011 | 0.4297 | 0.0009 | 7.20E-01 |
| | aligned_mafft | 0.3128 | 0.0407 | 0.0168 | 2.04E-02 | | aligned_mafft | 16.7955 | 0.2078 | 0.2348 | 2.59E-01 |
| | aligned_msaprobs | 1.7272 | 0.0162 | 0.2066 | 2.89E-02 | | aligned_msaprobs | 28.7887 | 0.1044 | 0.387 | 3.09E-01 |
| | aligned_muscle | 2.1685 | 0.0123 | 0.2041 | 3.65E-02 | | aligned_muscle | 13.372 | 0.2601 | 0.1127 | 3.20E-01 |
| | aligned_probalign | 0.6004 | 0.037 | 0.1676 | 1.14E-02 | | aligned_probalign | 24.5637 | 0.2459 | 0.3055 | 3.16E-01 |
| | aligned_probcons | 3.1011 | 0.0074 | 0.2542 | 4.53E-02 | | aligned_probcons | 20.6997 | 0.2212 | 0.1423 | 4.34E-01 |
| | aligned_tcoffee | -0.2079 | 0.0367 | 0.0032 | 3.11E-02 | | aligned_tcoffee | 27.4858 | -0.0589 | 0.2592 | 3.96E-01 |
| | structaligned | 1.5557 | 0.0162 | 0.0227 | 8.71E-02 | | structaligned | 50.9999 | 0.0561 | 0.1963 | 8.80E-01 |
| | correctaligned | 13.362 | 0.0134 | 0.0254 | 7.06E-01 | | correctaligned | 18.2189 | 0.1687 | 0.0007 | 6.02E+00 |
| | integratedaln | 2.285 | 0.0109 | 0.111 | 5.52E-02 | | integratedaln | 31.1916 | 0.1065 | 0.2329 | 4.83E-01 |

**Table S13. Errors in estimated structural stability and binding affinity of ancestral sequences are weakly correlated with errors in the inferred ancestral sequences.** We simulated replicate extant and ancestral sequence data sets by evolving the DSRM1 protein domain family along its empirically-determined phylogeny, using a structure-guided alignment of empirical DSRM1 sequences to determine the amino-acid composition and pattern of insertions/ deletions (see Methods). Structural and functional properties were unconstrained during the simulations. We modeled the structure of each simulated ancestral sequence and estimated its structural stability (ΔG) and dsRNA-binding affinity (pKd) using computational approaches (see Methods). For each alignment method, we inferred the maximum-likelihood ancestral sequence at each node on the phylogeny, modeled the structure of the inferred ancestral sequence at each node, and estimated structural stability and binding affinity from the modeled structure. Structural stability and binding affinity errors were calculated as the absolute value of the difference between the value calculated from the correct simulated ancestral sequence and that calculated from the inferred ancestral sequence. ASR errors were divided into 4 “errorType” categories: 1) residue errors, in which both correct and inferred ancestral sequences have a residue at a given position in the alignment, but the residues differ; 2) insertion errors, in which the inferred sequence has a residue at a given alignment position, but the correct sequence has a gap state; 3) deletion errors, in which the inferred sequence has a gap, but the correct sequence has a residue, and 4) total errors, which combine residue- insertion- and deletion-errors. Sequence-wide error rates (errors/site) were computed by dividing the number of errors by the length of the pairwise alignment of the inferred and correct ancestral sequences. For each type of ASR error and alignment method, we determined the best-fit linear relationship between ASR error rate and structural stability (left) or binding affinity (right) using ordinary least squares linear regression. We report the slope (and standard error), intercept and correlation (rsquared) of each linear regression.

| IndelRate | Alignment | Slope | SE(Slope) | P(Slope>0) | Intercept | $r^2$ |
|---|---|---|---|---|---|---|
| 0.01 | aligned_clustalw | 0.5008 | 4.29E-02 | 2.38E-05 | 0.08 | 0.9578 |
| | aligned_mafft | 0.4825 | 4.15E-02 | 2.44E-05 | 0.0802 | 0.9575 |
| | aligned_msaprobs | 0.4871 | 4.18E-02 | 2.41E-05 | 0.0816 | 0.9577 |
| | aligned_muscle | 0.4876 | 4.38E-02 | 3.12E-05 | 0.0813 | 0.9539 |
| | aligned_probalign | 0.4941 | 4.16E-02 | 2.17E-05 | 0.0799 | 0.9591 |
| | aligned_probcons | 0.4866 | 4.31E-02 | 2.87E-05 | 0.081 | 0.9551 |
| | aligned_tcoffee | 0.4921 | 4.28E-02 | 2.59E-05 | 0.0804 | 0.9567 |
| | correctaligned | 0.4725 | 4.47E-02 | 4.19E-05 | 0.0819 | 0.9491 |
| | integratedaln | 0.4623 | 4.46E-02 | 4.69E-05 | 0.0847 | 0.9472 |
| 0.02 | aligned_clustalw | 0.4992 | 5.02E-02 | 5.97E-05 | 0.0882 | 0.9428 |
| | aligned_mafft | 0.4779 | 4.74E-02 | 5.49E-05 | 0.0861 | 0.9444 |
| | aligned_msaprobs | 0.4838 | 4.78E-02 | 5.43E-05 | 0.0862 | 0.9446 |
| | aligned_muscle | 0.4916 | 4.73E-02 | 4.67E-05 | 0.0825 | 0.9473 |
| | aligned_probalign | 0.4944 | 4.75E-02 | 4.63E-05 | 0.0851 | 0.9474 |
| | aligned_probcons | 0.4877 | 4.84E-02 | 5.54E-05 | 0.0847 | 0.9442 |
| | aligned_tcoffee | 0.4922 | 4.92E-02 | 5.76E-05 | 0.0858 | 0.9435 |
| | correctaligned | 0.4643 | 4.83E-02 | 7.28E-05 | 0.0876 | 0.9389 |
| | integratedaln | 0.4524 | 4.92E-02 | 9.36E-05 | 0.0888 | 0.9337 |
| 0.03 | aligned_clustalw | 0.5046 | 5.35E-02 | 8.09E-05 | 0.0926 | 0.9368 |
| | aligned_mafft | 0.4744 | 4.71E-02 | 5.53E-05 | 0.0881 | 0.9443 |
| | aligned_msaprobs | 0.4846 | 4.79E-02 | 5.43E-05 | 0.0884 | 0.9446 |
| | aligned_muscle | 0.492 | 5.00E-02 | 6.32E-05 | 0.0872 | 0.9417 |
| | aligned_probalign | 0.4894 | 4.81E-02 | 5.26E-05 | 0.0894 | 0.9452 |
| | aligned_probcons | 0.4873 | 4.69E-02 | 4.63E-05 | 0.0868 | 0.9474 |
| | aligned_tcoffee | 0.4942 | 4.71E-02 | 4.38E-05 | 0.0874 | 0.9484 |
| | correctaligned | 0.4587 | 4.99E-02 | 9.29E-05 | 0.0905 | 0.9338 |
| | integratedaln | 0.4472 | 4.82E-02 | 8.86E-05 | 0.0914 | 0.9348 |
| 0.04 | aligned_clustalw | 0.483 | 6.39E-02 | 2.79E-04 | 0.1068 | 0.9048 |
| | aligned_mafft | 0.4556 | 5.51E-02 | 1.70E-04 | 0.0978 | 0.9192 |
| | aligned_msaprobs | 0.4564 | 5.77E-02 | 2.18E-04 | 0.1028 | 0.9124 |
| | aligned_muscle | 0.4765 | 5.86E-02 | 1.86E-04 | 0.0977 | 0.9167 |
| | aligned_probalign | 0.4703 | 5.72E-02 | 1.75E-04 | 0.1022 | 0.9185 |
| | aligned_probcons | 0.4655 | 5.62E-02 | 1.68E-04 | 0.0989 | 0.9195 |
| | aligned_tcoffee | 0.4696 | 5.84E-02 | 1.99E-04 | 0.1021 | 0.915 |
| | correctaligned | 0.4398 | 5.71E-02 | 2.52E-04 | 0.0999 | 0.9081 |
| | integratedaln | 0.4187 | 5.59E-02 | 2.92E-04 | 0.1022 | 0.9035 |
| 0.05 | aligned_clustalw | 0.459 | 6.68E-02 | 4.68E-04 | 0.1228 | 0.8873 |
| | aligned_mafft | 0.4437 | 5.47E-02 | 1.89E-04 | 0.1057 | 0.9164 |
| | aligned_msaprobs | 0.4425 | 5.54E-02 | 2.05E-04 | 0.1121 | 0.914 |
| | aligned_muscle | 0.473 | 5.43E-02 | 1.27E-04 | 0.1047 | 0.9266 |
| | aligned_probalign | 0.4535 | 5.69E-02 | 2.07E-04 | 0.1141 | 0.9138 |
| | aligned_probcons | 0.4523 | 5.37E-02 | 1.53E-04 | 0.1076 | 0.922 |

| | | | | | | |
|---|---|---|---|---|---|---|
| | aligned_tcoffee | 0.4592 | 5.69E-02 | 1.94E-04 | 0.1108 | 0.9156 |
| | correctaligned | 0.4268 | 5.70E-02 | 2.92E-04 | 0.1072 | 0.9034 |
| | integratedaln | 0.4086 | 5.35E-02 | 2.63E-04 | 0.1098 | 0.9067 |
| 0.06 | aligned_clustalw | 0.4581 | 7.16E-02 | 6.86E-04 | 0.1286 | 0.8722 |
| | aligned_mafft | 0.437 | 5.66E-02 | 2.47E-04 | 0.1142 | 0.9086 |
| | aligned_msaprobs | 0.4418 | 5.88E-02 | 2.88E-04 | 0.117 | 0.9038 |
| | aligned_muscle | 0.4884 | 5.56E-02 | 1.21E-04 | 0.1045 | 0.9278 |
| | aligned_probalign | 0.4446 | 6.01E-02 | 3.15E-04 | 0.1211 | 0.9011 |
| | aligned_probcons | 0.4485 | 5.75E-02 | 2.33E-04 | 0.1138 | 0.9103 |
| | aligned_tcoffee | 0.4515 | 6.20E-02 | 3.41E-04 | 0.1185 | 0.8984 |
| | correctaligned | 0.4252 | 5.63E-02 | 2.79E-04 | 0.1123 | 0.9049 |
| | integratedaln | 0.4044 | 5.52E-02 | 3.30E-04 | 0.1144 | 0.8995 |
| 0.07 | aligned_clustalw | 0.4409 | 7.53E-02 | 1.09E-03 | 0.139 | 0.8512 |
| | aligned_mafft | 0.4233 | 6.35E-02 | 5.50E-04 | 0.1234 | 0.8811 |
| | aligned_msaprobs | 0.4183 | 6.35E-02 | 5.90E-04 | 0.1281 | 0.8784 |
| | aligned_muscle | 0.4777 | 5.82E-02 | 1.77E-04 | 0.1127 | 0.9182 |
| | aligned_probalign | 0.4246 | 6.66E-02 | 7.00E-04 | 0.1328 | 0.8713 |
| | aligned_probcons | 0.4275 | 6.29E-02 | 4.96E-04 | 0.1247 | 0.8851 |
| | aligned_tcoffee | 0.4327 | 6.42E-02 | 5.19E-04 | 0.1283 | 0.8834 |
| | correctaligned | 0.3991 | 6.47E-02 | 8.32E-04 | 0.1253 | 0.8639 |
| | integratedaln | 0.3793 | 6.03E-02 | 7.53E-04 | 0.1242 | 0.8682 |
| 0.08 | aligned_clustalw | 0.435 | 7.33E-02 | 1.02E-03 | 0.1444 | 0.8543 |
| | aligned_mafft | 0.4398 | 6.10E-02 | 3.62E-04 | 0.119 | 0.8964 |
| | aligned_msaprobs | 0.4316 | 5.80E-02 | 3.04E-04 | 0.1267 | 0.9021 |
| | aligned_muscle | 0.4947 | 5.80E-02 | 1.43E-04 | 0.1135 | 0.9237 |
| | aligned_probalign | 0.4309 | 6.14E-02 | 4.17E-04 | 0.1321 | 0.8914 |
| | aligned_probcons | 0.4331 | 5.88E-02 | 3.21E-04 | 0.1257 | 0.9004 |
| | aligned_tcoffee | 0.4416 | 5.78E-02 | 2.63E-04 | 0.1291 | 0.9068 |
| | correctaligned | 0.4031 | 6.27E-02 | 6.71E-04 | 0.125 | 0.8732 |
| | integratedaln | 0.3924 | 5.59E-02 | 4.18E-04 | 0.1216 | 0.8914 |

| BranchLength | Alignment | Slope | SE(Slope) | P(Slope>0) | Intercept | r2 |
|---|---|---|---|---|---|---|
| 0.1 | aligned_clustalw | 0.4271 | 3.29E-02 | 1.29E-05 | 0.0858 | 0.9656 |
| | aligned_mafft | 0.2894 | 2.87E-02 | 5.49E-05 | 0.086 | 0.9444 |
| | aligned_msaprobs | 0.327 | 4.07E-02 | 1.98E-04 | 0.087 | 0.915 |
| | aligned_muscle | 0.2825 | 2.82E-02 | 5.70E-05 | 0.0865 | 0.9437 |
| | aligned_probalign | 0.3726 | 3.55E-02 | 4.38E-05 | 0.0861 | 0.9484 |
| | aligned_probcons | 0.3268 | 3.59E-02 | 9.86E-05 | 0.0857 | 0.9325 |
| | aligned_tcoffee | 0.3725 | 4.02E-02 | 8.93E-05 | 0.0852 | 0.9347 |
| | correctaligned | 0.2727 | 3.13E-02 | 1.27E-04 | 0.0858 | 0.9267 |
| | integratedaln | 0.2656 | 3.11E-02 | 1.41E-04 | 0.0863 | 0.924 |
| 0.2 | aligned_clustalw | 0.7636 | 4.91E-02 | 4.48E-06 | 0.1666 | 0.9758 |
| | aligned_mafft | 0.5093 | 8.42E-02 | 9.25E-04 | 0.1659 | 0.8591 |
| | aligned_msaprobs | 0.6071 | 5.46E-02 | 3.15E-05 | 0.1654 | 0.9537 |
| | aligned_muscle | 0.5685 | 5.23E-02 | 3.60E-05 | 0.1646 | 0.9516 |
| | aligned_probalign | 0.7061 | 6.78E-02 | 4.59E-05 | 0.164 | 0.9476 |
| | aligned_probcons | 0.6021 | 6.19E-02 | 6.75E-05 | 0.1639 | 0.9405 |
| | aligned_tcoffee | 0.6739 | 4.89E-02 | 9.11E-06 | 0.1644 | 0.9693 |
| | correctaligned | 0.4964 | 6.20E-02 | 2.02E-04 | 0.1642 | 0.9145 |
| | integratedaln | 0.4562 | 6.51E-02 | 4.21E-04 | 0.1645 | 0.8911 |
| 0.3 | aligned_clustalw | 0.8162 | 8.92E-02 | 9.61E-05 | 0.2429 | 0.9331 |
| | aligned_mafft | 0.43 | 6.73E-02 | 6.90E-04 | 0.2401 | 0.8719 |
| | aligned_msaprobs | 0.5168 | 6.85E-02 | 2.80E-04 | 0.2423 | 0.9047 |
| | aligned_muscle | 0.62 | 5.88E-02 | 4.30E-05 | 0.2381 | 0.9487 |
| | aligned_probalign | 0.6067 | 8.12E-02 | 2.97E-04 | 0.2433 | 0.903 |
| | aligned_probcons | 0.4725 | 7.27E-02 | 6.30E-04 | 0.2427 | 0.8757 |
| | aligned_tcoffee | 0.579 | 7.07E-02 | 1.79E-04 | 0.2428 | 0.9178 |
| | correctaligned | 0.3969 | 5.67E-02 | 4.24E-04 | 0.2388 | 0.8909 |
| | integratedaln | 0.2893 | 5.52E-02 | 1.94E-03 | 0.2408 | 0.8206 |
| 0.4 | aligned_clustalw | 0.846 | 1.13E-01 | 2.97E-04 | 0.3028 | 0.9029 |

| | | | | | | |
|---|---|---|---|---|---|---|
| | aligned_mafft | 0.5282 | 1.17E-01 | 4.09E-03 | 0.2941 | 0.7717 |
| | aligned_msaprobs | 0.4907 | 8.84E-02 | 1.44E-03 | 0.299 | 0.8372 |
| | aligned_muscle | 0.5299 | 4.03E-02 | 1.19E-05 | 0.2996 | 0.9665 |
| | aligned_probalign | 0.585 | 9.01E-02 | 6.36E-04 | 0.3 | 0.8754 |
| | aligned_probcons | 0.5427 | 7.56E-02 | 3.70E-04 | 0.2973 | 0.8956 |
| | aligned_tcoffee | 0.5755 | 9.60E-02 | 9.71E-04 | 0.3 | 0.8568 |
| | correctaligned | 0.4323 | 1.02E-01 | 5.35E-03 | 0.2933 | 0.7511 |
| | integratedaln | 0.2458 | 6.79E-02 | 1.11E-02 | 0.2957 | 0.6859 |
| 0.5 | aligned_clustalw | 0.8567 | 7.68E-02 | 3.11E-05 | 0.3483 | 0.9539 |
| | aligned_mafft | 0.4829 | 6.94E-02 | 4.39E-04 | 0.3379 | 0.8896 |
| | aligned_msaprobs | 0.4794 | 4.89E-02 | 6.49E-05 | 0.3424 | 0.9412 |
| | aligned_muscle | 0.7386 | 7.79E-02 | 7.82E-05 | 0.3386 | 0.9375 |
| | aligned_probalign | 0.5631 | 5.16E-02 | 3.53E-05 | 0.3436 | 0.952 |
| | aligned_probcons | 0.4999 | 3.92E-02 | 1.42E-05 | 0.3413 | 0.9644 |
| | aligned_tcoffee | 0.5512 | 5.06E-02 | 3.54E-05 | 0.3453 | 0.9519 |
| | correctaligned | 0.3511 | 6.41E-02 | 1.55E-03 | 0.3371 | 0.8334 |
| | integratedaln | 0.1962 | 5.49E-02 | 1.17E-02 | 0.3352 | 0.6807 |
| 0.6 | aligned_clustalw | 0.4164 | 8.01E-02 | 2.02E-03 | 0.3961 | 0.8183 |
| | aligned_mafft | 0.2517 | 5.63E-02 | 4.24E-03 | 0.3797 | 0.769 |
| | aligned_msaprobs | 0.1785 | 5.81E-02 | 2.20E-02 | 0.3863 | 0.6109 |
| | aligned_muscle | 0.4879 | 5.89E-02 | 1.67E-04 | 0.3848 | 0.9196 |
| | aligned_probalign | 0.2488 | 5.85E-02 | 5.36E-03 | 0.3893 | 0.751 |
| | aligned_probcons | 0.1417 | 6.27E-02 | 6.45E-02 | 0.3881 | 0.46 |
| | aligned_tcoffee | 0.2664 | 3.89E-02 | 4.77E-04 | 0.3901 | 0.8866 |
| | correctaligned | 0.0923 | 6.26E-02 | 1.91E-01 | 0.3802 | 0.266 |
| | integratedaln | -0.0495 | 5.64E-02 | 4.14E-01 | 0.3748 | 0.1138 |
| 0.7 | aligned_clustalw | 0.0237 | 4.89E-02 | 6.46E-01 | 0.4347 | 0.0376 |
| | aligned_mafft | -0.1107 | 9.98E-02 | 3.10E-01 | 0.4162 | 0.1701 |
| | aligned_msaprobs | -0.191 | 8.84E-02 | 7.41E-02 | 0.4261 | 0.4375 |
| | aligned_muscle | 0.3887 | 8.70E-02 | 4.24E-03 | 0.4146 | 0.7691 |
| | aligned_probalign | -0.1379 | 4.69E-02 | 2.59E-02 | 0.4302 | 0.5905 |
| | aligned_probcons | -0.1813 | 5.41E-02 | 1.54E-02 | 0.4265 | 0.6521 |
| | aligned_tcoffee | -0.0925 | 6.88E-02 | 2.28E-01 | 0.4278 | 0.2314 |
| | correctaligned | -0.3151 | 7.82E-02 | 6.86E-03 | 0.415 | 0.7304 |
| | integratedaln | -0.403 | 7.76E-02 | 2.03E-03 | 0.4103 | 0.8179 |
| 0.8 | aligned_clustalw | -0.1818 | 7.08E-02 | 4.25E-02 | 0.4481 | 0.5234 |
| | aligned_mafft | -0.1796 | 4.45E-02 | 6.79E-03 | 0.431 | 0.7313 |
| | aligned_msaprobs | -0.1835 | 9.13E-02 | 9.11E-02 | 0.435 | 0.4025 |
| | aligned_muscle | 0.3887 | 9.04E-02 | 5.10E-03 | 0.4239 | 0.7549 |
| | aligned_probalign | -0.2145 | 8.54E-02 | 4.57E-02 | 0.443 | 0.5127 |
| | aligned_probcons | -0.1157 | 8.34E-02 | 2.15E-01 | 0.4345 | 0.2429 |
| | aligned_tcoffee | -0.0704 | 1.04E-01 | 5.23E-01 | 0.4382 | 0.0712 |
| | correctaligned | -0.3271 | 6.47E-02 | 2.32E-03 | 0.4226 | 0.8099 |
| | integratedaln | -0.3873 | 9.44E-02 | 6.34E-03 | 0.4163 | 0.7372 |

**Table S14. Ancestral sequence reconstruction error rates were positively correlated with increasing branch lengths and weakly correlated with increasing insertion-deletion rates in 3-taxon simulations.** We simulated protein sequences along a 3-taxon phylogeny with equal branch lengths and the same insertion-deletion (indel) rate across the phylogeny (see Methods, Supplementary Fig. S12). The resulting extant sequence data was aligned using 7 different sequence-alignment methods (see Methods). We used each alignment to infer the most likely ancestral sequence at the single node on the 3-taxon phylogeny. Additionally, alignment-integrated ancestral sequences were generated by combining inferences from the 7 sequence-alignment methods (see Methods). In each case, we compared the inferred ancestral sequence to the correct simulated sequence to estimate ancestral sequence reconstruction (ASR) error rates (see Methods). Sequence-wide error rates (errors/site) were computed by dividing the total number of errors by the length of the pairwise alignment of the inferred and correct ancestral sequences. Top table reports results from linear-regression of indel rate vs ASR error rate, for each branch length and alignment method. Bottom table reports results from linear-regression of branch length vs ASR error rate, for each indel rate and alignment method. In each case, we report the slope (and standard error) of the best-fit regression line, the *p*-value obtained by testing if the slope is significantly different from zero, the intercept of the best-fit regression line, and the correlation ($r^2$) between the predictive (branch length or indel rate) and response (ASR error rate) variable.

#### Supplementary Figures

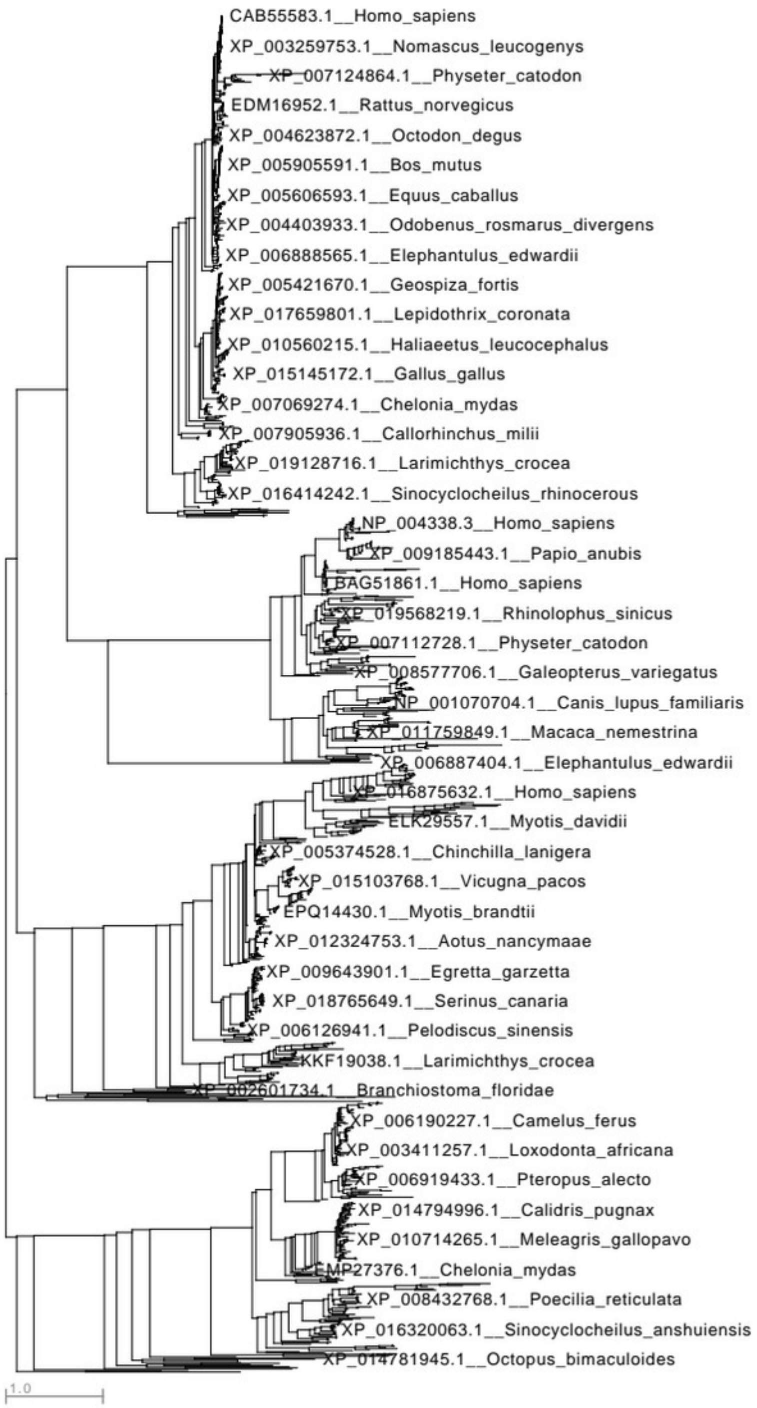


**Figure S1 Part 1, CARD domain. Empirical protein domain family phylogenies used to simulate sequence data.** We simulated protein sequence data along the indicated empirical phylogenies. Branch lengths are scaled to the expected number of substitutions/site. Part 1 shows the empirical phylogeny of the CARD domain family; other domain family phylogenies are shown in parts 2-5, below.

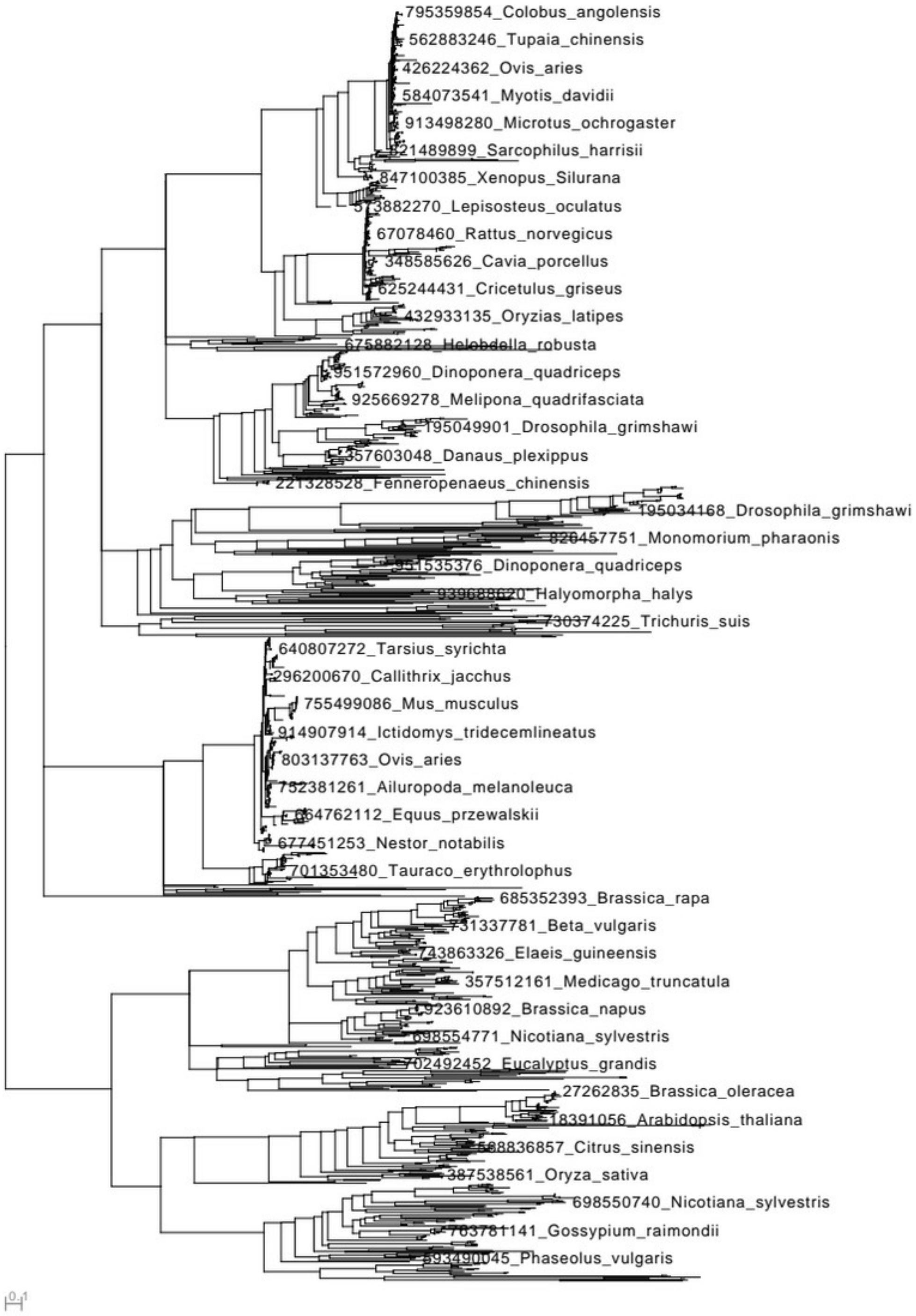


**Figure S1 Part 2, DSRM1 domain.**

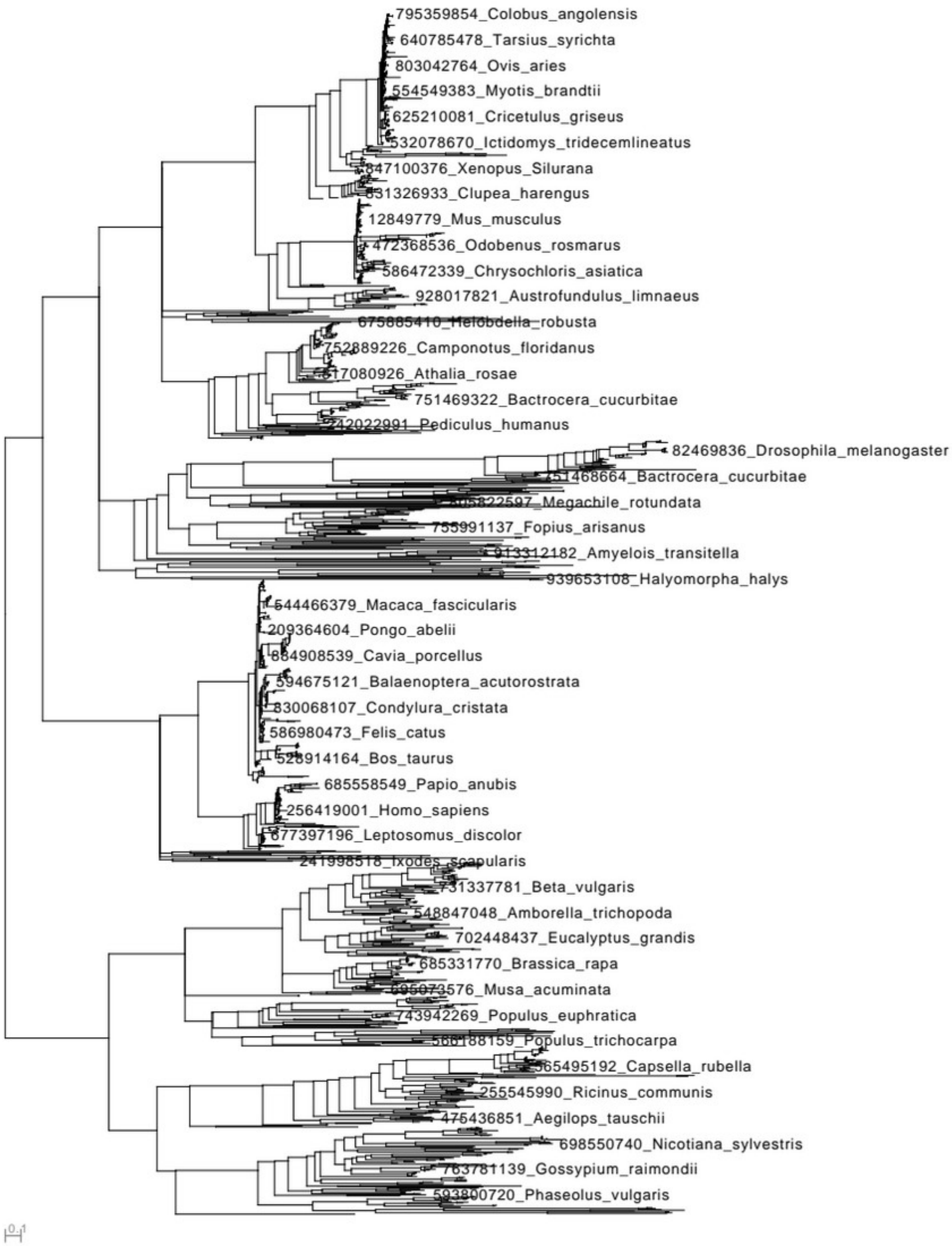


**Figure S1 Part 3, DSRM2 domain.**

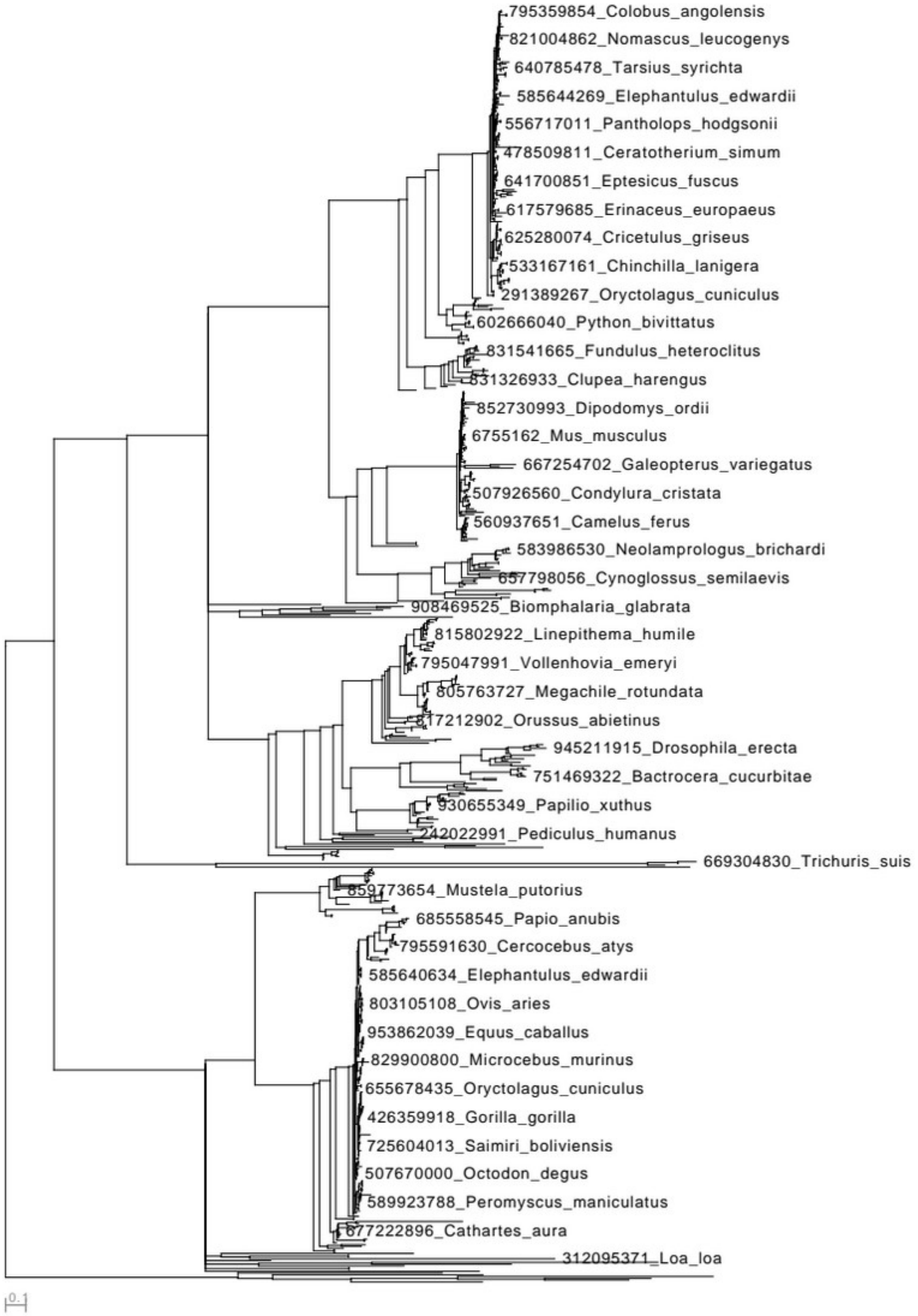


**Figure S1 Part 4, DSRM3 domain.**

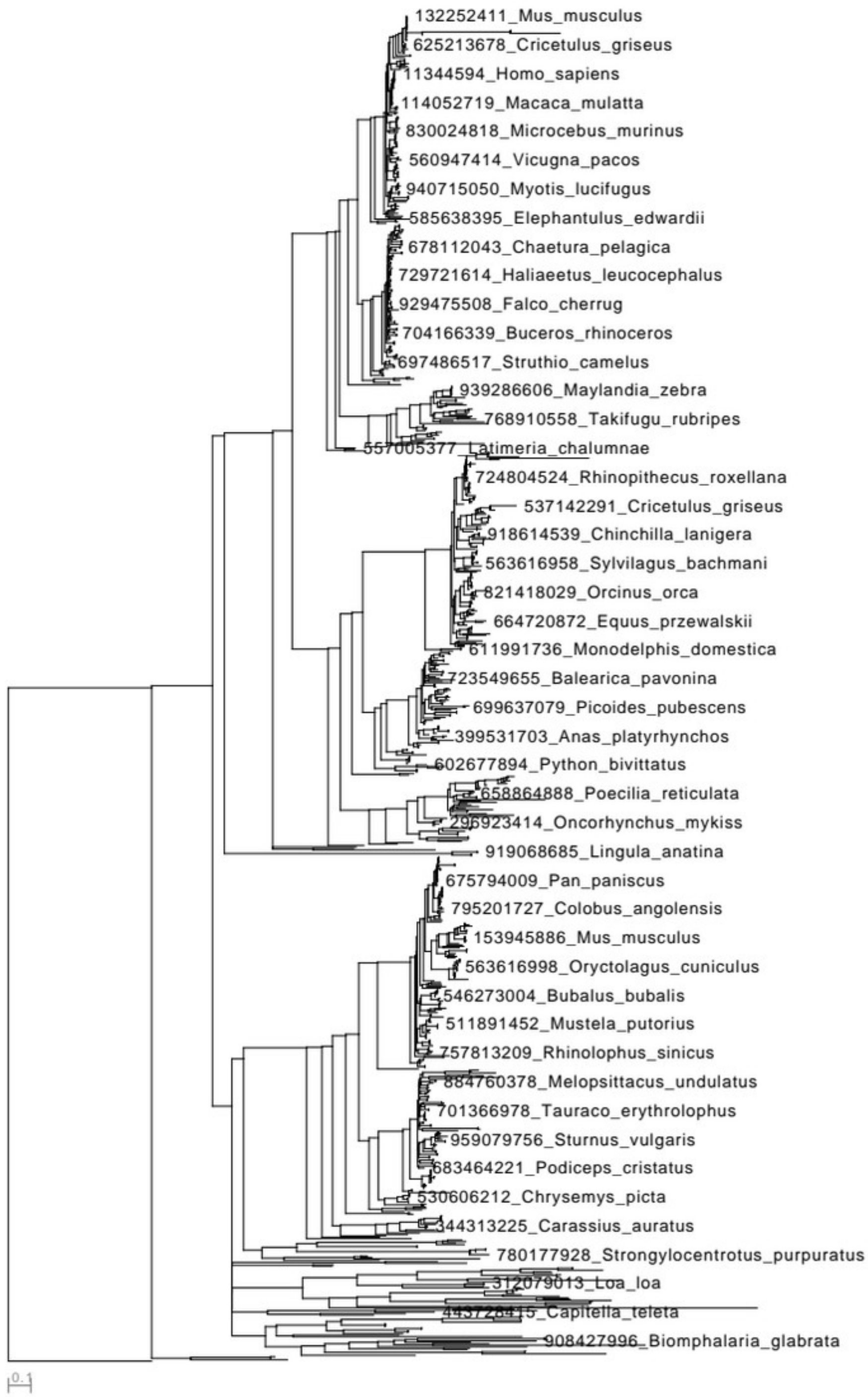


**Figure S1 Part 5, RD domain.**

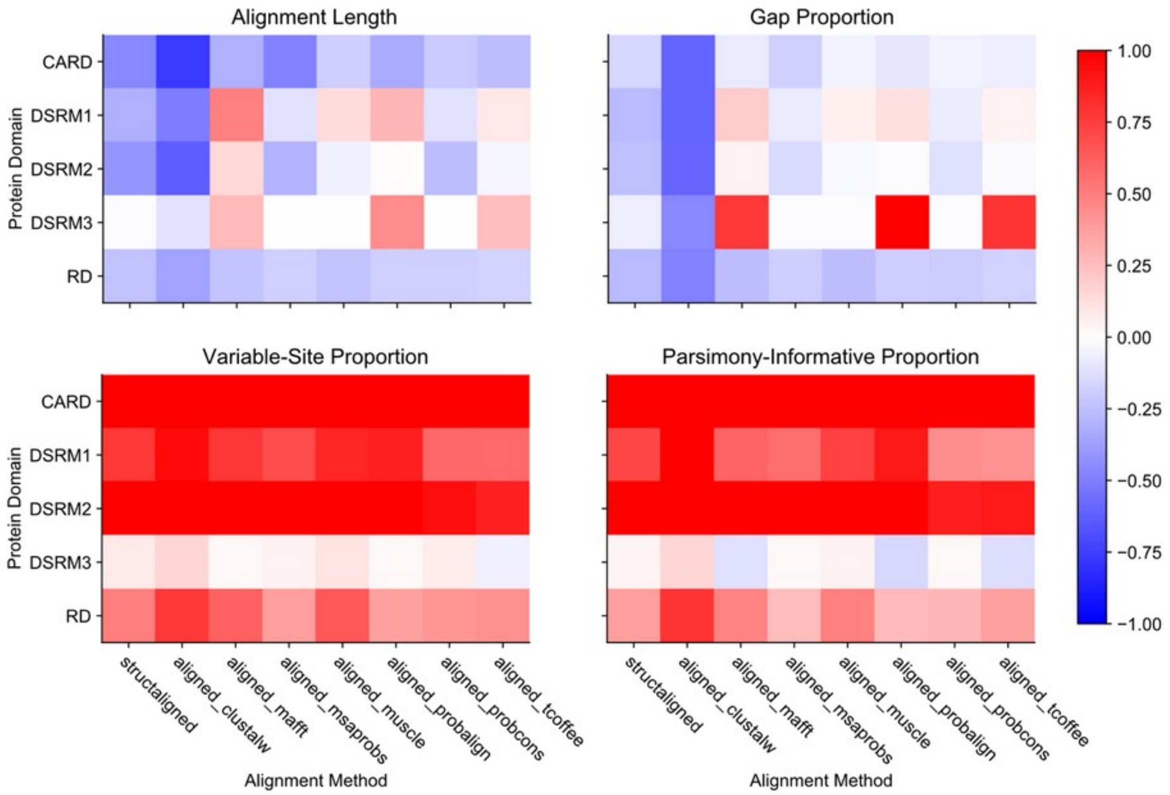


**Figure S2. Structure-guided and sequence-alignment methods typically underestimate correct alignment lengths and gap proportions while overestimating the proportions of variable and parsimony-informative sites.** We simulated alignments of 5 protein domain families, using empirically-derived simulation conditions, and aligned the resulting replicate sequence data using structure-guided and 7 different sequence-alignment methods (see Methods). Colors indicate the proportion of the correct-alignment's value by which the inferred value deviates from the correct-alignment's value, with negative numbers (blue) indicating underestimation of the correct-alignment's value, and positive numbers (red) indicating overestimation of the correct-alignment's value. A value of 1 (-1) indicates over- (under-) estimation of the correct value by 50%.

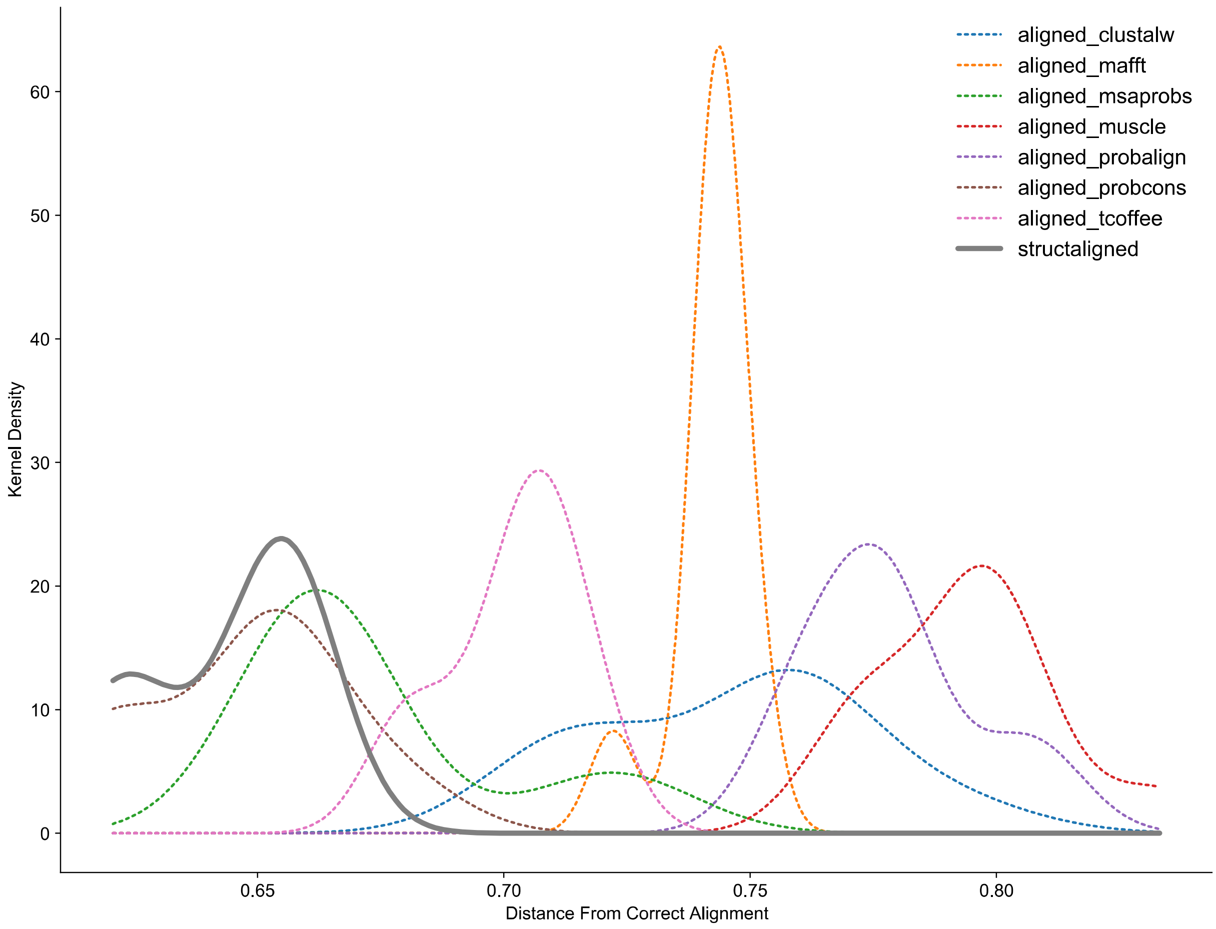


**Figure S3 Part 1, CARD Domain. Alignment distance distributions differ across alignment methods and protein domain families.** We simulated alignments of 5 protein domain families, using empirically-derived simulation conditions, and aligned the resulting replicate sequence data using structure-guided and 7 different sequence-alignment methods (see Methods). For each sequence alignment, we measured the position-wise distance of that alignment to the correct simulated alignment, which estimates the probability of randomly selecting an incorrectly-aligned residue from the alignment (see Methods). We estimated the distribution of alignment distances across replicate simulations using kernel density estimation. Part 1 shows results for the CARD domain family; results for other domain families are shown in parts 2-5, below.

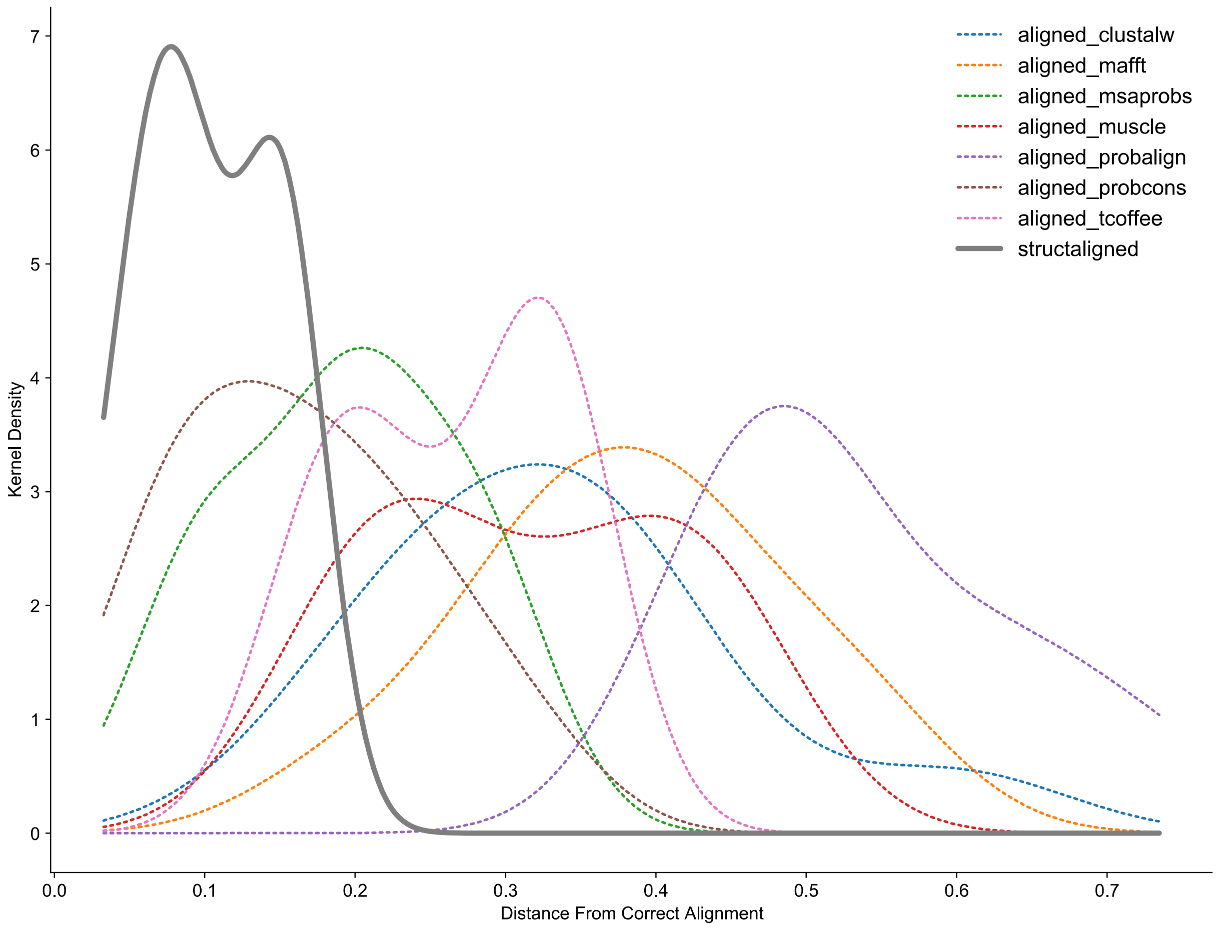


**Figure S3 Part 2, DSRM1 Domain.**

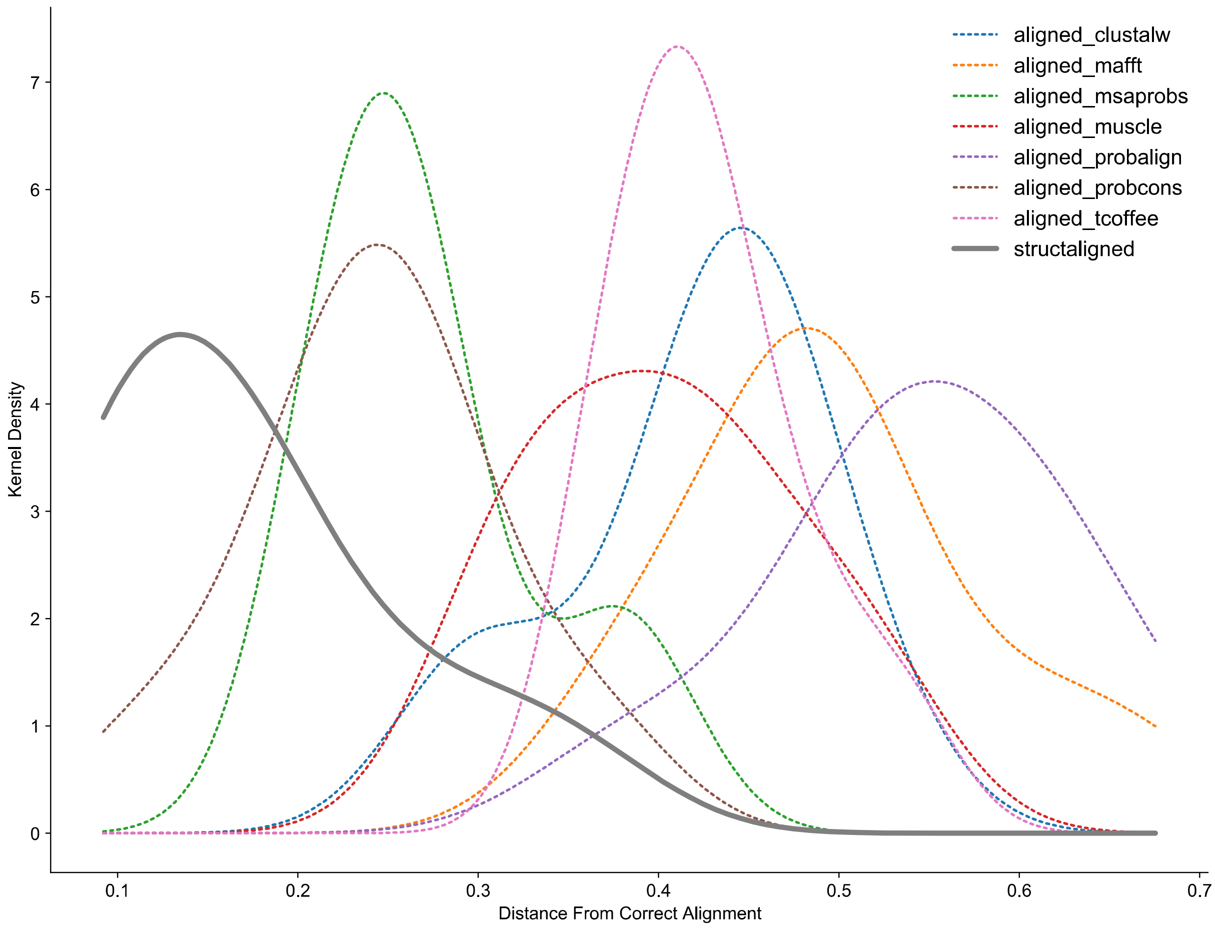


**Figure S3 Part 3, DSRM2 Domain.**

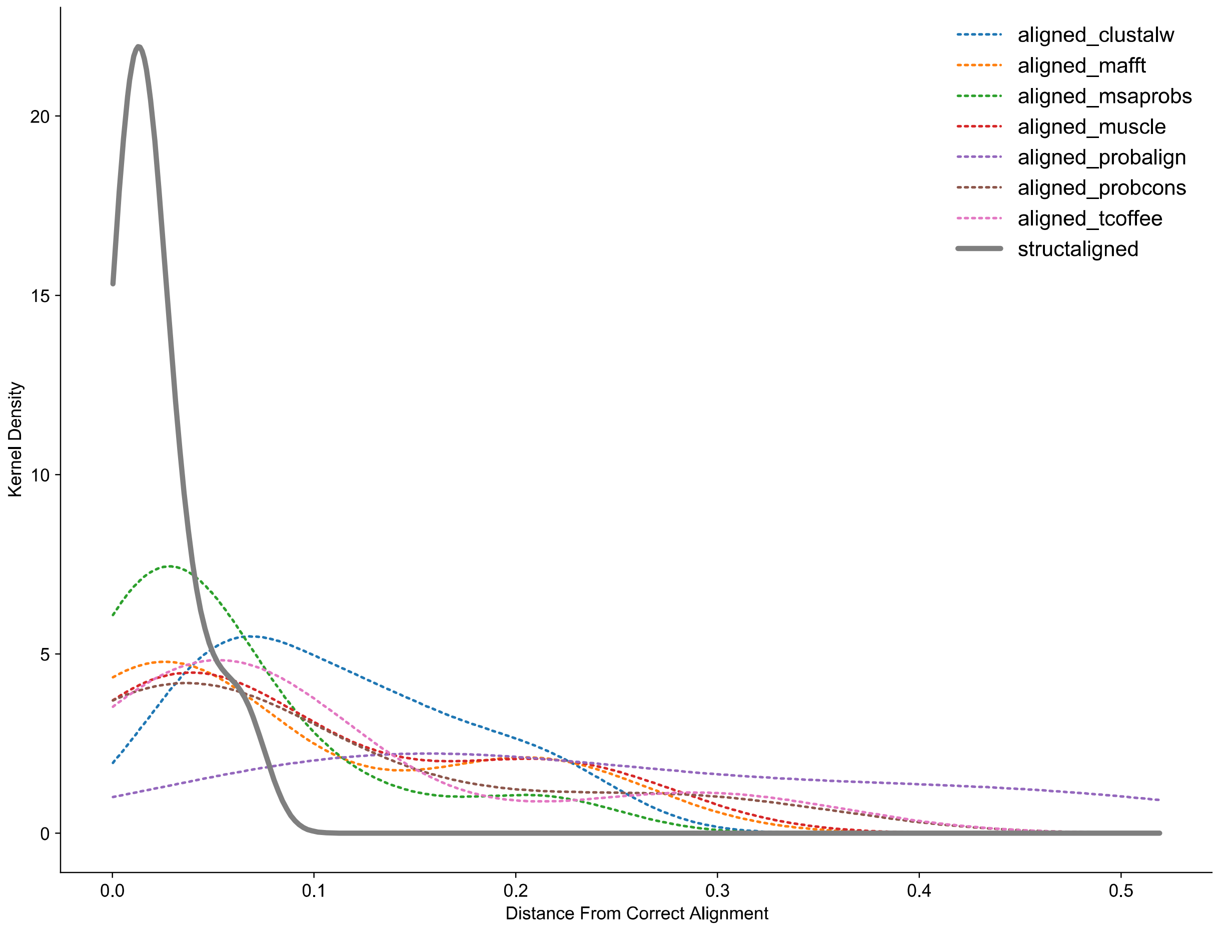


**Figure S3 Part 4, DSRM3 Domain.**

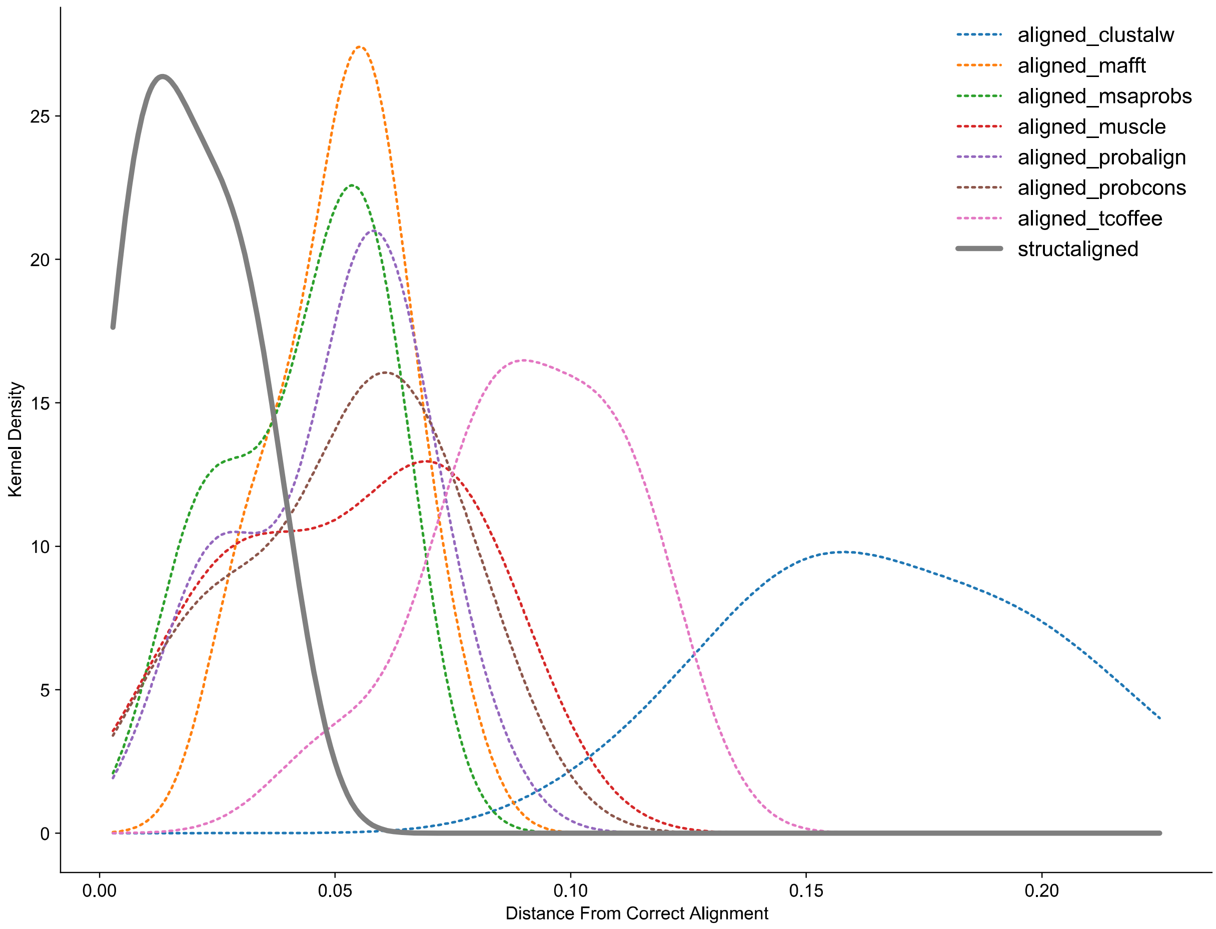


**Figure S3 Part 5, RD Domain.**

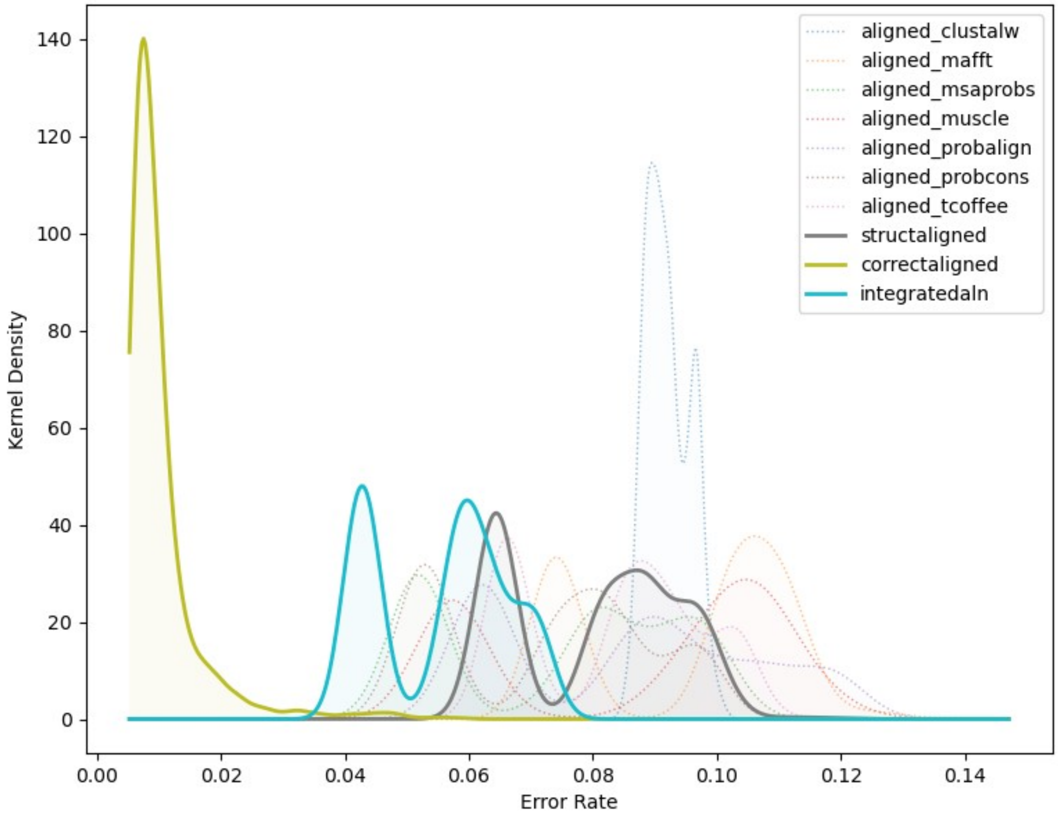


**Figure S4 Part 1, CARD Domain. Total ancestral sequence reconstruction (ASR) error rates are low when the correct alignment is known in advance, higher when the alignment is inferred using sequence-alignment methods, and intermediate when structure-guided alignments or alignment-integrated ASR is used.** We simulated extant and ancestral sequences for 5 protein domain families, using empirically-derived conditions, and aligned the resulting extant sequence data using structure-guided and 7 different sequence-alignment methods (see Methods). We used each alignment to infer the most likely ancestral sequence at each node on the phylogeny. Additionally, alignment-integrated ancestral sequences were generated by combining inferences from the 7 sequence-alignment methods (see Methods). In each case, we compared the inferred ancestral sequences to the correct simulated sequence to estimate ancestral sequence reconstruction (ASR) error rates (see Methods). ASR errors were divided into 4 "errorType" categories: 1) residue errors, in which both correct and inferred ancestral sequences have a residue at a given position in the alignment, but the residues differ; 2) insertion errors, in which the inferred sequence has a residue at a given alignment position, but the correct sequence has a gap state; 3) deletion errors, in which the inferred sequence has a gap, but the correct sequence has a residue, and 4) total errors, which combine residue- insertion- and deletion-errors. Sequence-wide error rates (errors/site) were computed by dividing the number of errors by the length of the pairwise alignment of the inferred and correct ancestral sequences. Error rate distributions were inferred by kernel density estimation. Here we report the total ASR-error distributions. Part 1 shows results for the CARD domain family; results for other domains are shown in parts 2-5 below.

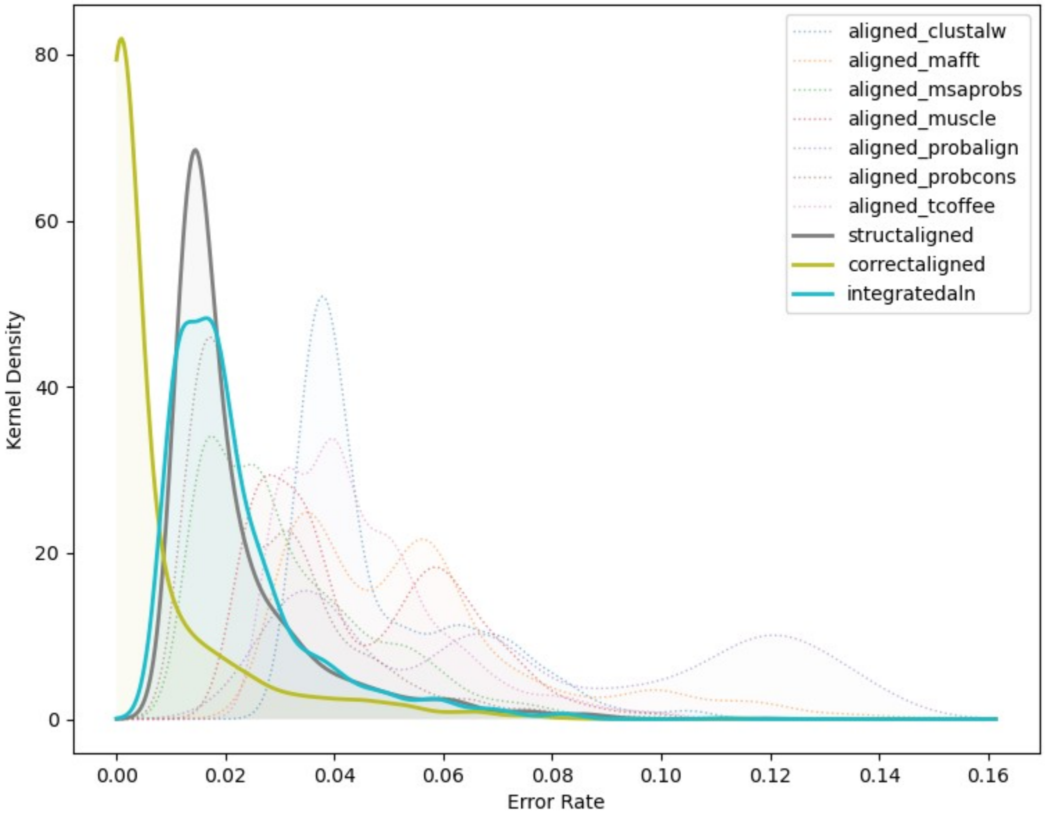


**Figure S4 Part 2, DSRM1 Domain.**

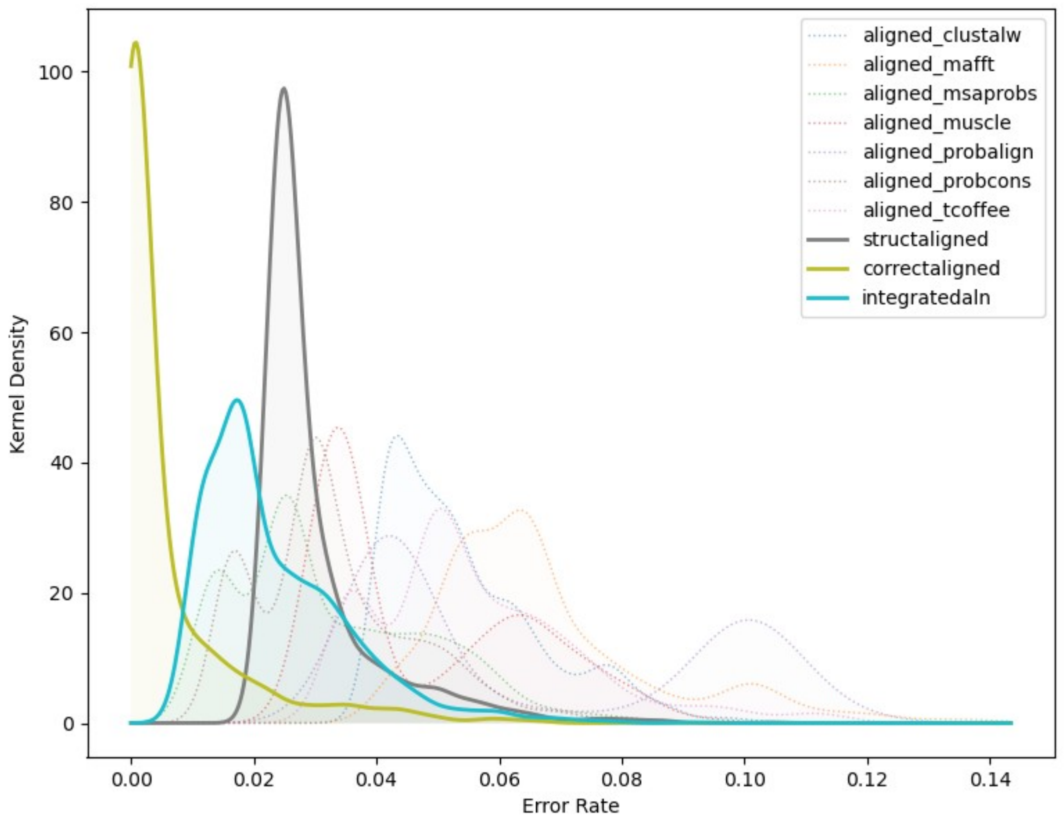


**Figure S4 Part 3, DSRM2 Domain.**

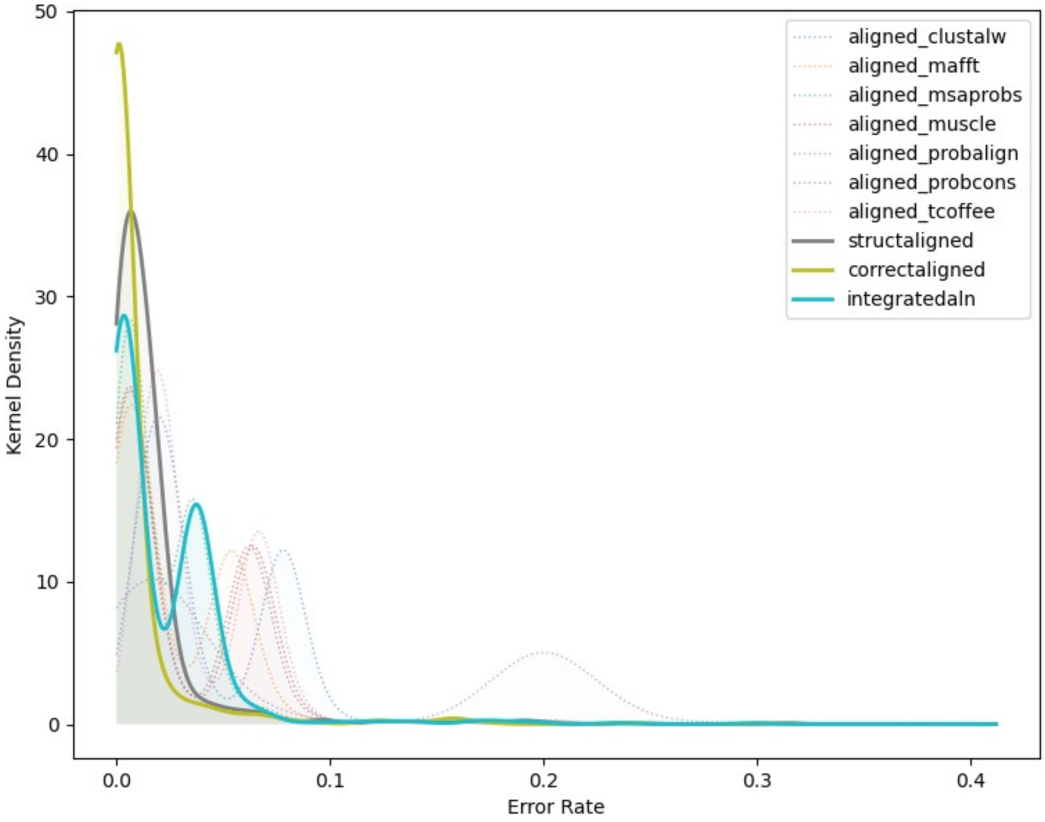


**Figure S4 Part 4, DSRM3 Domain.**

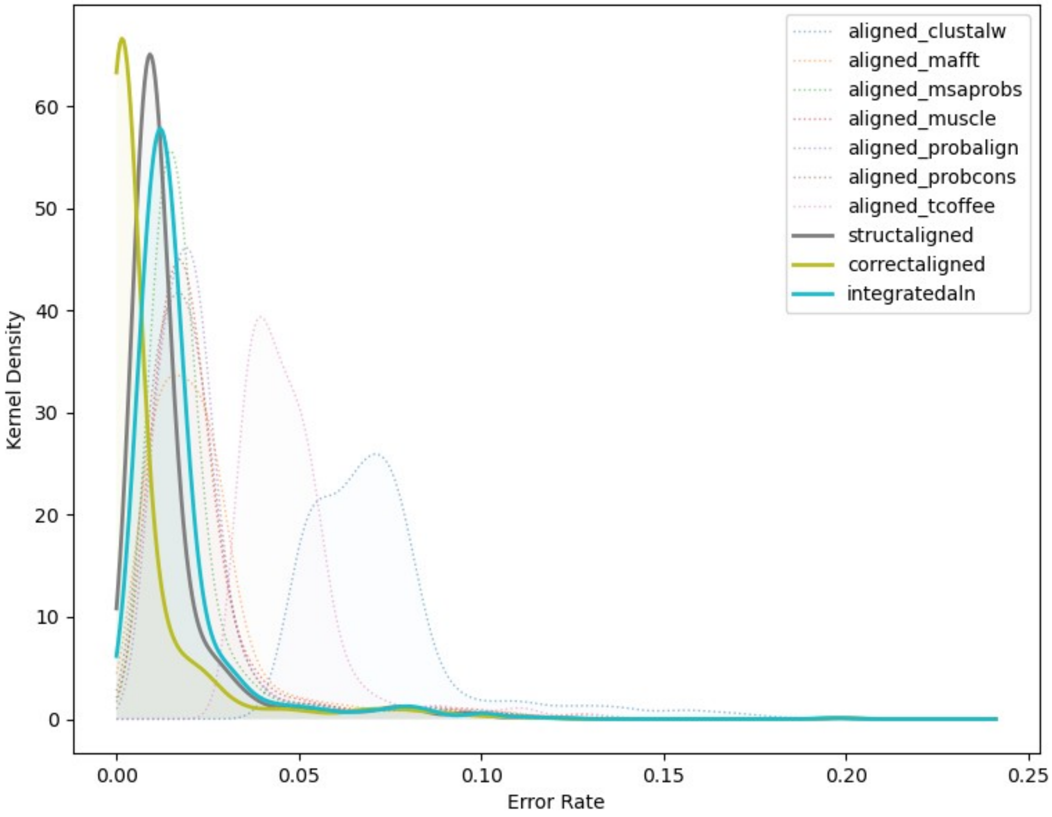


**Figure S4 Part 5, RD Domain.**

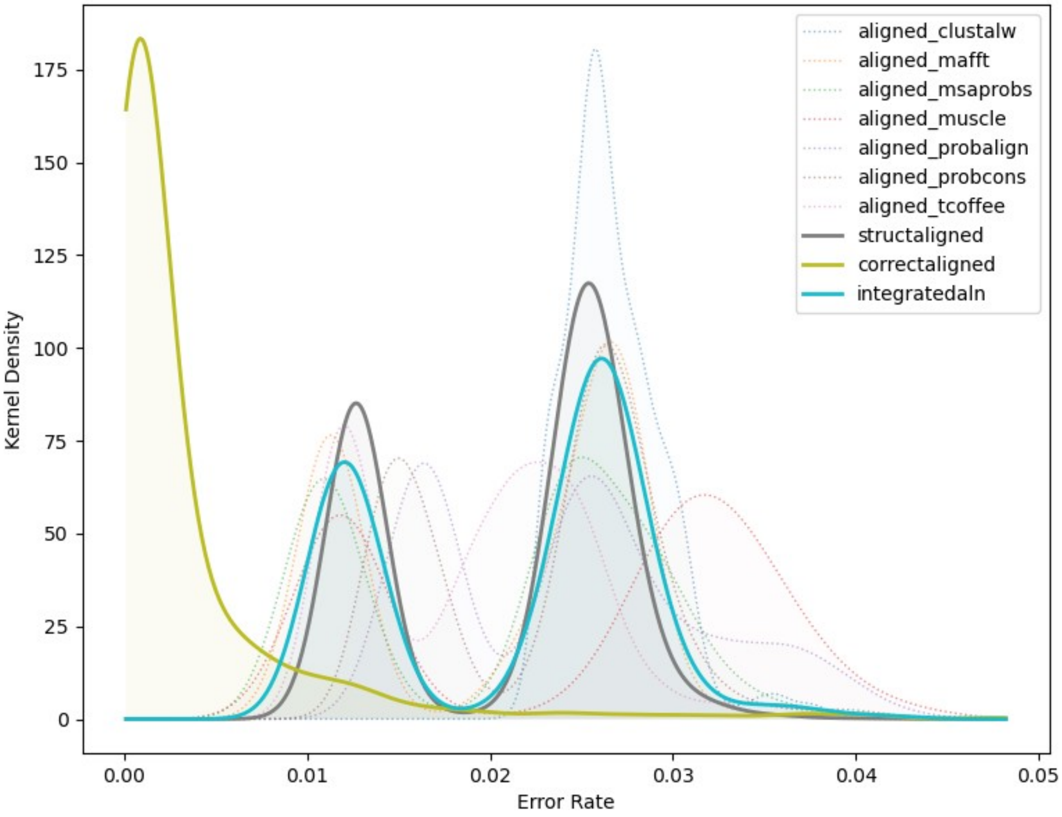


**Figure S5 Part 1, CARD Domain. Ancestral sequence reconstruction (ASR) residue-reconstruction error rates are low when the correct alignment is known in advance, higher when the alignment is inferred using sequence-alignment methods, and intermediate when structure-guided alignments or alignment-integrated ASR is used.** We simulated extant and ancestral sequences for 5 protein domain families, using empirically-derived conditions, and aligned the resulting extant sequence data using structure-guided and 7 different sequence-alignment methods (see Methods). We used each alignment to infer the most likely ancestral sequence at each node on the phylogeny. Additionally, alignment-integrated ancestral sequences were generated by combining inferences from the 7 sequence-alignment methods (see Methods). In each case, we compared the inferred ancestral sequences to the correct simulated sequence to estimate ancestral sequence reconstruction (ASR) error rates (see Methods). ASR errors were divided into 4 “errorType” categories: 1) residue errors, in which both correct and inferred ancestral sequences have a residue at a given position in the alignment, but the residues differ; 2) insertion errors, in which the inferred sequence has a residue at a given alignment position, but the correct sequence has a gap state; 3) deletion errors, in which the inferred sequence has a gap, but the correct sequence has a residue, and 4) total errors, which combine residue- insertion- and deletion-errors. Sequence-wide error rates (errors/site) were computed by dividing the number of errors by the length of the pairwise alignment of the inferred and correct ancestral sequences. Error rate distributions were inferred by kernel density estimation. Here we report the ASR-error distributions for residue errors. Part 1 shows results for the CARD domain family; results for other domains are shown in parts 2-5 below.

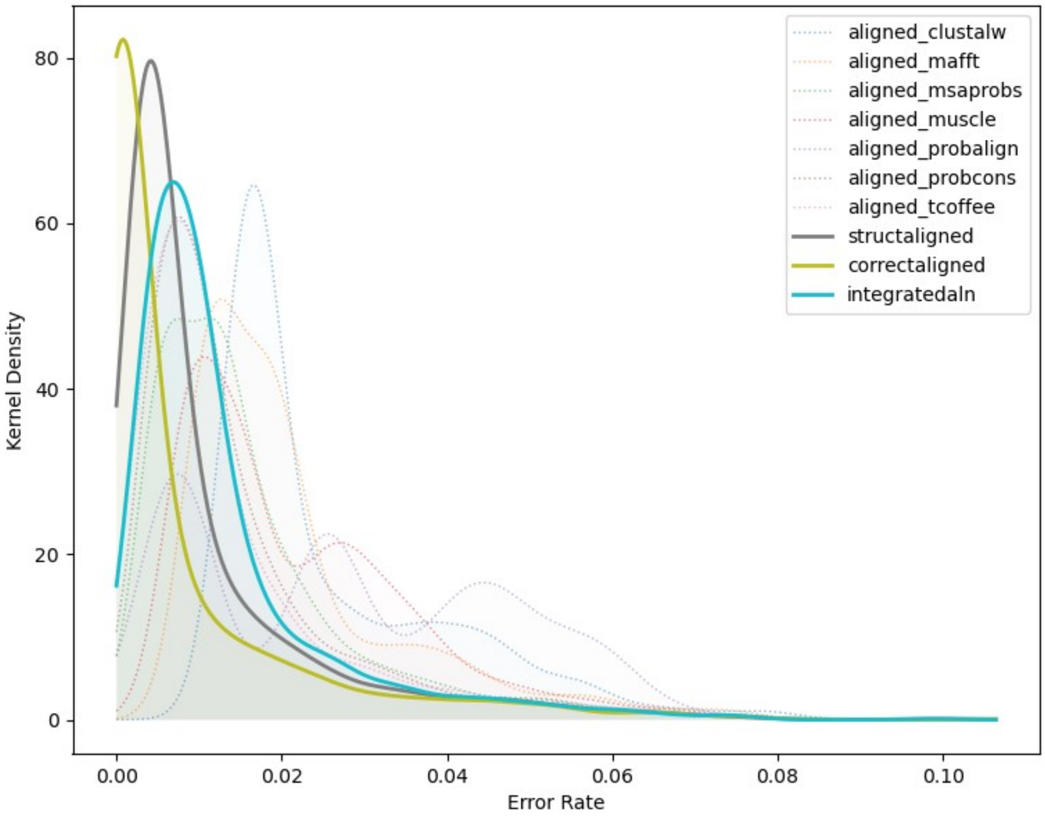


**Figure S5 Part 2, DSRM1 Domain.**

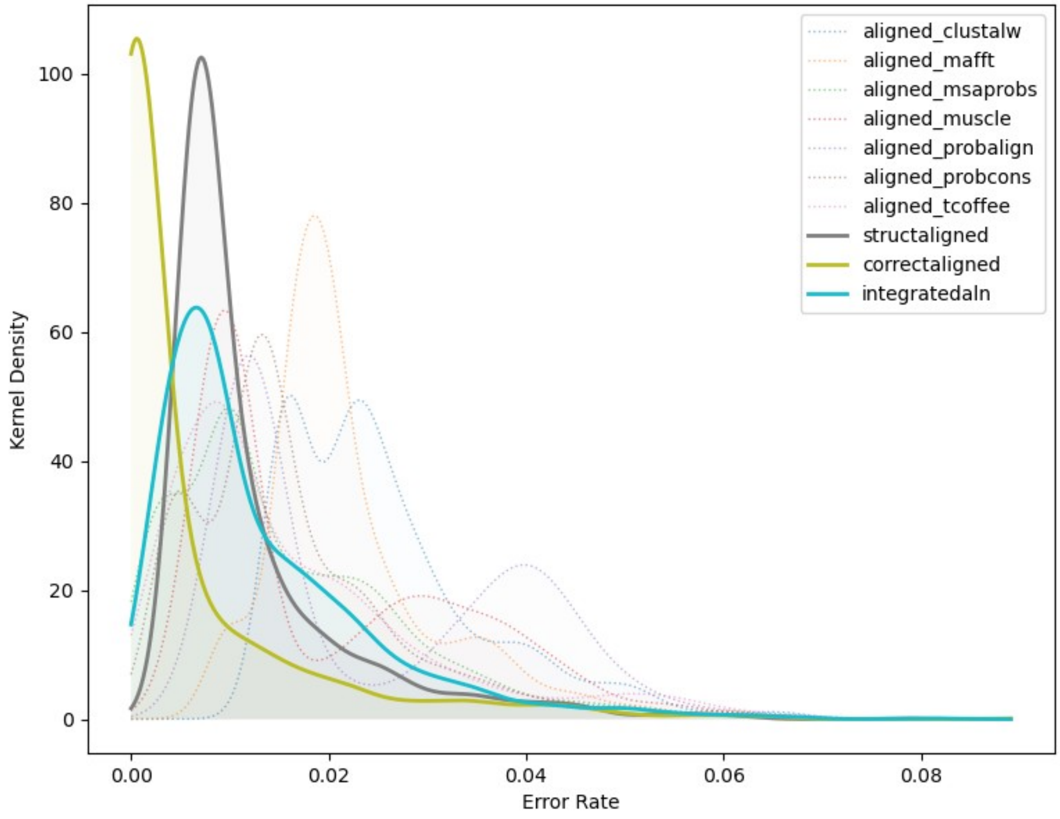


**Figure S5 Part 3, DSRM2 Domain.**

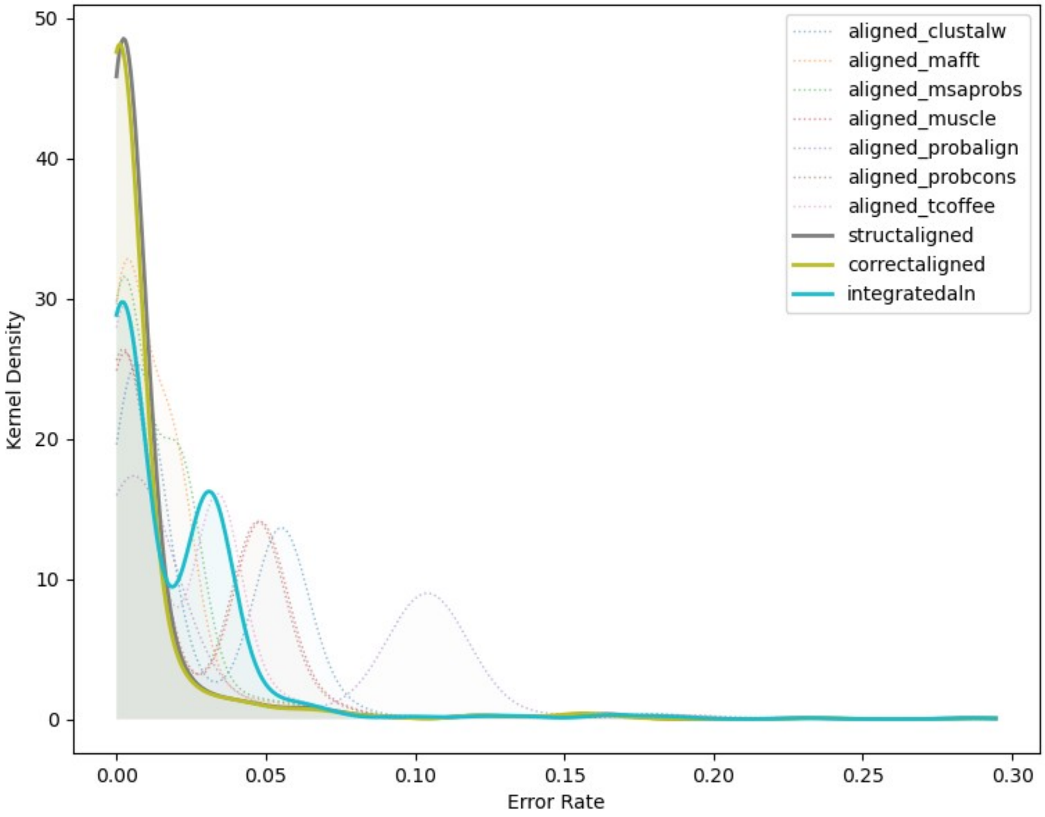


**Figure S5 Part 4, DSRM3 Domain.**

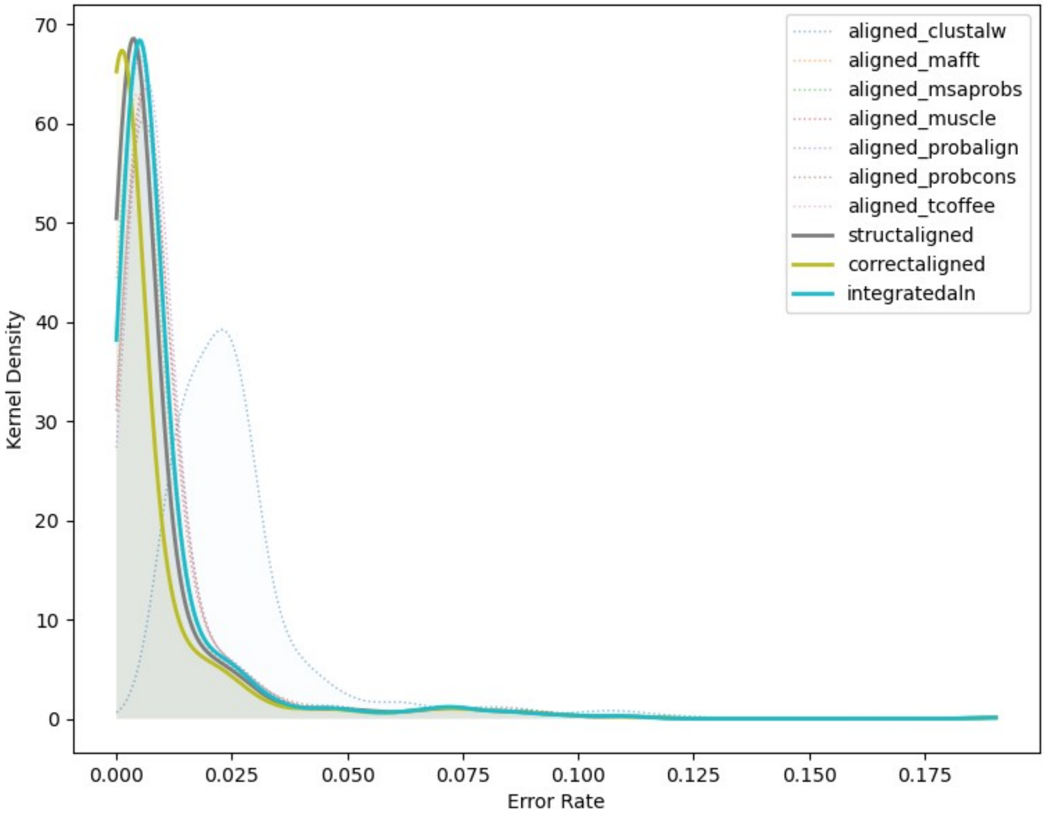


**Figure S5 Part 5, RD Domain.**

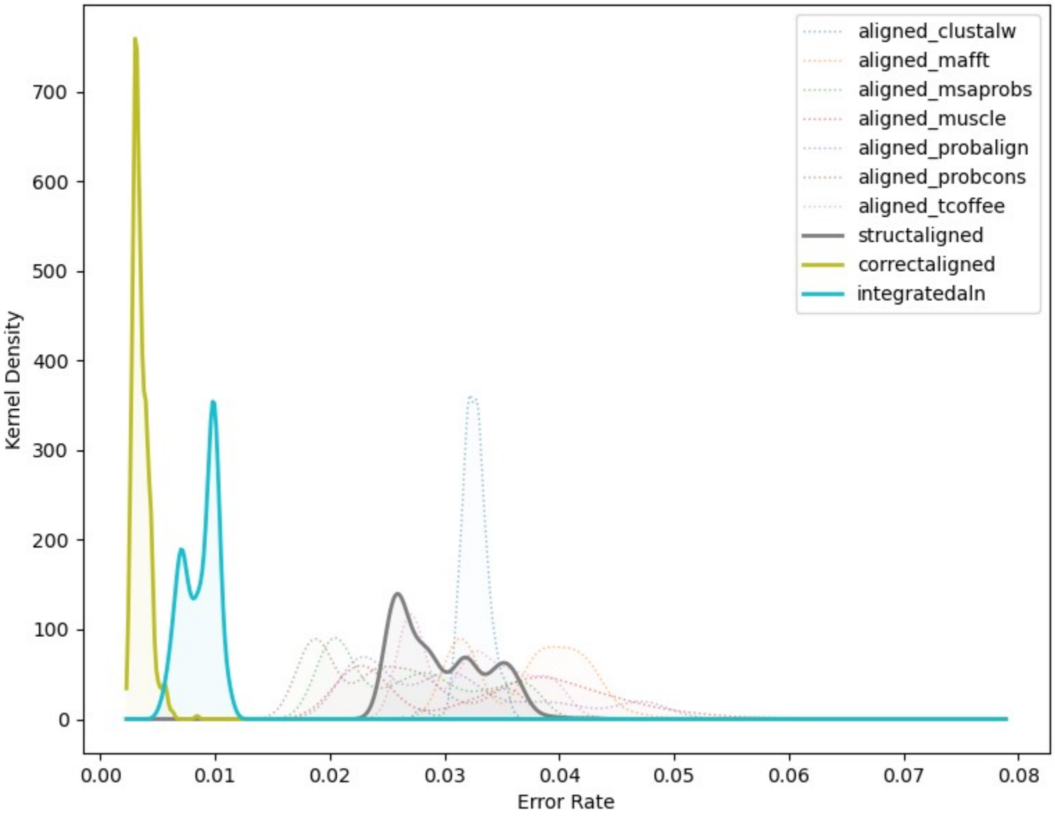


**Figure S6 Part 1, CARD Domain. Ancestral sequence reconstruction (ASR) insertion error rates are low when the correct alignment is known in advance, higher when the alignment is inferred using sequence-alignment methods, and intermediate when structure-guided alignments or alignment-integrated ASR is used.** We simulated extant and ancestral sequences for 5 protein domain families, using empirically-derived conditions, and aligned the resulting extant sequence data using structure-guided and 7 different sequence-alignment methods (see Methods). We used each alignment to infer the most likely ancestral sequence at each node on the phylogeny. Additionally, alignment-integrated ancestral sequences were generated by combining inferences from the 7 sequence-alignment methods (see Methods). In each case, we compared the inferred ancestral sequences to the correct simulated sequence to estimate ancestral sequence reconstruction (ASR) error rates (see Methods). ASR errors were divided into 4 “errorType” categories: 1) residue errors, in which both correct and inferred ancestral sequences have a residue at a given position in the alignment, but the residues differ; 2) insertion errors, in which the inferred sequence has a residue at a given alignment position, but the correct sequence has a gap state; 3) deletion errors, in which the inferred sequence has a gap, but the correct sequence has a residue, and 4) total errors, which combine residue- insertion- and deletion-errors. Sequence-wide error rates (errors/site) were computed by dividing the number of errors by the length of the pairwise alignment of the inferred and correct ancestral sequences. Error rate distributions were inferred by kernel density estimation. Here we report the ASR-error distributions for insertion errors. Part 1 shows results for the CARD domain family; results for other domains are shown in parts 2-5 below.

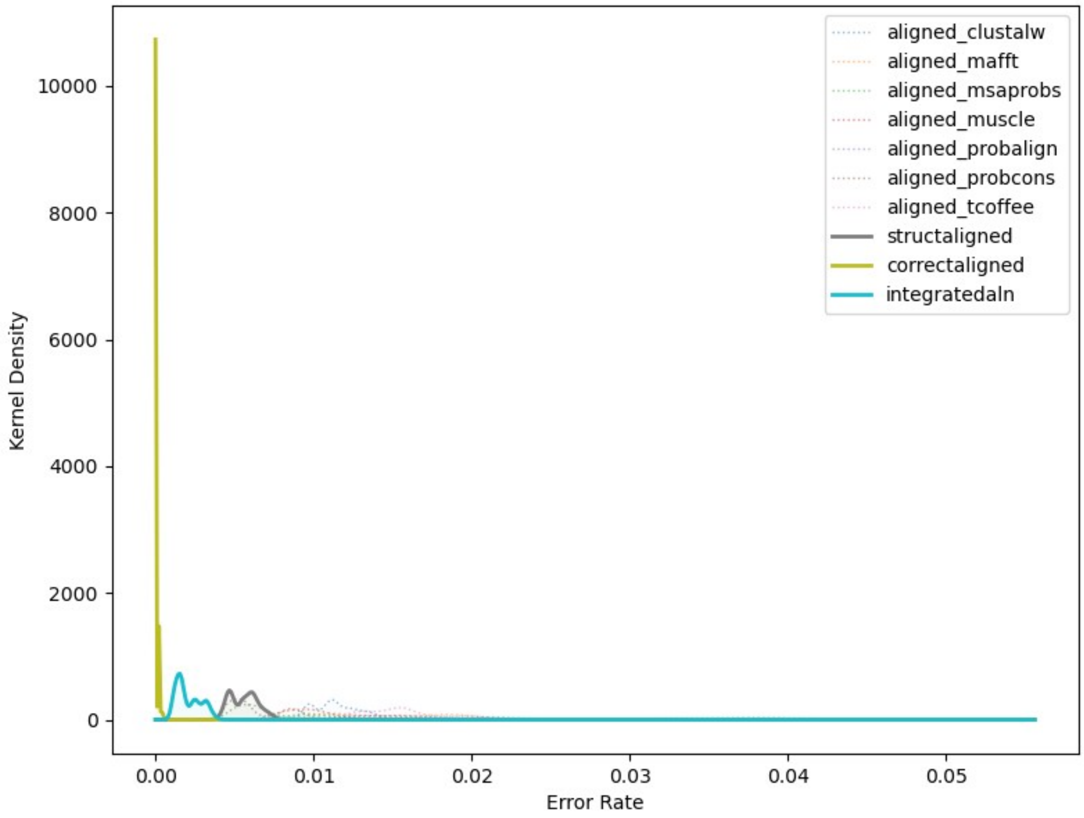


**Figure S6 Part 2, DSRM1 Domain.**

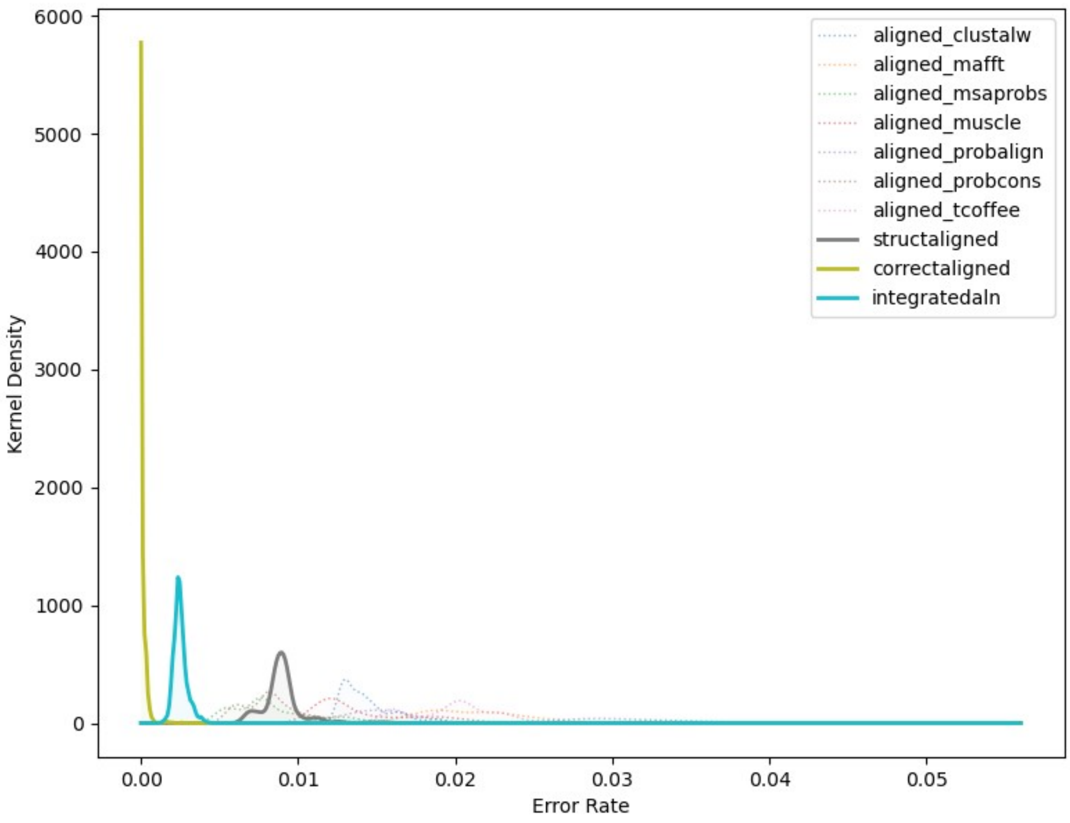


**Figure S6 Part 3, DSRM2 Domain.**

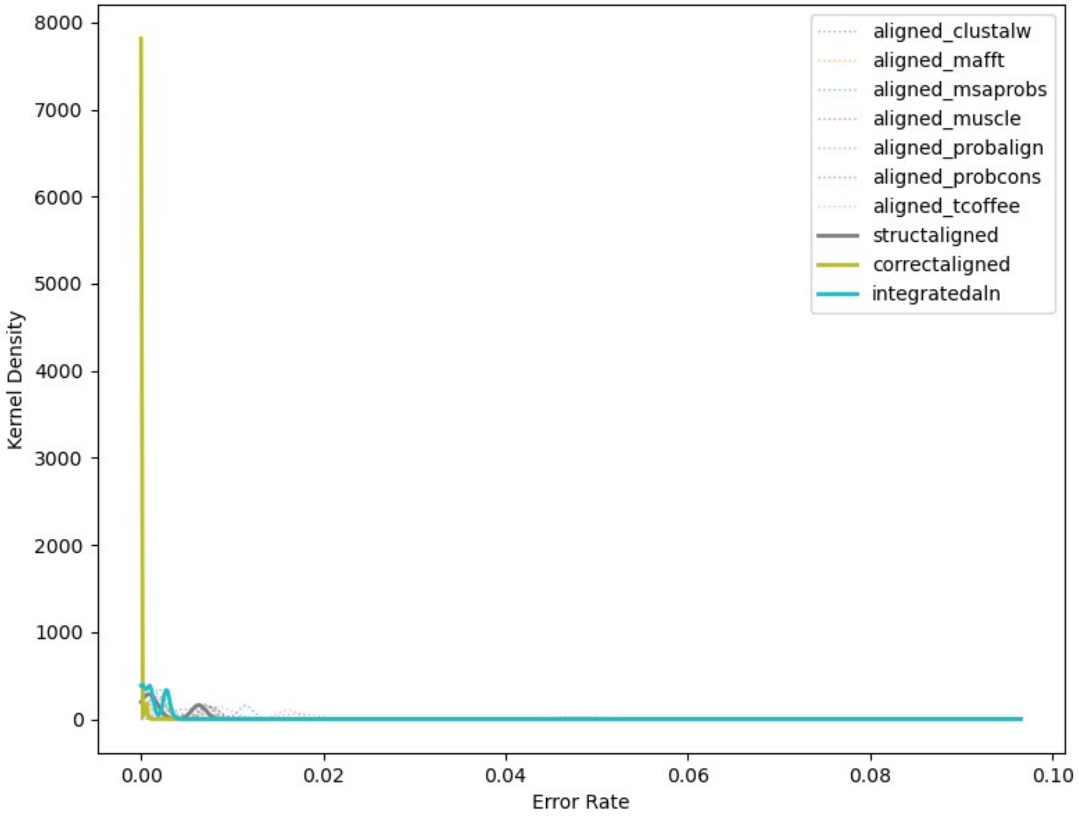


**Figure S6 Part 4, DSRM3 Domain.**

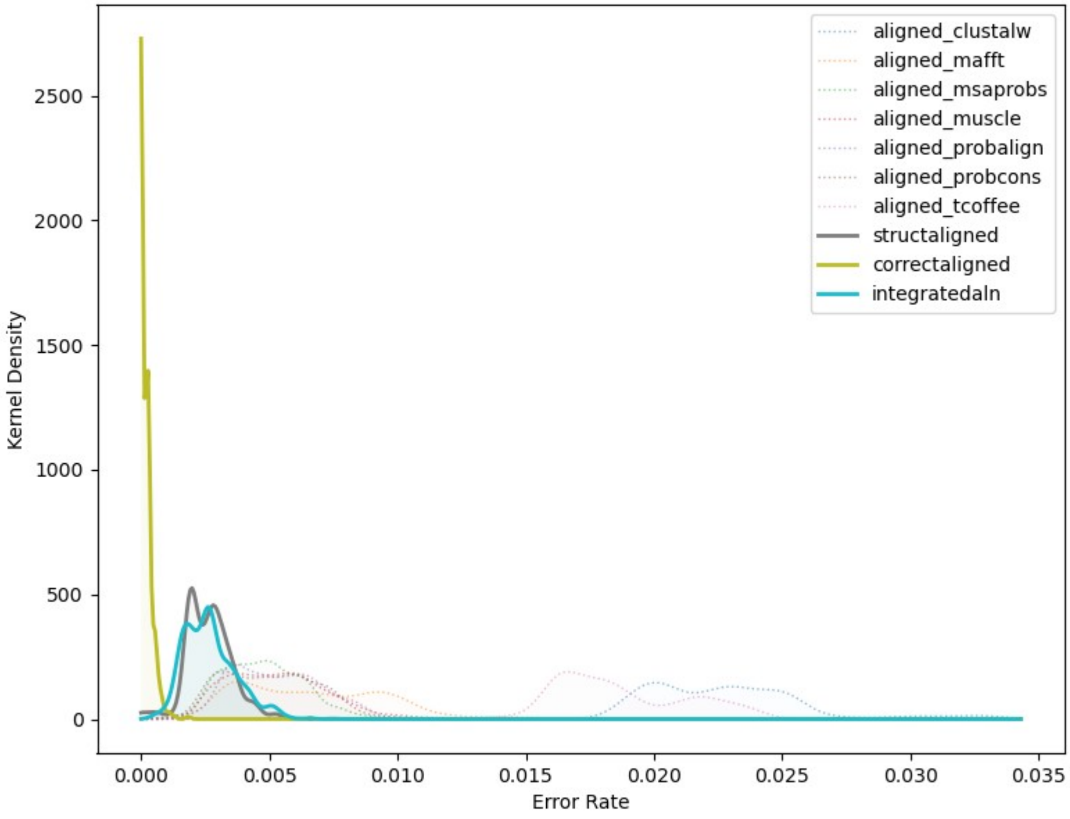


**Figure S6 Part 5, RD Domain.**

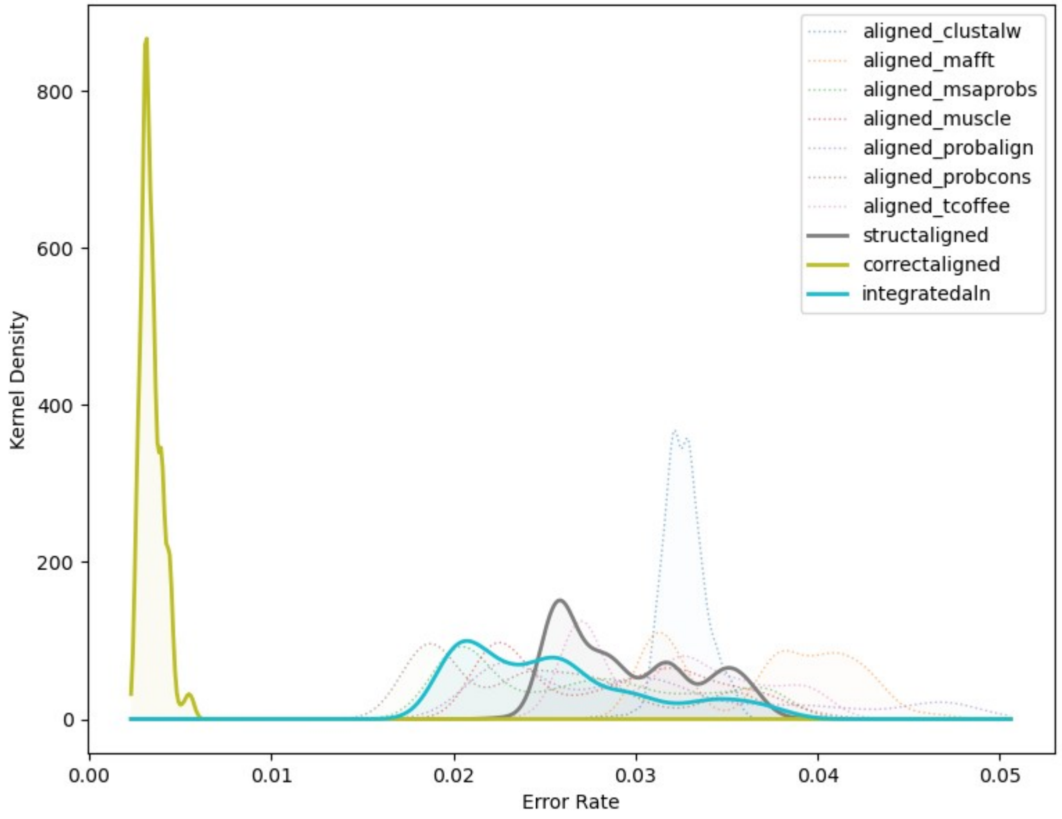


**Figure S7 Part 1, CARD Domain. Ancestral sequence reconstruction (ASR) deletion error rates are low when the correct alignment is known in advance, higher when the alignment is inferred using sequence-alignment methods, and intermediate when structure-guided alignments or alignment-integrated ASR is used.** We simulated extant and ancestral sequences for 5 protein domain families, using empirically-derived conditions, and aligned the resulting extant sequence data using structure-guided and 7 different sequence-alignment methods (see Methods). We used each alignment to infer the most likely ancestral sequence at each node on the phylogeny. Additionally, alignment-integrated ancestral sequences were generated by combining inferences from the 7 sequence-alignment methods (see Methods). In each case, we compared the inferred ancestral sequences to the correct simulated sequence to estimate ancestral sequence reconstruction (ASR) error rates (see Methods). ASR errors were divided into 4 “errorType” categories: 1) residue errors, in which both correct and inferred ancestral sequences have a residue at a given position in the alignment, but the residues differ; 2) insertion errors, in which the inferred sequence has a residue at a given alignment position, but the correct sequence has a gap state; 3) deletion errors, in which the inferred sequence has a gap, but the correct sequence has a residue, and 4) total errors, which combine residue- insertion- and deletion-errors. Sequence-wide error rates (errors/site) were computed by dividing the number of errors by the length of the pairwise alignment of the inferred and correct ancestral sequences. Error rate distributions were inferred by kernel density estimation. Here we report the ASR-error distributions for deletion errors. Part 1 shows results for the CARD domain family; results for other domains are shown in parts 2-5 below.

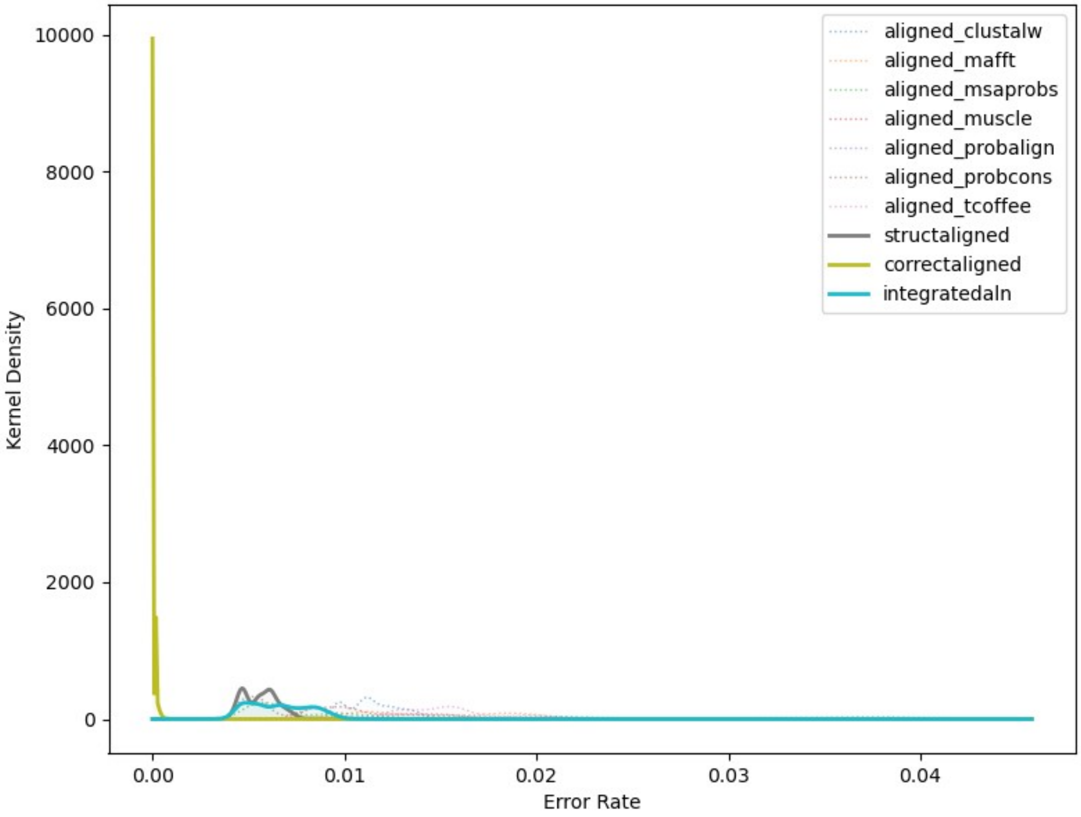


**Figure S7 Part 2, DSRM1 Domain.**

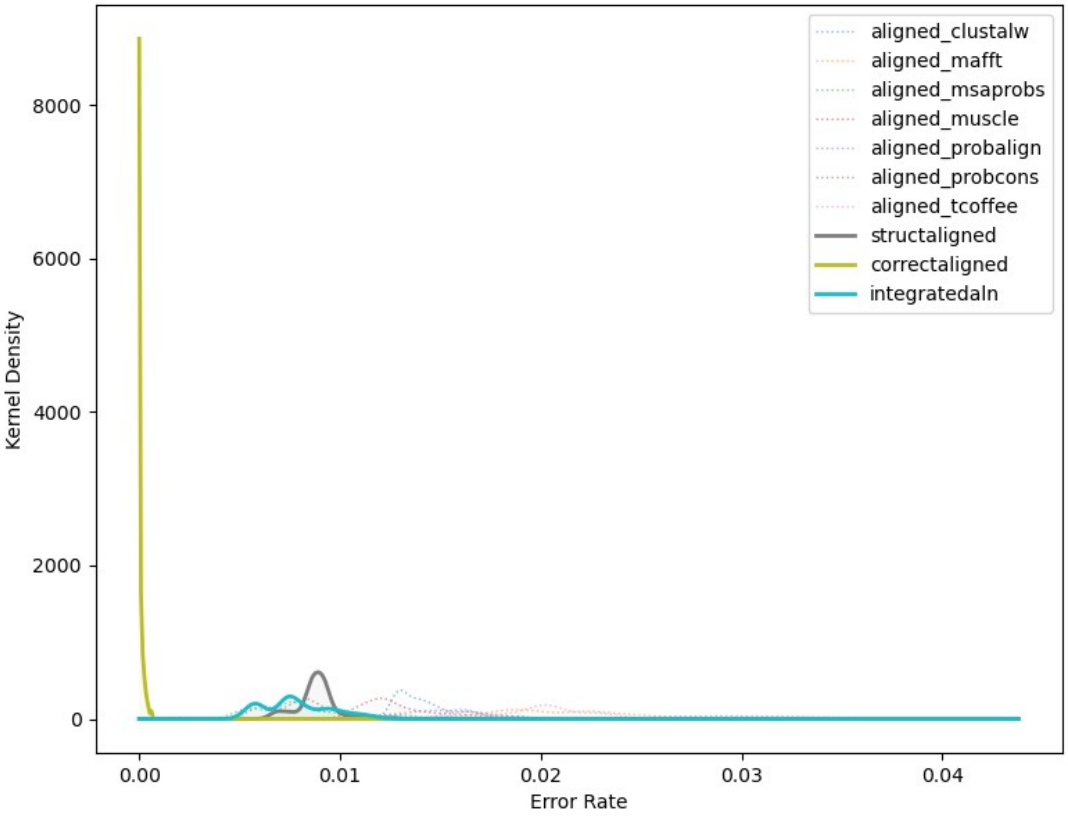


**Figure S7 Part 3, DSRM2 Domain.**

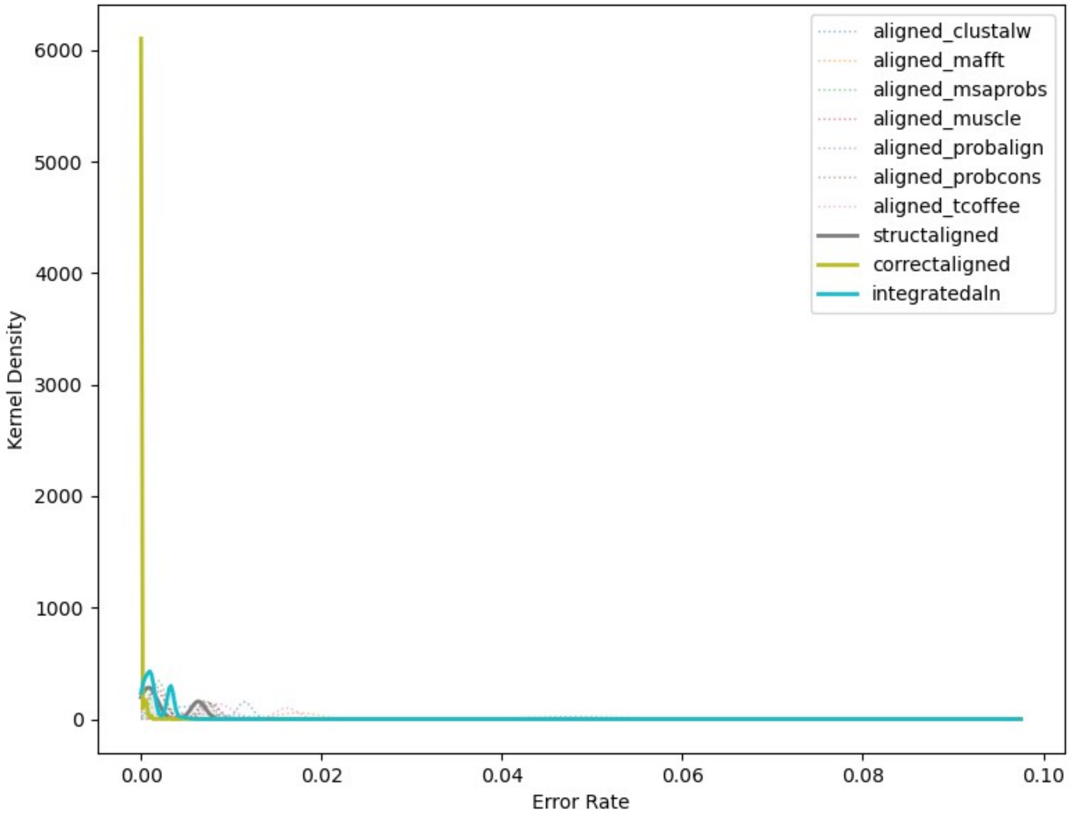


**Figure S7 Part 4, DSRM3 Domain.**

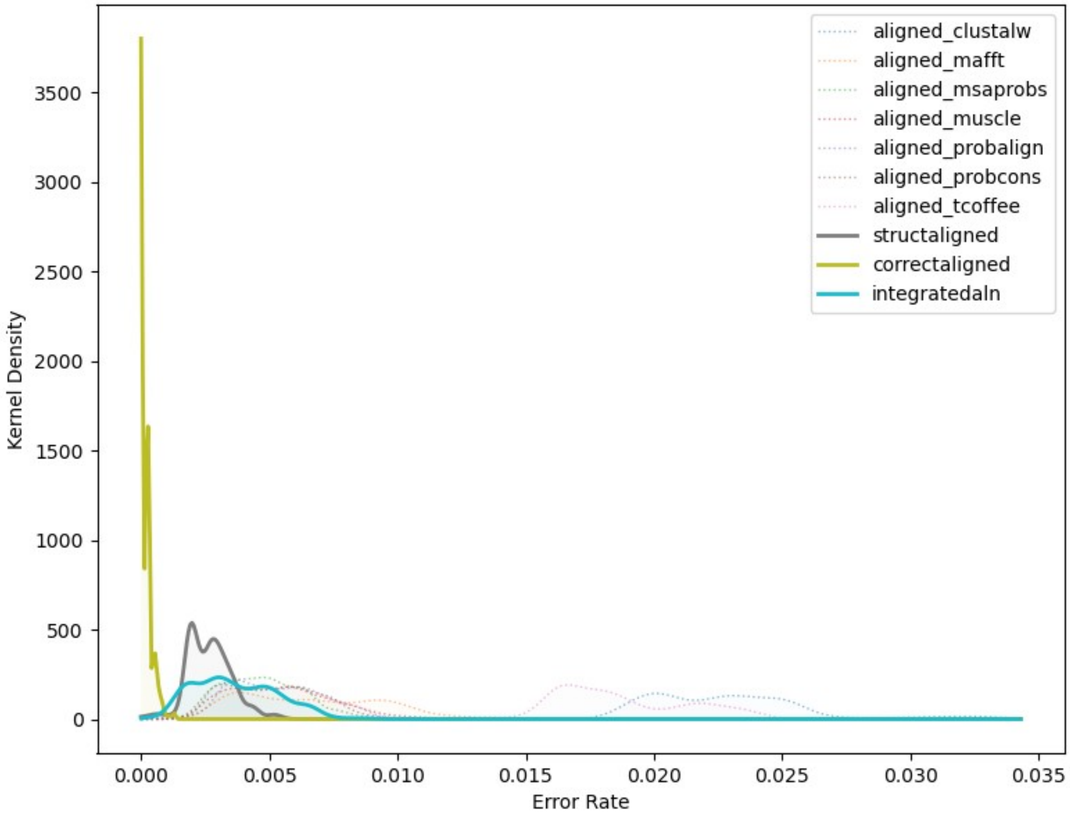


**Figure S7 Part 5, RD Domain.**

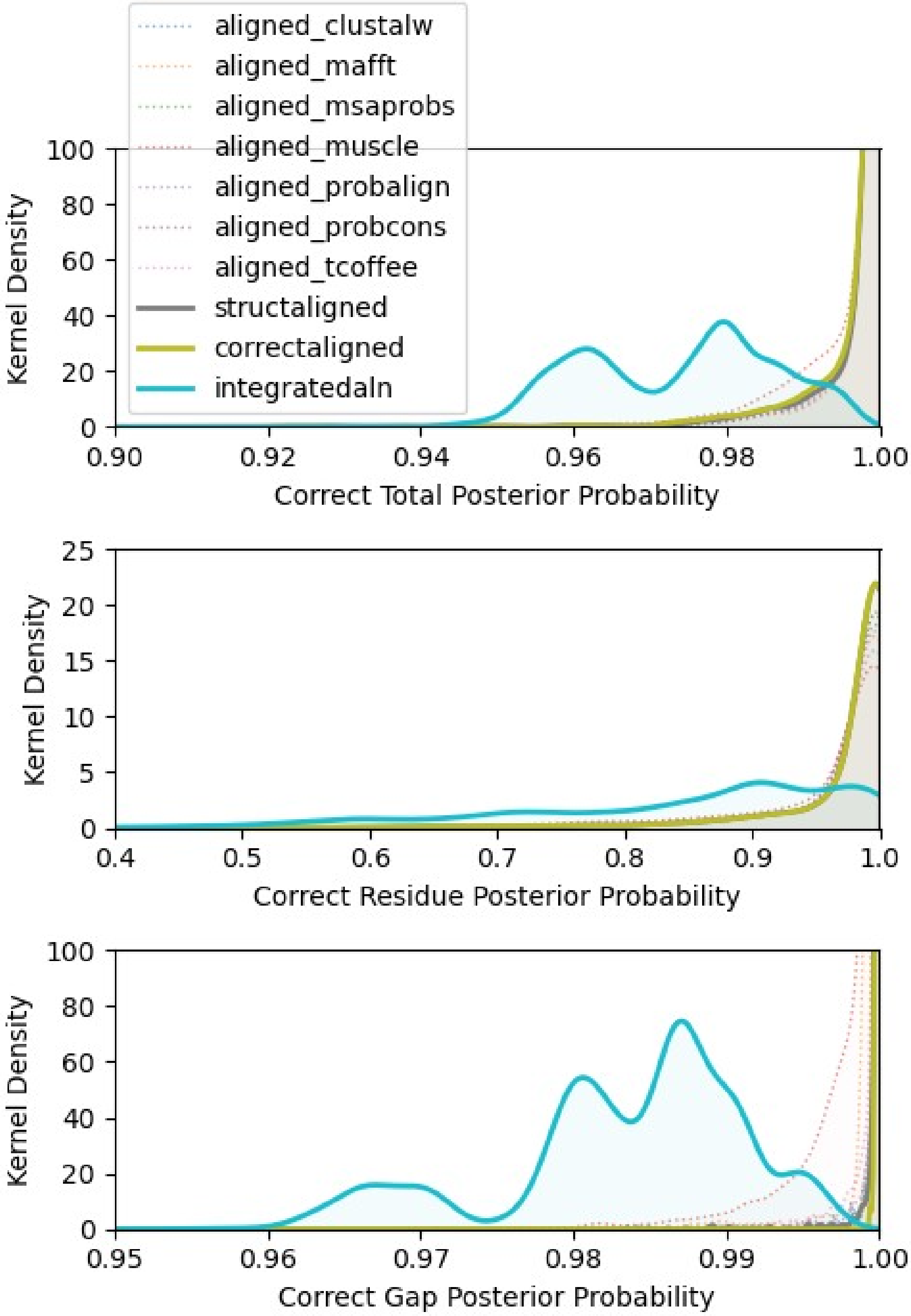


**Figure S8. Support for the correct ancestral state is reduced by alignment-integrated ancestral sequence reconstruction when the correct state is inferred by maximum-likelihood.** We simulated extant and ancestral sequences for 5 protein domain families, using empirically-derived conditions, and aligned the resulting extant sequence data using structure-guided and 7 different sequence-alignment methods (see Methods). We used each alignment to infer the most likely ancestral sequence at each node on the phylogeny. Additionally, alignment-integrated ancestral sequences were generated by combining inferences from the 7 sequence-alignment methods (see Methods). In each case, we compared the inferred ancestral sequences to the correct simulated sequence to identify ancestral sequence reconstruction (ASR) errors (see Methods). When the correct ancestral state was inferred as the maximum-likelihood state, we calculated the posterior probability (PP) of the correct state using an empirical-Bayesian approach (see Methods). The distribution of posterior probabilities for the correct ancestral state, when no ASR error was made, was inferred by kernel density estimation. We report kernel density distributions for correctly-inferred ancestral gap states, residues and total (gaps + residues).

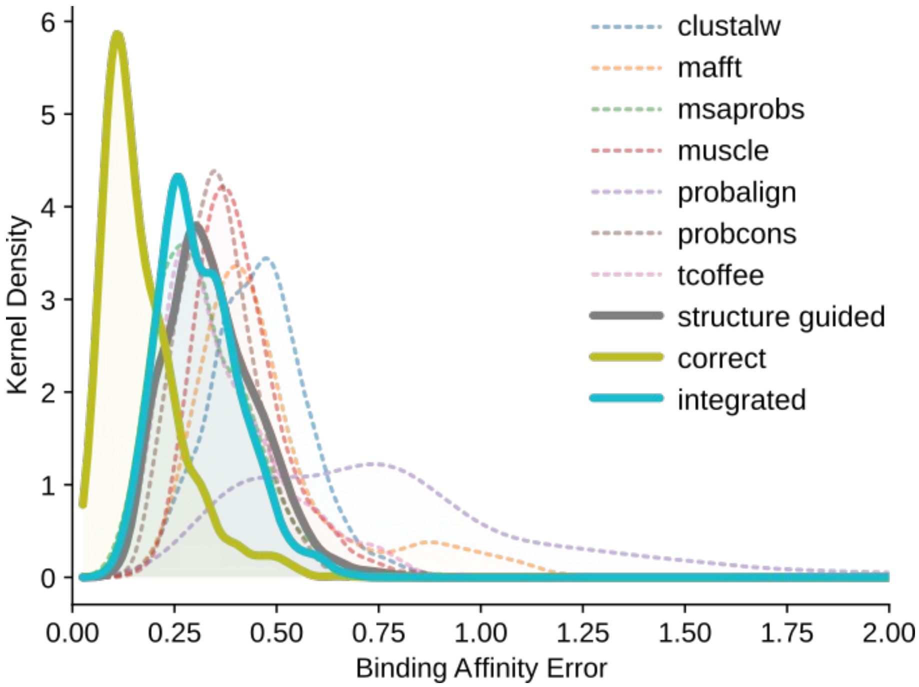


**Figure S9. Alignment-integrated and structure-guided approaches produce less error in inferred structural and functional properties of ancestral proteins than single sequence-alignment methods.** We simulated replicate extant and ancestral sequences by evolving an RNA-binding protein domain along its empirically-determined phylogeny, using a structure-guided alignment to determine the amino-acid composition and pattern of insertions/deletions. Ancestral sequences were inferred using structure-guided alignment, 7 different sequence-alignment methods and alignment-integration. We modeled the structure of each ancestral sequence and estimated its dsRNA binding affinity (pKd) using a computational approach. Errors in binding affinity were calculated by comparing values estimated from the correct ancestral sequences to those estimated using each alignment method. We used kernel density estimation to calculate the frequency distribution of binding-affinity errors.

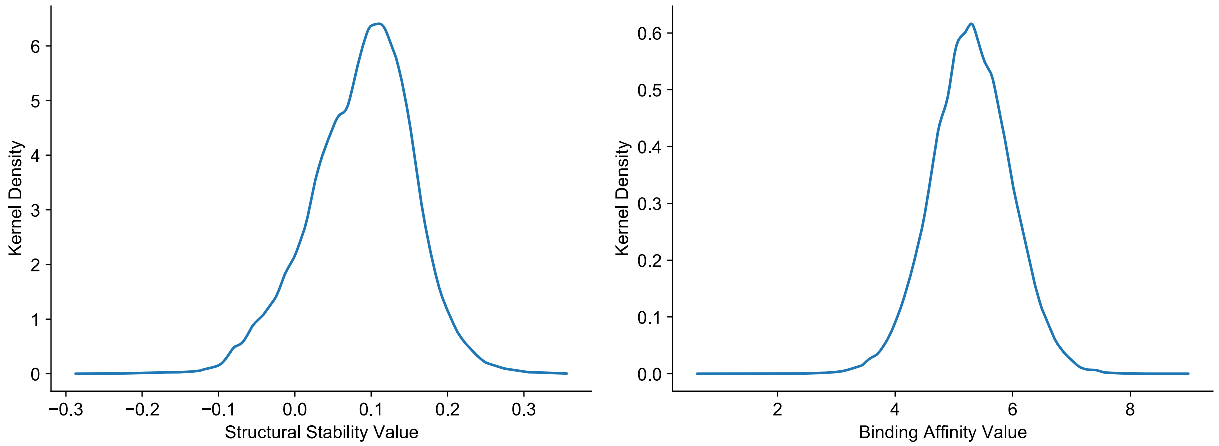


| property | meanValue | medianValue | highestMode | StdDev | SEmean | \| max \| |
|---|---|---|---|---|---|---|
| Structural Stability | 8.74E-02 | 9.37E-02 | 1.10E-01 | 6.74E-02 | 1.82E-04 | 3.56E-01 |
| Binding Affinity | 5.29E+00 | 5.29E+00 | 5.29E+00 | 6.72E-01 | 1.93E-03 | 8.99E+00 |

**Figure S10. Simulated ancestral DSRM1 protein domains generated a range of values for structural stability and dsRNA-binding affinity.** We simulated replicate extant and ancestral sequence data sets by evolving the DSRM1 protein domain family along its empirically-determined phylogeny, using a structure-guided alignment of empirical DSRM1 sequences to determine the amino-acid composition and pattern of insertions/deletions (see Methods). Structural and functional properties were unconstrained during the simulations. We modeled the structure of each simulated ancestral sequence and estimated its structural stability (ΔG) and dsRNA-binding affinity (pKd) using computational approaches (see Methods). The distributions of simulated structural stability (left) and binding affinity (right) values were inferred using kernel density estimation. Table inset reports the mean (and standard error), median, mode, standard deviation and absolute-maximum of each distribution.

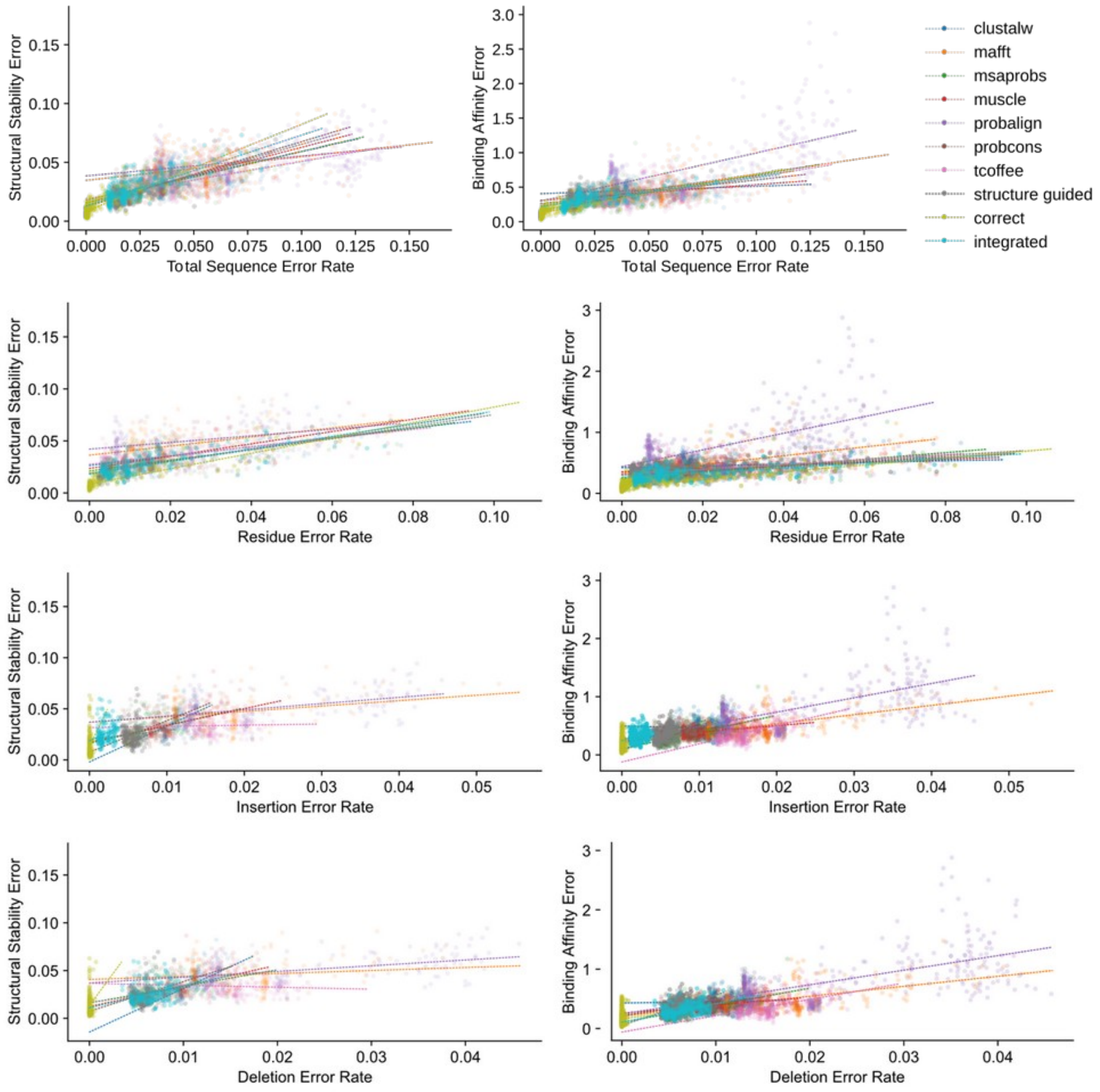


**Figure S11. Errors in estimated structural stability and binding affinity of ancestral sequences are weakly correlated with errors in the inferred ancestral sequences.** We simulated replicate extant and ancestral sequence data sets by evolving the DSRM1 protein domain family along its empirically-determined phylogeny, using a structure-guided alignment of empirical DSRM1 sequences to determine the amino-acid composition and pattern of insertions/ deletions (see Methods). Structural and functional properties were unconstrained during the simulations. We modeled the structure of each simulated ancestral sequence and estimated its structural stability (ΔG) and dsRNA-binding affinity (pKd) using computational approaches (see Methods). For each alignment method, we inferred the maximum-likelihood ancestral sequence at each node on the phylogeny, modeled the structure of the inferred ancestral sequence at each node, and estimated structural stability and binding affinity from the modeled structure. Structural stability and binding affinity errors were calculated as the absolute value of the difference between the value calculated from the correct simulated ancestral sequence and that calculated from the inferred ancestral sequence. ASR errors were divided into 4 “errorType” categories: 1) residue errors, in which both correct and inferred ancestral sequences have a residue at a given position in the alignment, but the residues differ; 2) insertion errors, in which the inferred sequence has a residue at a given alignment position, but the correct sequence has a gap state; 3) deletion errors, in which the inferred sequence has a gap, but the correct sequence has a residue, and 4) total errors, which combine residue- insertion- and deletion-errors. Sequence-wide error rates (errors/site) were computed by dividing the number of errors by the length of the pairwise alignment of the inferred and correct ancestral sequences. For each type of ASR error and alignment method, we determined the best-fit linear relationship between ASR error rate and structural stability (left column) or binding affinity (right column) using ordinary least squares linear regression (dotted lines). Results for total ASR error rates are reported in the main text.

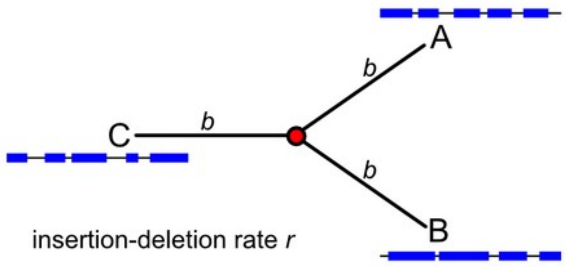


**Figure S12. A minimal 3-taxon simulation provides a model system for investigating ancestral sequence reconstruction errors.** We simulated protein sequences along a 3-taxon phylogeny with equal branch lengths (*b* substitutions/site) and the same insertion-deletion (indel) rate across the phylogeny (*r* indels/substitution/site). Red dot indicates the single ancestral node, and blue bars indicate possible aligned extant sequences generated by the simulation process.

While large-scale simulations are useful for evaluating the potential impact of alignment errors on ancestral sequence reconstruction studies in practice, it is difficult to systematically isolate the relevant variables necessary to determine why alignment errors generate ASR errors from such large-scale systems. Small model problems are needed to reliably infer causative mechanisms. In phylogenetic inference, the ‘4-taxon problem’ represents the smallest possible tree with multiple unrooted topologies, and studies of this simple ‘toy’ problem have led to major advancements in our understanding of the causes of phylogenetic inference errors and their potential solutions.

Here we propose a ‘3-taxon problem’ as the equivalent minimalist system for examining ancestral sequence reconstruction accuracy. For each node on the phylogeny, the most commonly-used marginal ASR algorithms use information from the 3 directly-connected nodes (typically 2 descendent nodes (eg, A,B) and one node representing the local ‘root’ (eg, C)) to infer the ancestral state probability distribution. The simple 3-taxon problem therefore allows us to learn potentially generalizable information about the factors impacting ASR accuracy while limiting the number of simulation variables to a small number.

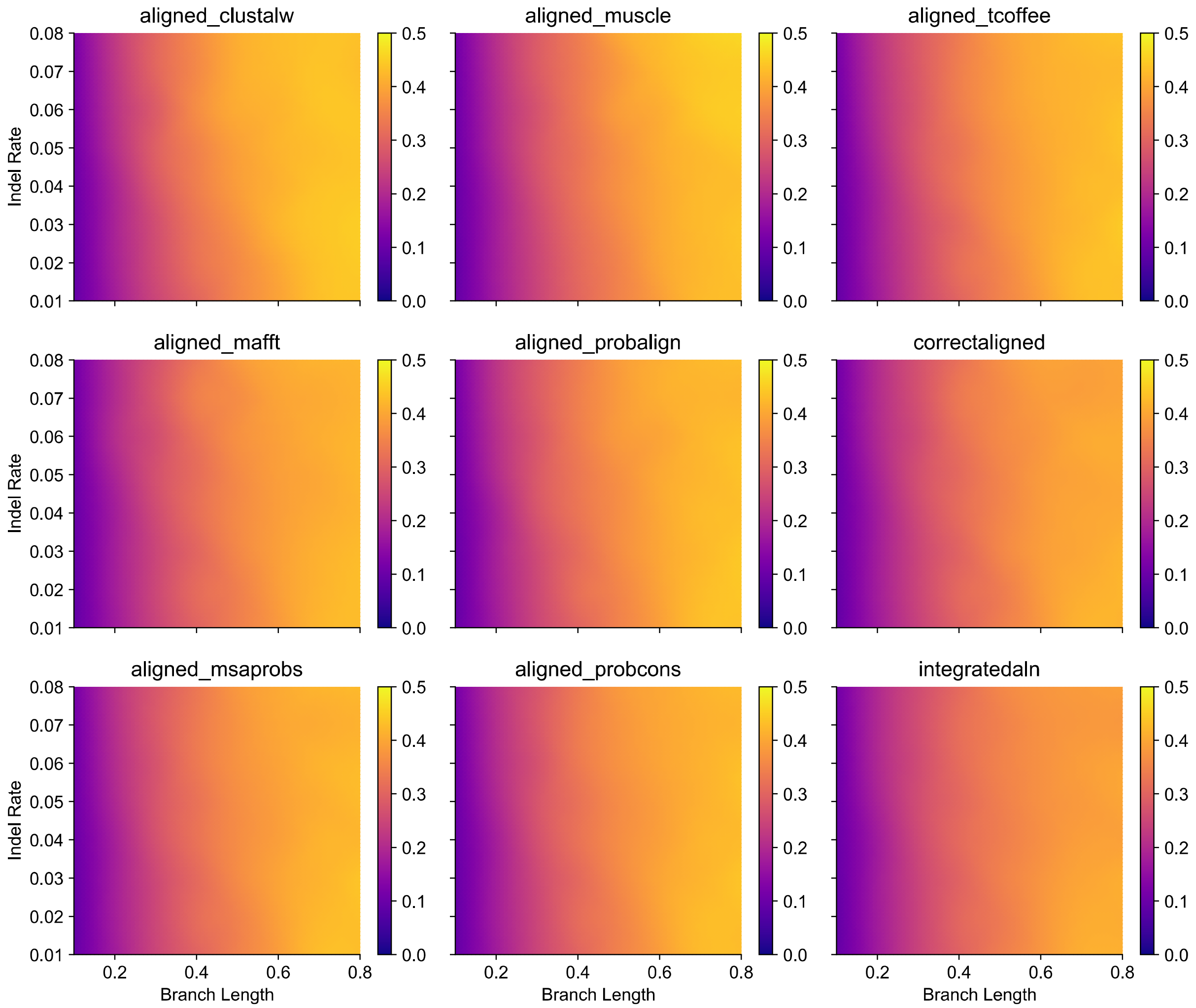


**Figure S13. Alignment-integrated ancestral sequence reconstruction reduces total ASR error rates in 3-taxon simulations across a variety of conditions.** We simulated protein sequences along a 3-taxon phylogeny with equal branch lengths and the same insertion-deletion (indel) rate across the phylogeny (see Methods, Supplementary Fig. S12). The resulting extant sequence data was aligned using 7 different sequence-alignment methods (see Methods). We used each alignment to infer the most likely ancestral sequence at the single node on the 3-taxon phylogeny. Additionally, alignment-integrated ancestral sequences were generated by combining inferences from the 7 sequence-alignment methods (see Methods). In each case, we compared the inferred ancestral sequence to the correct simulated sequence to estimate ancestral sequence reconstruction (ASR) error rates (see Methods). ASR errors were divided into 4 categories: 1) residue errors, in which both correct and inferred ancestral sequences have a residue at a given position in the alignment, but the residues differ; 2) insertion errors, in which the inferred sequence has a residue at a given alignment position, but the correct sequence has a gap state; 3) deletion errors, in which the inferred sequence has a gap, but the correct sequence has a residue, and 4) total errors, which combine residue-insertion- and deletion-errors. Sequence-wide error rates (errors/site) were computed by dividing the number of errors by the length of the pairwise alignment of the inferred and correct ancestral sequences. For each combination of branch-length and indel-rate, we plot the average rate of total ASR errors over 100 replicate data sets for each alignment method, with purple indicating low ASR error rates, and yellow indicating high rates of ASR errors.

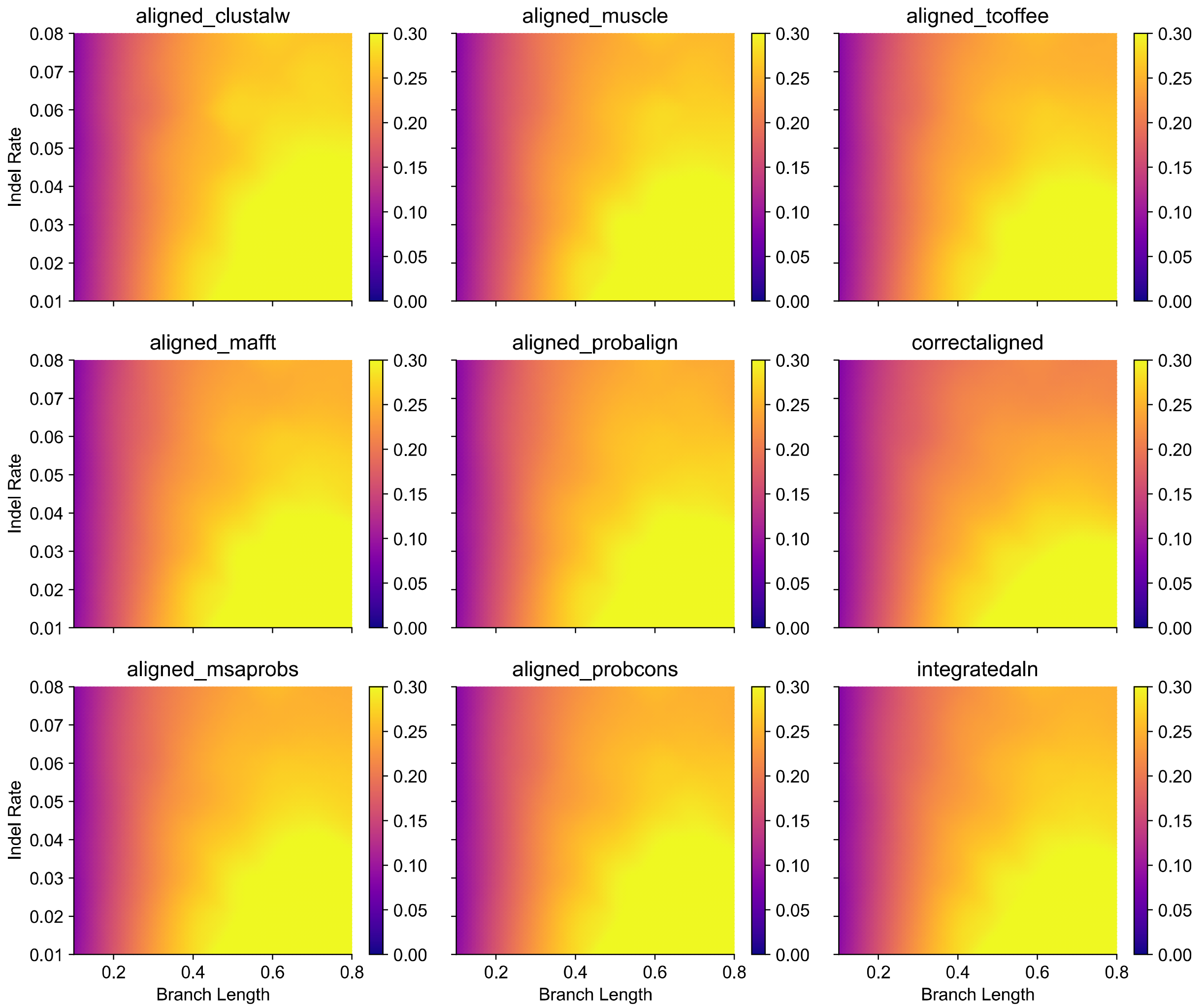


**Figure S14. Alignment-integrated ancestral sequence reconstruction reduces residue ASR error rates in 3-taxon simulations across a variety of conditions.** We simulated protein sequences along a 3-taxon phylogeny with equal branch lengths and the same insertion-deletion (indel) rate across the phylogeny (see Methods, Supplementary Fig. S12). The resulting extant sequence data was aligned using 7 different sequence-alignment methods (see Methods). We used each alignment to infer the most likely ancestral sequence at the single node on the 3-taxon phylogeny. Additionally, alignment-integrated ancestral sequences were generated by combining inferences from the 7 sequence-alignment methods (see Methods). In each case, we compared the inferred ancestral sequence to the correct simulated sequence to estimate ancestral sequence reconstruction (ASR) error rates (see Methods). ASR errors were divided into 4 categories: 1) residue errors, in which both correct and inferred ancestral sequences have a residue at a given position in the alignment, but the residues differ; 2) insertion errors, in which the inferred sequence has a residue at a given alignment position, but the correct sequence has a gap state; 3) deletion errors, in which the inferred sequence has a gap, but the correct sequence has a residue, and 4) total errors, which combine residue-insertion- and deletion-errors. Sequence-wide error rates (errors/site) were computed by dividing the number of errors by the length of the pairwise alignment of the inferred and correct ancestral sequences. For each combination of branch-length and indel-rate, we plot the average rate of residue ASR errors over 100 replicate data sets for each alignment method, with purple indicating low ASR error rates, and yellow indicating high rates of ASR errors.

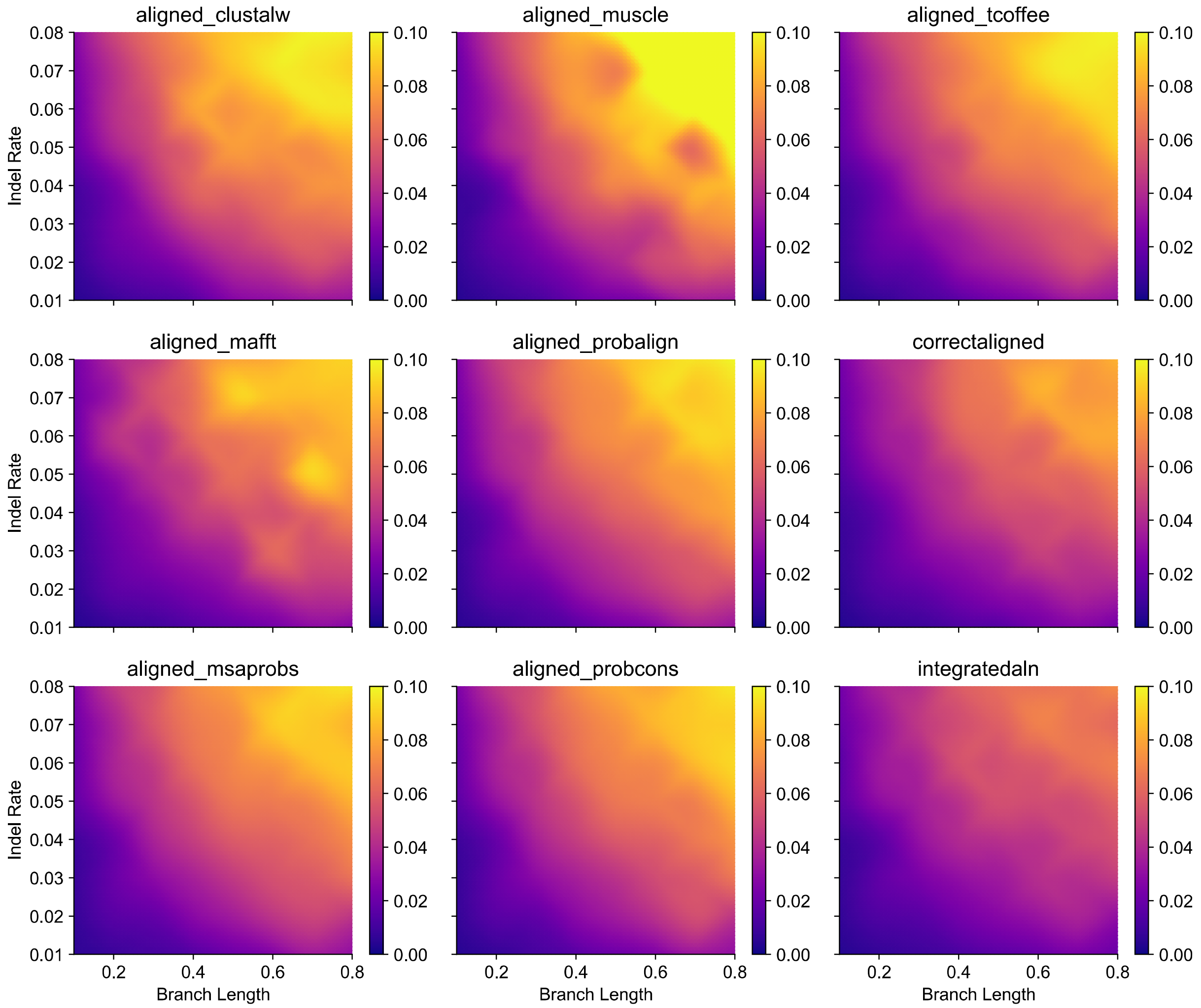


**Figure S15. Alignment-integrated ancestral sequence reconstruction reduces insertion ASR error rates in 3-taxon simulations across a variety of conditions.** We simulated protein sequences along a 3-taxon phylogeny with equal branch lengths and the same insertion-deletion (indel) rate across the phylogeny (see Methods, Supplementary Fig. S12). The resulting extant sequence data was aligned using 7 different sequence-alignment methods (see Methods). We used each alignment to infer the most likely ancestral sequence at the single node on the 3-taxon phylogeny. Additionally, alignment-integrated ancestral sequences were generated by combining inferences from the 7 sequence-alignment methods (see Methods). In each case, we compared the inferred ancestral sequence to the correct simulated sequence to estimate ancestral sequence reconstruction (ASR) error rates (see Methods). ASR errors were divided into 4 categories: 1) residue errors, in which both correct and inferred ancestral sequences have a residue at a given position in the alignment, but the residues differ; 2) insertion errors, in which the inferred sequence has a residue at a given alignment position, but the correct sequence has a gap state; 3) deletion errors, in which the inferred sequence has a gap, but the correct sequence has a residue, and 4) total errors, which combine residue- insertion- and deletion-errors. Sequence-wide error rates (errors/site) were computed by dividing the number of errors by the length of the pairwise alignment of the inferred and correct ancestral sequences. For each combination of branch-length and indel-rate, we plot the average rate of insertion ASR errors over 100 replicate data sets for each alignment method, with purple indicating low ASR error rates, and yellow indicating high rates of ASR errors.

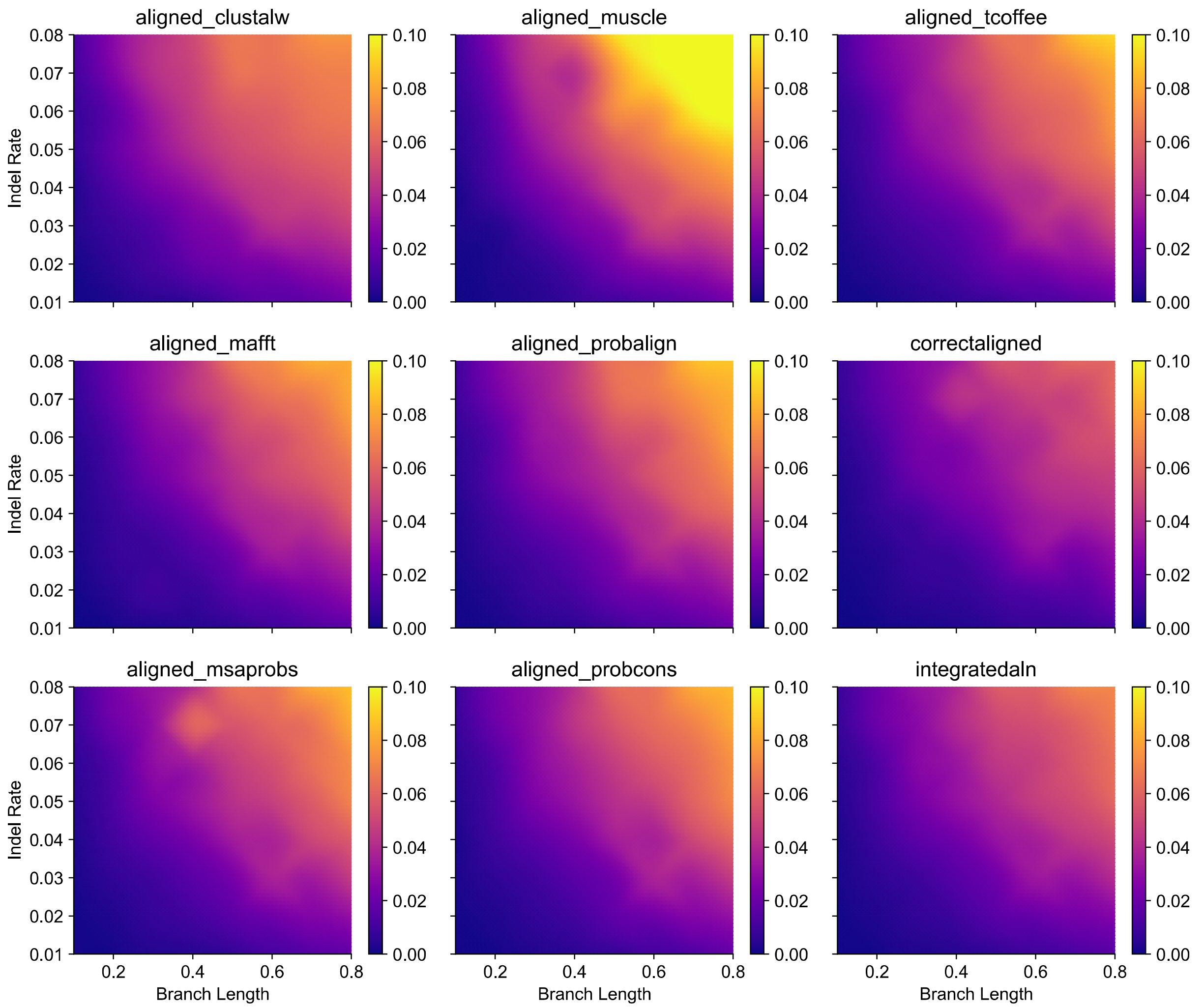


**Figure S16. Alignment-integrated ancestral sequence reconstruction reduces deletion ASR error rates in 3-taxon simulations across a variety of conditions.** We simulated protein sequences along a 3-taxon phylogeny with equal branch lengths and the same insertion-deletion (indel) rate across the phylogeny (see Methods, Supplementary Fig. S12). The resulting extant sequence data was aligned using 7 different sequence-alignment methods (see Methods). We used each alignment to infer the most likely ancestral sequence at the single node on the 3-taxon phylogeny. Additionally, alignment-integrated ancestral sequences were generated by combining inferences from the 7 sequence-alignment methods (see Methods). In each case, we compared the inferred ancestral sequence to the correct simulated sequence to estimate ancestral sequence reconstruction (ASR) error rates (see Methods). ASR errors were divided into 4 categories: 1) residue errors, in which both correct and inferred ancestral sequences have a residue at a given position in the alignment, but the residues differ; 2) insertion errors, in which the inferred sequence has a residue at a given alignment position, but the correct sequence has a gap state; 3) deletion errors, in which the inferred sequence has a gap, but the correct sequence has a residue, and 4) total errors, which combine residue- insertion- and deletion-errors. Sequence-wide error rates (errors/site) were computed by dividing the number of errors by the length of the pairwise alignment of the inferred and correct ancestral sequences. For each combination of branch-length and indel-rate, we plot the average rate of deletion ASR errors over 100 replicate data sets for each alignment method, with purple indicating low ASR error rates, and yellow indicating high rates of ASR errors.
